# RNAMaRs: an interpretable framework for inferring multivalent RNA Motifs and cognate Regulators of Splicing

**DOI:** 10.64898/2026.01.31.703040

**Authors:** Mariachiara Grieco, Tommaso Becchi, Gabriele Boscagli, Francesca Priante, Livia Caizzi, Uberto Pozzoli, Matteo Cereda

## Abstract

Alternative splicing expands proteomic diversity and is shaped by interactions between RNA-binding proteins (RBPs) and multivalent RNA motifs. Linking sequence elements to regulatory proteins remains difficult from sequence information alone. Here we present RNAMaRs, a interpretable statistical framework that combines motif discovery with *in vivo* binding and splicing responses to infer motif-RBP relationships. RNAMaRs learns RBP binding principles, weights signal quality, and optimizes motif discovery in an RBP-specific manner. Across ENCODE datasets RNAMaRs consistently prioritizes the perturbed regulator, especially for large splicing effects. Independent validation in prostate cancer cells recapitulates HNRNPK binding signatures, supporting transferability across an unseen cellular context.

## Background

Alternative splicing expands proteome diversity by selecting different exon combinations in mature transcripts. Catalyzed by core spliceosomal factors, alternative splicing events (ASEs) are finely controlled by auxiliary RNA-binding proteins (RBPs), which recognize clusters of short sequence motifs, known as multivalent RNA motifs (MRMs) [1]. Being multivalent, these motifs can be bound by multiple RBPs and their position relative to regulated exons helps determine distinct splicing outcomes. Notably, MRMs are enriched in sequences encoding intrinsically disordered regions, with GA-rich mRNAs pivotal to maintain collective balance of condensation-prone proteins (*i.e.,* interstasis) [2].

Disruption of RBP-RNA interactions is a recurring feature of human diseases and it is increasingly being explored as a therapeutic entry point [3–5]. A central challenge is to identify which RBPs bind which motifs in a cellular context, and which of those relationships are functionally coupled with aberrant splicing. Addressing this gap is necessary to derive mechanistic rules of the splicing code and identify novel actionable regulatory targets.

In this light, *in vivo* cross-linking and immunoprecipitation experiments (CLIP) assays have been exploited to determine individual RBP-mRNA interactions [6]. ENCODE has further leveraged enhanced CLIP (eCLIP) to profile splicing-associated regulatory principles for ∼150 RBPs across two cell lines [7], providing an important resource but still covering only a fraction of known RBPs and physiological conditions. Critically, however, the splicing principles distilled from these datasets are typically formulated in a protein-centric framework, even though most splicing outcomes are governed by combinatorial regulation among multiple RBPs.

Computational motif discovery has progressed from position-specific scoring matrices from primary sequences [8] to frameworks that capture dependencies, degeneracy, gapped k-mers, and co-occurring regulators, as well as approaches that incorporate RNA secondary structure [9–15]. Deep learning further improves predictive performance but often reduces interpretability, making it difficult to translate predictions into testable mechanisms [16–20], However, a key unresolved challenge remains linking MRMs to their cognate RBPs from primary sequence alone.

To address this challenge, we introduce RNAMaRs, a highly interpretable algorithm designed to identify MRMs and link them to cognate RBPs from primary sequence. Building on RNAmotifs [1], RNAMaRs integrates eCLIP- and RNA-seq data to establish expectations of positional regulatory behaviour of RBPs, while incorporating robustness-aware metrics to improve MRM detection. Applied to ENCODE knockdown datasets, RNAMaRs prioritizes the binding principles of the perturbed protein, with its performance scaling with the magnitude of splicing changes. Our framework further provides integrated summary visualization that jointly reports MRM enrichments, regional binding preferences, and ranked candidate regulatory proteins. Finally, applied to RNA-seq data upon HNRNPK depletion in prostate cancer cells, RNAMaRs robustly prioritized HNRNPK-associated positional and sequence signatures consistent with its known specificity, supporting the applicability of the framework across cell types and datasets.

## Results

### Overview of the RNAMaRs framework

RNAMaRs employs the RNAmotifs algorithm to identify clusters of short degenerate four-nucleotide motifs, namely multivalent RNA motifs (MRMs) that dictate RNA-binding protein (RBP) binding preferences [1]. RNAMaRs then links MRMs with their cognate RBPs and prioritize putative splicing regulators directly from RNA-seq-derived exon inclusion changes (Fig.1). This prioritization integrates complementary evidence from ENCODE resources ([7,21] by combining eCLIP-derived RBP-RNA interaction profiles (physical association) with RNA-seq defined RBP-regulated exons (functional evidence). Together, these data provide an empirical basis for ranking proteins whose positional binding profiles (*i.e.*, splicing maps) most closely recapitulate MRM-specific maps. This framework supports inference of cognate MRM-RBP relationships from shared binding principles while weighting associations by the strength and consistency of the underlying evidence.

**Figure 1.**
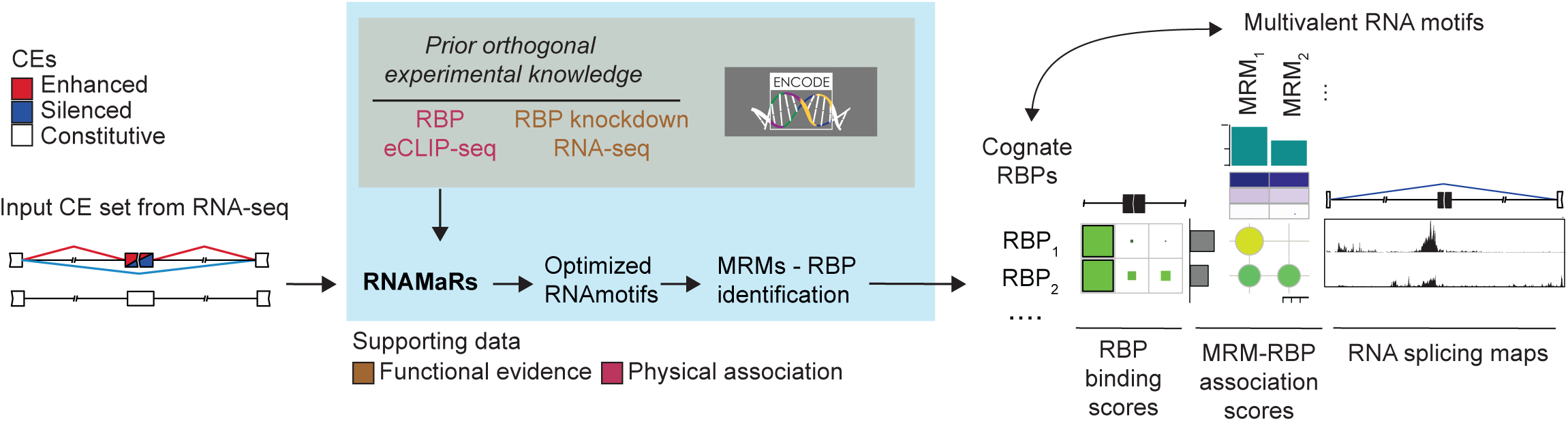
Overview of the RNAMaRs framework for identifying splicing regulatory RBPs from RNA-seq. RNAMaRs takes as input a set of cassette exons (CEs) derived from RNA-seq and annotated as enhanced, silenced, or constitutive. The framework leverages prior orthogonal experimental knowledge from ENCODE, including eCLIP-seq binding profiles (physical association) and RNA-seq following RBP knockdown (functional evidence), together with optimized RNAmotifs settings to identify multivalent RNA motifs (MRMs). RNAMaRs links MRMs to cognate RBPs by comparing MRM-specific and RBP-specific RNA splicing maps, producing MRM-RBP association scores alongside region-specific RBP binding scores. Outputs include ranked candidate RBPs, enriched MRMs, and RNA splicing maps that summarize positional binding and regulatory signatures.

RNAMaRs takes as input a set of cassette exons (CE) annotated with differential inclusion class (enhanced or silenced), together with a reference cell line for eCLIP data (HepG2 or K562). The algorithms runs RNAmotifs using RBP-optimized parameters to identify MRMs enriched around the input exons, then computes motif-protein association scores (ASs) by comparing motif-derived splicing maps to eCLIP-derived binding maps (Methods and Additional file 1: Fig. S1). To limit the impact of noisy binding signals, ASs are down-weighted using binding-robustness measures. The primary outputs are two AS matrices (*i.e.*, one for enhanced and one for silenced exons), along with summary visualizations that combine motif enrichment, regional binding preferences, and a ranked list of candidate regulators.

In the following sections, we describe the workflow in three parts: (*i*) ENCODE data retrieval and preprocessing to quantify RBP splice-proximal binding, (*ii*) RNAmotifs parameter optimization by grid search, and (*iii*) application of RNAMaRs to user-defined exons to infer candidate *trans*-acting regulators from *cis*-acting motif signatures.

### Multi-layer evidence integration for MRM-RBP associations

To reliably associate MRMs with their cognate RBPs, we first quantified how reproducibly RBP binding patterns could be detected using available eCLIP and RNA-seq data. As eCLIP signal strength and consistency vary across proteins and datasets [6], we benchmarked the detectability of position-dependent binding principles prior to motif-protein association. We retrieved ENCODE datasets comprising (*i*) cassette exons (CEs) showing differential inclusion upon RBP depletion and (*ii*) eCLIP cross-linking sites in HepG2 and K562 cells [21] (Methods). We retained only RBPs for which both data types were available, yielding 70 and 80 RBPs in HepG2 and K562, respectively (Fig.2A). We further required the availability of an mCross position weight matrix (PWM) [11] and excluded RBPs without an associated PWM, thereby restricting downstream analyses to proteins with validated sequence-specific binding models (Additional file 2: Table S1). Finally, because at least 50 exons are required for robust inference of positional binding patterns [22], we retained only RBPs with at least 50 regulated CEs (Fig.2B and Additional file 1: Fig. S2A, Additional file 2: Table S2). This filtering resulted in a final set of 21 RBPs, including U2AF1, U2AF2, PTBP1, SRSF1, PRPF8, HNRNPU and SF3B4 in both cell lines.

**Figure 2.**
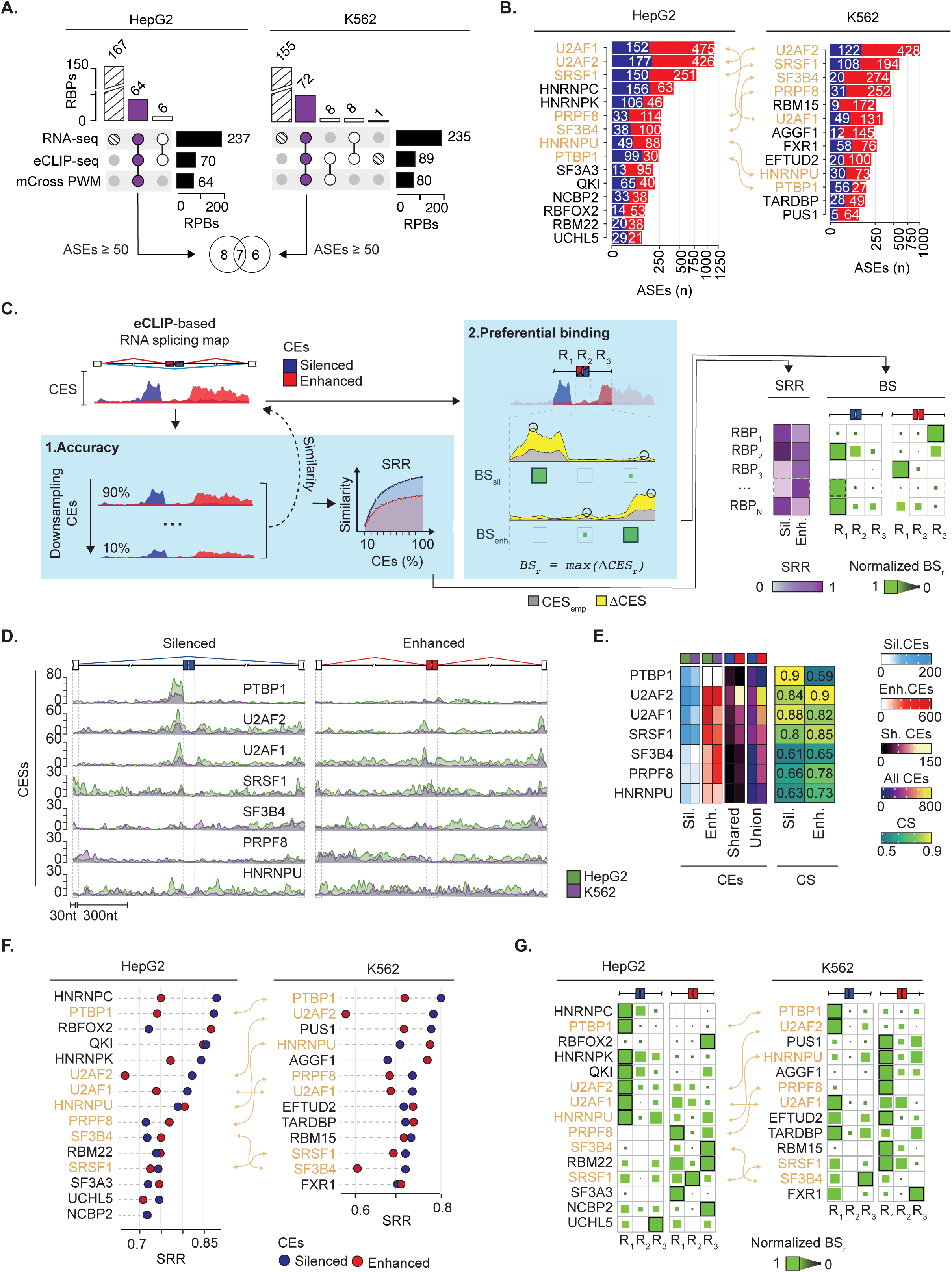
Robustness and positional specificity of eCLIP-derived RBP binding profiles. **A.** Overview of supporting datasets used for RBP selection in HepG2 and K562 cells, including RNA-seq-based splicing changes following RBP depletion (functional evidence), eCLIP-seq binding data (physical association), and available mCross PWMs (intrinsic specificity). Only RBPs with ≥50 regulated cassette exons (CEs) and available PWMs were retained. **B.** Number of alternative splicing events (ASEs) per RBP in HepG2 and K562 cells. Orange arrows connect RBPs with available data in HepG2 and K562 cell lines. **C.** Schematic of the analysis framework used to assess accuracy and binding principles of RBP from eCLIP data. Binding accuracy was evaluated by downsampling exons and computing the signal recovery rate (SRR), while preferential binding was quantified across three splice-proximal regions (R1-R3) using normalized binding score (BS). Circles refer to maximum CESs. Green squares with black contour refer to maximum CESs. CES = cross-linking enrichment score. **D.** eCLIP-based RNA splicing maps for RBPs with available data in both cell lines, shown separately for silenced and enhanced exons, illustrating position-dependent cross-linking enrichment scores (CESs). **E.** Heatmap of cosine similarity of RNA splicing maps between cell lines. Reported are the number of silenced, enhanced, shared, or combined exon sets for HepG2 and K562. **F.** Distribution of SRR values across RBPs in HepG2 and K562, highlighting proteins with the most robust binding profiles. Orange arrows connect RBPs with available data in HepG2 and K562 cell lines. **G.** Heatmaps of normalized BS values across regulatory regions (R1-R3) for silenced and enhanced exons, summarizing RBP-specific positional binding preferences.

We next established an eCLIP splicing-map benchmarking framework to evaluate map robustness and positional regulatory principles (Fig. 2C). For each RBP, we generated RNA splicing maps by quantifying the enrichment of eCLIP cross-linking sites at each nucleotide position around enhanced or silenced CEs relative to constitutive exons, defining a cross-linking enrichment score (CES, Methods and Additional file 1: Fig. S2B-C). CES provides a position-resolved measure of preferential binding along the splicing map. First, to assess robustness, we performed downsampling simulations in which CES profiles were recomputed using increasing fractions of regulated exons (Fig.2C, left panel, and Additional file 1: Fig. S2C). We quantified agreement between each downsampled map and the corresponding full map using cosine similarity (CS) and summarized recovery across sampling fractions using the Signal Recovery Rate (SRR), defined as the area under the similarity curve. SRR therefore captures how reliably an eCLIP-derived positional profile can be reconstructed from the available events. Second, to quantify binding principles, we evaluated enrichment within three canonical regions flanking splice junctions [1] against an empirical background obtained by permuting exon labels (Fig.2C, central panel, Methods). We classified a position significantly bound when its CES exceeded the 95^th^ percentile of the background of the empirical null distribution. We then summarized positional binding propensity as the difference between the observed CES profile and the empirical background (ΔCES). Finally, we condensed binding propensity of each region into a Binding Score (BS), defined as the maximum ΔCES within that region, and rescaled to allow comparisons across RBPs.

Before interpreting regional preferences, we assessed reproducibility across cellular context by comparing CES profiles between HepG2 and K562 for common RBPs (Fig.2D). Overall, splicing maps were broadly concordant, with a mean CS of 0.76 across enhanced and silenced exon classes (Fig.2E). Concordance increased with the number of CEs used to build each map (*i.e.,* PTBP1, U2AF1/2, and SRSF1), consistent with improved stability at larger sample sizes.

Having established map reproducibility, we applied the SRR-BS framework (Fig.2C) to quantify map robustness (SRR) and summarize positional binding preferences (BS) across RBPs and exon classes within each cell line. SRR values were generally high (mean = 0.78, Fig.2F), with the strongest recovery observed for HNRNPC at silenced exons in HepG2 (SRR = 0.92), consistent with a robust positional signal. In HepG2, HNRNPC, PTBP1, QKI and U2AF2 showed higher SRR for silenced than for enhanced CEs, whereas RBFOX2 showed higher SRR for enhanced events. In K562, U2AF2 and SF3B4 similarly exhibited higher SRR for silenced exons. By contrast, RBPs such as HNRNPU, SRSF1, and TARDBP displayed comparable SRR across exon classes, indicating similar robustness for enhanced and silenced maps.

We then compared BS profiles across RBPs to summarize region-specific binding preferences. In HepG2, BS recapitulated established splicing regulatory architectures [7,23]. Specifically, HNRNPC, PTBP1 and U2AF1/2 showed strongest upstream binding near silenced exons, SRSF1 peaked within the exonic core of enhanced events, and RBFOX2 preferentially bound downstream of enhanced exons. In K562, BS similarly recovered known preferences, including upstream binding of PTBP1 and U2AF1/2 near silenced exons and PRPF8 enrichment upstream of enhanced exons.

Together, these results support the use of SRR and BS as complementary quantities in the downstream MRM-RBP association step. SRR captures how reliably an eCLIP-derived splicing map can be reconstructed from the available events, whereas BS provides a region-level summary of where binding is concentrated along the splice-proximal landscape. In practice, we use them to temper associations that are driven by weak or poorly reproducible binding profiles, rather than letting those signals dominate motif linking.

### Optimization of RNAmotifs to capture RBP-specific MRMs

Building on the SRR and BS analyses, which established robust and interpretable splice-proximal binding principles, we next optimized RNAmotifs [1] to better recovery of RBP-specific MRMs, thereby increasing sensitivity for subsequent MRM-RBP association. Because MRM detection depends on motif clustering and enrichment at exon-intron boundaries, we systematically tuned RNAmotifs settings by grid search and selected optimal values using AUROC-based performance (Fig.3A and Additional file 1: Fig. S3). Specifically, we run RNAmotifs across distinct clustering (10, 30, 50, and 70 nts) and enrichment window sizes (30, 50, 100, 200, and 300 nts). For each configuration, we identified enriched MRMs for enhanced and silenced exons and constructed motif-specific RNA splicing maps. We then compared them with eCLIP-derived RBP binding profiles restricted to exons containing each MRM using cosine similarity. Finally, we weighted these values by the SRR to account for differences in binding robustness across RBPs, yielding MRM-RBP ASs (Methods, Fig.3A and Additional file 1: Fig. S3). The ASs range from 0 to 1, where 1 indicates a strong likelihood that the RBP binds to the MRM to modulate splicing.

**Figure 3.**
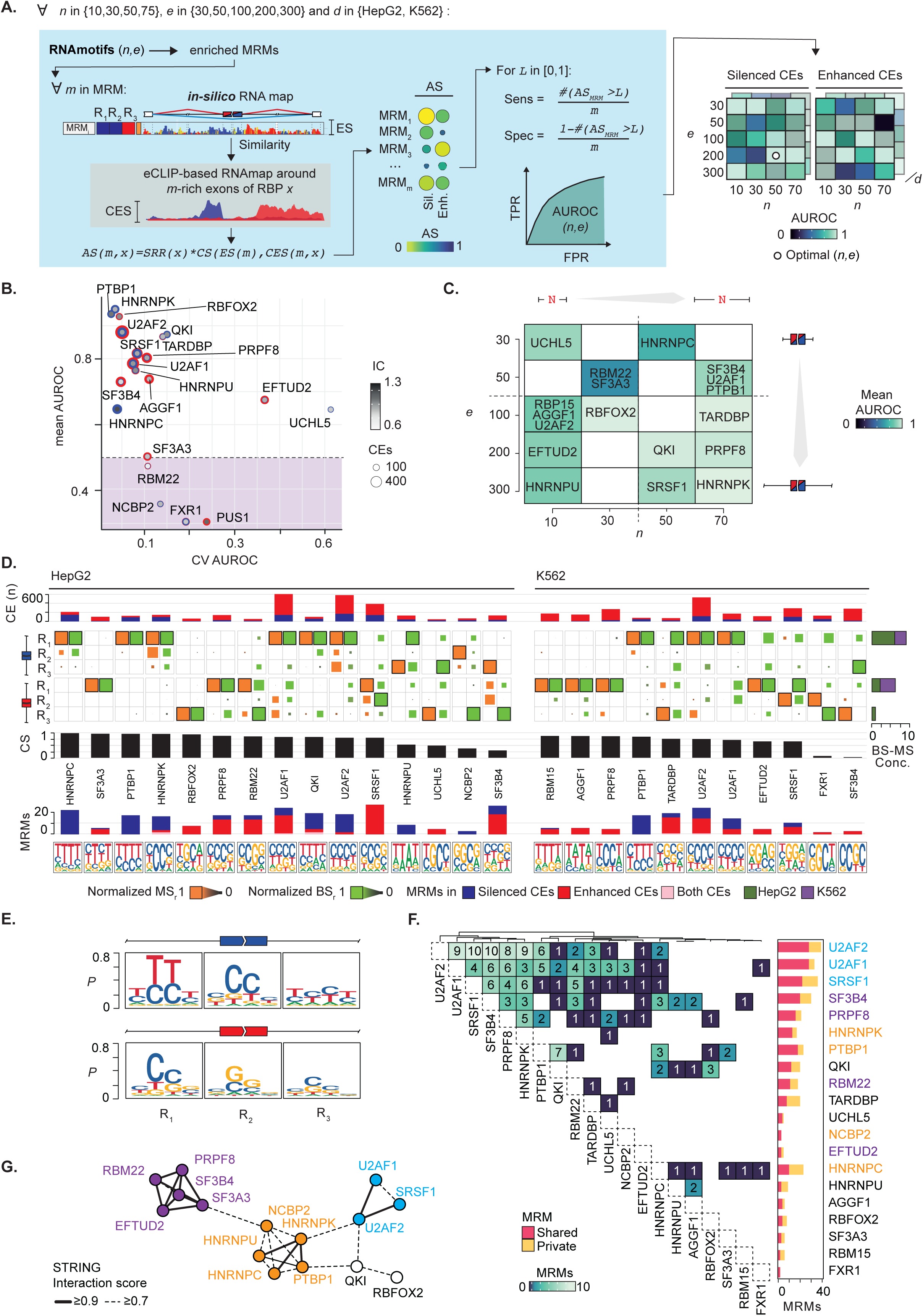
Optimization of RNAmotifs parameters and construction of MRM-RBP association scores. **A.** Schematic of the parameter-tuning strategy. RNAmotifs is run across a grid of clustering (*n*) and enrichment (*e*) window sizes to identify enriched MRMs and derive *in silico* MRM-specific RNA splicing maps. For each MRM and threshold *L* in [0,1], ASs are used to compute sensitivity and specificity, generate ROC curves, and quantify performance by AUROC to select optimal (*n,e*) settings. **B.** Mean AUROC versus AUROC variability (coefficient of variation, CV) across parameter sets for each RBP, highlighting proteins with consistently high performance and those with lower or less stable discrimination. Point size reflects the number of regulated CEs (blue=silenced, red=enhanced), and shading reflects PWM information content (IC). **C.** Heatmap of mean AUROC across the (*n,e*) grid, showing optimal parameters for individual RBPs with best AUROC ≥0.5. **D.** Summary of optimized outputs in HepG2 and K562 for RBPs having ≥1 enriched MRM in the RNAmotifs run with optimized parameters. The top barplot depicts the number of input CEs for each RBP. The central heatmaps report normalized MRM scores (MS) and normalized binding scores (BS) across regulatory regions (R1-R3) for silenced and enhanced exons. Barplots show the cosine similarity (CS) between motif-derived and eCLIP-derived profiles and the number of enriched MRMs, with datasets ordered by decreasing CS. Sequence logos summarised enriched MRMs. Right barplots indicate the number of concordant cases showing maxima MS and BS values in the same region. **E.** Sequence logos summarised enriched MRMs across regions (R1-R3) for enhanced and silenced exons, illustrating region-specific motif localization patterns. **F.** Overlap of enriched MRMs across RBPs, separating shared and RBP-private motifs. Colorcode refers to the main protein-protein interaction summarised in panel H. **G.** STRING protein-protein interaction network of the RBPs analyzed, with edges indicating interaction confidence thresholds.

To evaluate performance, we applied a series of cutoffs to AS values to classify MRMs as associated, or not, with each RBP and constructed receiver operating characteristic (ROC) curves across these cutoffs. We used the area under the ROC curve (AUROC) to select parameter sets maximizing AUROC, corresponding to the RNAmotifs configuration that best captured RBP-specific MRMs. Together, RNAmotifs optimization revealed RBP-specific differences in motif detectability driven mainly by data availability and, to a lesser extent, sequence specificity (Fig.3B, Additional file 1: Fig. S4A, and Additional file 2: Table S3). Most RBPs reached mean AUROC>0.7, with canonical splicing regulators (*i.e.,* PTBP1, HNRNPK, RBFOX2, and U2AF2) among the top performers.

The AUROC landscape further allowed us to relate optimal parameter choices to the known binding behavior of individual RBPs (Fig.3C). Splice-site proximal binder such as U2AF2 achieved maximal AUROC with short clustering regions (∼10nts), while U2AF1 peaked with a small enrichment windows (∼50nts), consistent with their recognition of short sequence elements that cluster over limited distances [7]. In contrast, RBPs known to bind broader or more distributed footprints, such as PTBP1, HNRNPK and QKI, showed optimal performance with larger clustering windows (50-70nts) and intermediate-to-large enrichment windows (50-300 nts), capturing clusters of motifs spread across wider intronic regions [7]. Together, these examples illustrate that optimal RNAmotifs parameters reflect distinct RBP binding architectures.

We next tested whether MRM enrichments from the optimal RNAmotifs runs recovered the expected RBP binding principles. Specifically, we assessed whether: (*i*) enriched MRMs localized to the same splice-proximal regions where eCLIP shows maximal binding for the corresponding RBP, and (*ii*) the inferred motifs compositions reflected the established nucleotide preferences for that protein.

First, to quantify agreement between RNAmotifs and eCLIP-based results, we compared the cumulative motif regional enrichment, namely MRM score (MS), and the corresponding BS using CS for silenced and enhanced exons (Methods, Fig.3D). Eighteen datasets showed high concordance (CS>0.7) indicating that the optimal RNAmotifs runs generally recovered motif enrichments that matched the binding profiles. Specifically, enriched MRMs consistently localize to the same regions showing maximal BS in 16 datasets (57%). This was corroborated by the strong correlation between MS and BS across enrichment regions (Additional file 1: Fig. S4B). The most frequent shared region of maximal MS and BS was upstream 3’ splice site (ss) of silenced exons, observed in nine cases across cell lines (Fig.3D). The relative distribution of MRMs across exon classes further reflected RBP-specific regulatory preferences, with SR and hnRNP proteins enriched for motifs in enhanced and silenced exons, respectively (Fig.3D).

Second, we examined whether datasets showing enrichment in the same splice-proximal regions also shared common sequence preferences. To this end, we derived PWMs from enriched MRMs using their MSs (Methods). Across RBPs, the inferred MRMs recapitulated expected sequence patterns, with PWMs frequently highlighting nucleotide compositions characteristic of known RBP-binding preferences (Fig.3D lower part). In particular, datasets peaking in upstream intronic regions often produced pyrimidine- or C/T-rich PWMs, consistent with the prevalence of low-complexity polypyrimidine-like elements near the 3′ ss that are commonly exploited by splicing factors [24,25]. In line with this, when we aggregated enriched MRMs across all regulatory regions (Methods), we observed that silenced exons were dominated by pyrimidine-rich cores, whereas enhanced exons showed more G/C-rich core dinucleotides (Fig.3E). In both classes, PWMs showed the highest information content (IC) upstream of the 3′ss, indicating that the strongest signal localized to this region.

Finally, to examine RBP sequence specificity, we measured overlap of enriched MRMs across RBPs in the two cell lines (Fig.3F and Additional file 1: Fig. S4C). Several MRMs were shared among functionally related RBPs, such as U2AF2, U2AF1, and SRSF1, while others were RBP-specific, like HNRNPC and TARDBP. To independently evaluate whether shared MRMs reflect shared regulatory modules, we retrieved the RBP protein-protein interaction network from STRING database [26]. RBPs with overlapping MRMs tended to cluster within established interaction modules, whereas proteins with unique MRMs were more peripheral (Fig.3G). Overall, these results indicate that the optimized RNAmotifs parameters recover both common regulatory programs and protein-specific motif signatures.

Collectively, these analyses demonstrate that RNAmotifs optimization, guided by binding robustness and positional enrichment, reliably recovers biologically meaningful MRMs whose localization, sequence composition, and sharing patterns are consistent with known RBP binding architectures. These properties provide a concrete basis for the subsequent MRM-RBP association step, rather than relying on sequence enrichment alone.

### Validation of RNAMaRs using ENCODE RNA-seq perturbation data

To assess the ability of RNAMaRs to recover known splicing regulators, we evaluated the method on differentially spliced CEs regulated by RBPs that we characterized in the previous sections. For each dataset, exons were used as input and RNAMaRs was run in a blinded setting (*i.e.,* without specifying the perturbed RBP). All results are available as Additional file 1: Fig. S7-34.

As an illustrative example, we examined RNAMaRs results for PTBP1-regulated silenced exons (Fig.4A and Additional file 1: Fig. S5A-B). RNAMaRs prioritizes PTBP1 as the top candidate regulator using data of both cell lines, with high ASs driven by C/T-rich MRMs enriched at the polypyrimidine tract [1]. The corresponding RNA splicing maps recapitulate the well-established PTBP1 regulatory architecture [27–29], providing a biologically consistent visualization of the results. U2AF2 and U2AF1 emerged as the second and third top-ranked candidates, respectively, consistent with partial overlap in splice-proximal binding preferences and known competition between PTBP1 and U2AF factors for 3′ss recognition [30]. PTBP1 depletion was also associated with increased U2AF1/2 expression (Fig.4A). Although this change was not statistically significant, it is compatible with the compensatory feedback. Together, these findings illustrate the ability of RNAMaRs to recover the dominant regulator while also highlighting closely related RBPs within a shared regulatory landscape.

**Figure 4.**
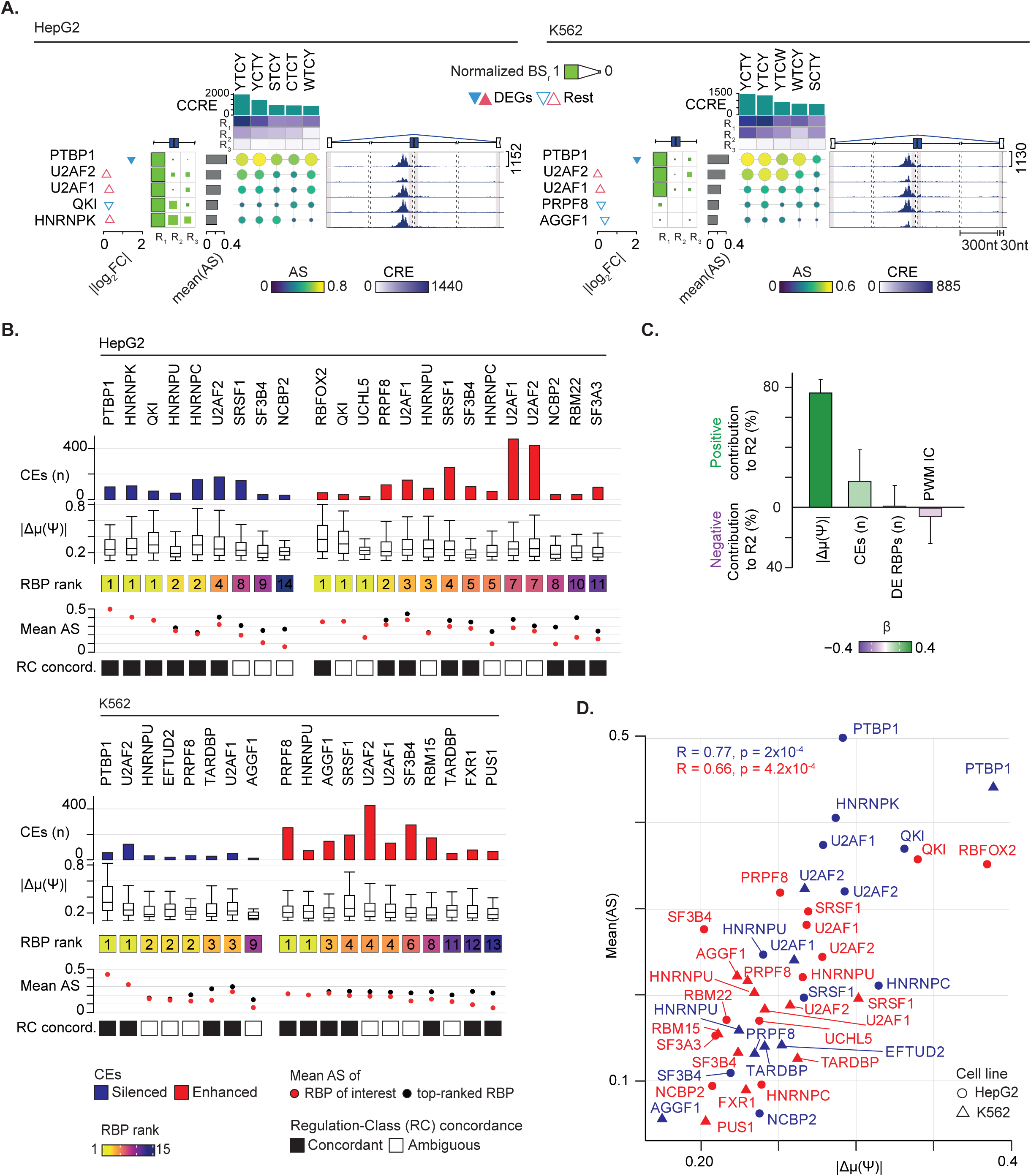
Validation of RNAMaRs on ENCODE RBP perturbation datasets. **A.** Representative RNAMaRs output for PTBP1-regulated silenced exons in HepG2 (left) and K562 (right). The visualization integrates normalized BSs across splice-proximal regions, MRM-RBP ASs, cumulative combined regional enrichment (CCRE) across enriched MRMs, and the corresponding RNA splicing maps. Differential expression upon knockdown is shown as triangles. The upper and lower vertices indicated up- and down-regulation, respectively, and filled triangles denoted differential expression changes (*i.e*., |log2FC| ≥0.1 and adjusted p-values ≤0.1). Barplots indicate the mean AS per candidate regulator. **B.** Summary of RNAMaRs performance across all evaluated RBPs and both cell lines. For each dataset, the figure reports the cell line, exon class (enhanced or silenced), regulation-class (RC) concordance category, number of regulated CEs, |Δμ(Ψ)| distribution, the rank of the true regulatory RBP, and mean AS (dots denote the RBP of interest and the top-ranked RBP). Only datasets with at least one enriched MRM are shown. **C.** Multivariate analysis relating prediction performance (mean AS) to technical and biological features, including mean |Δμ(Ψ)|, number of input CEs, mCrossDB PWM information content (IC), and the number of differentially expressed (DE) RBPs. Bars indicate relative importance to explained variance (R²). **D.** Relationship between mean |Δμ(Ψ)| and mean AS across datasets, with points labeled by RBP and colored by regulation class. Pearson’s correlation values stratified by regulation classes are reported.

We summarized RNAMaRs performance across all evaluated RBPs in HepG2 and K562 cells in Fig.4B. For each dataset, we reported the rank of the true regulatory RBP together with the number of regulated exons, the distribution of absolute inclusion changes (|Δμ(Ψ|), and the mean AS. We restricted evaluation to datasets with at least one enriched MRM, resulting in 42 evaluable datasets (75%). RNAMaRs ranked the true regulator first in ten cases (24%) and placed it within the top five in 28 cases (67%). Among these top-five cases, 19 (68%) showed regulation-class concordance, defined as agreement between the predicted regulation class (enhanced or silenced) and the exon class in which the RBP showed its strongest BS.

To investigate how biological and technical features influenced prioritization, we modelled RNAMaRs performance using a multivariable covariance approach and quantified the relative contribution of each feature as previously proposed [31] (Methods). Specifically, we used a generalized linear regression fitting the AS as a function of |Δμ(Ψ)|, number of CEs, sequence specificity (PWM Information Content), and the number of differentially expressed RBPs upon protein depletion. The magnitude of splicing changes (|Δμ(Ψ)|) emerged as the strongest positive predictor of RNAMaRs performance, followed by the number of input CEs, whereas the remaining features showed marginal non-significant contributions (Fig.4C). Consistently, after stratification by exon regulation classes, we observed a significant positive correlation between |Δμ(Ψ)| and the mean AS across datasets (Fig.4D), indicating that stronger splicing perturbations yield higher-confidence RNAMaRs predictions across both cell lines.

Overall, RNAMaRs prioritizes known splicing regulators, most reliably in datasets with stronger inclusion changes and when the inferred regulation aligns with canonical RBP binding principles. Consistently, performance increases with the magnitude of the underlying regulatory signal.

### Validation of RNAMaRs on RNA-seq from HNRNPK depletion in PC3 cells

To further validate RNAMaRs on an independent dataset, we focused on HNRNPK, which showed robust RNAMaRs performance in our ENCODE benchmarks and has been implicated as splicing regulator in prostate cancer [31]. We generated RNA-seq data following HNRNPK depletion in PC3 cells with samples segregated by conditions as expected (Additional file 1: Fig. S6A). We identified 4,641 differentially spliced CEs and 3,601 constitutive exons (Fig.5A), which were used as input for RNAMaRs to identify the candidate regulator and associated MRMs. Because eCLIP data for HNRNPK in K562 did not meet the minimum exon requirements (Additional file 1: Fig. S2A), we used HepG2 eCLIP data as the binding reference for RNAMaRs. Notably, HNRNPK-regulated exons in PC3 showed a limited overlap with those in HepG2 (7% and 3% for silenced and enhanced exons, respectively; Additional file 1: Fig. S6B), providing a stringent test of RNAMaRs robustness across largely non-overlapping exon sets.

**Figure 5.**
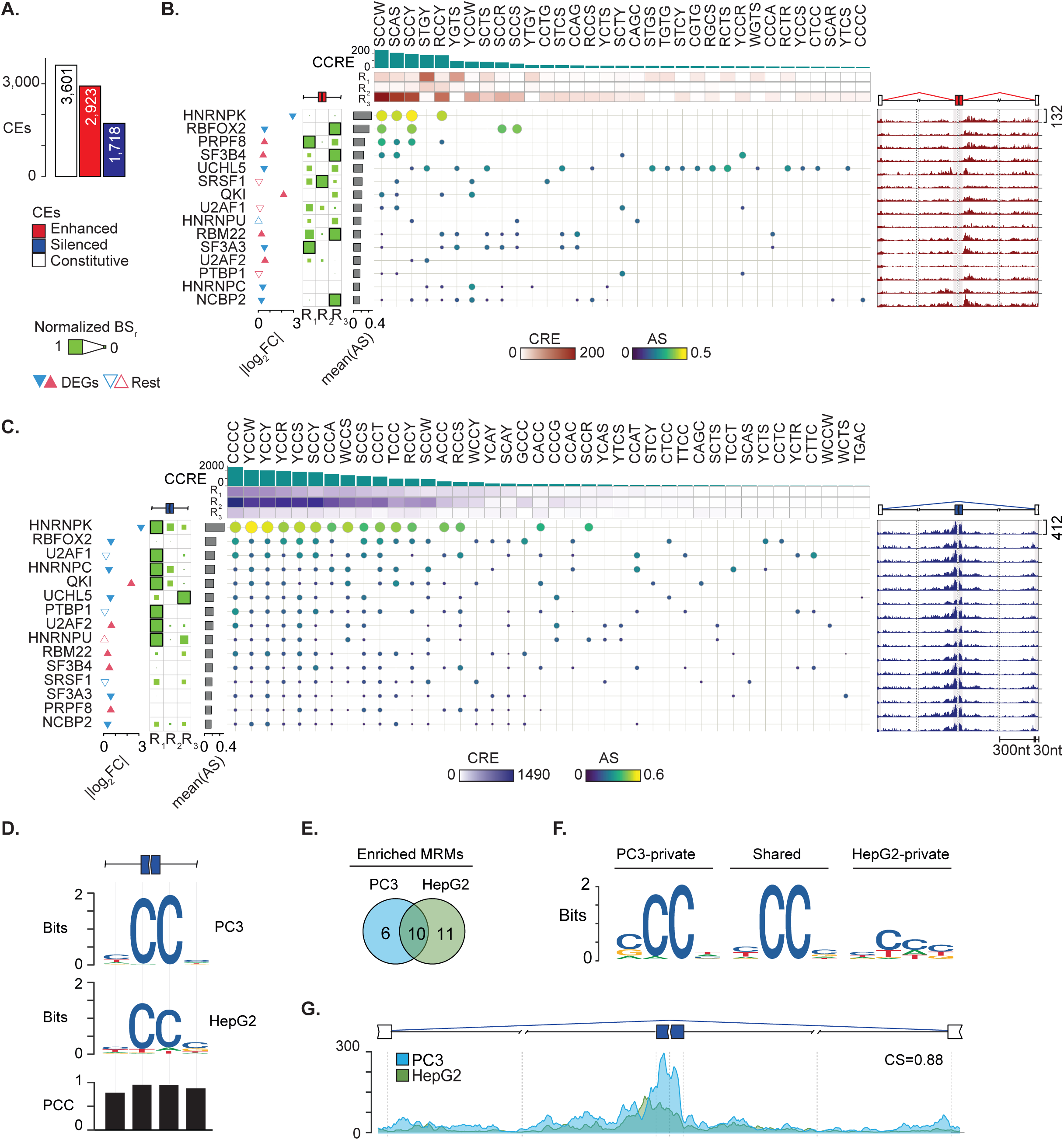
RNAMaRs identifies HNRNPK as a splicing regulator in HNRNPK-depleted PC3 cells. **A.** Distribution of differentially spliced CEs classified as constitutive, enhanced, or silenced following HNRNPK depletion in PC3 cells. **B-C.** RNAMaRs summary visualizations for enhanced (B) and silenced (C) exons, showing region-specific normalized binding scores (BSs), association scores (ASs) between MRMs and candidate RBPs, and RNA splicing maps. **D.** Sequence logos summarizing enriched MRMs in silenced exons associated with HNRNPK across datasets. Barplot reports positional similarities between the two PWMs measured as Pearson Correlation Coefficients (PCC). **E.** Venn diagram representing the intersection of MRMs enriched in silenced exons between PC3 and HepG2 cell lines. **F.** Sequence logos summarizing MRMs shared and private between cell lines. **G.** MRM-specific RNA splicing maps for silenced exons in the two cell lines.

RNAMaRs accurately prioritized HNRNPK-MRM associations for both silenced and enhanced exons (Fig.5B-C), driven by CC-rich motifs enriched across splice-proximal regions. These motifs were preferentially associated with HNRNPK relative to other RBPs, consistent with its known affinity for CC-rich sequences [7,11,32]. Enrichment was most pronounced for silenced exons, where MRMs localized mainly within the exon body and upstream the 3’ss, consistent with regions exhibiting the highest BSs (Fig.5C). RNAMaRs further ranked RBFOX2 as the second candidate regulator, particularly for enhanced exons, in line with its established role in promoting exon inclusion [7]. RBFOX2 was also downregulated upon HNRNPK depletion, consistent with reported interactions between RBFOX2 and HNRNPK [32].

We next compared association outputs between PC3 and HepG2 by testing AS concordance and quantifying MRM similarity between cell lines (Methods). Overall, we observed that MRMs were significantly positively correlated (PCC=0.63), with HNRNPK-enriched MRMs highly concordant (Additional file 1: Fig. S6C). PWMs derived from MRMs enriched in silenced exons were highly correlated between datasets (mean PCC=0.95, Fig.5D), suggesting that HNRNPK binding principles are more conserved for exon repression than for inclusion. We then divided enriched MRMs into shared versus cell line specific sets and collapsed each set into a representative PWM (Fig.5E-F). The shared set showed the greatest conservation, consistent with a core motif recovered robustly by RNAMaRs (Fig.5F). Stratification by enrichment regions yielded comparable results (Additional file 1: Fig. S6D), supporting an HNRNPK regulatory program that remains conserved in R1 and R2.

Finally, we evaluated positional consistency by comparing MRM-specific RNA splicing maps with eCLIP-derived HNRNPK binding profiles for silenced exons. Motif-based and binding-based profiles showed high CS between datasets (CS=0.88), indicating strong agreement in the shape and localization of binding signals (Fig.5G).

Despite minimal overlap between input exon sets, RNAMaRs consistently prioritized HNRNPK and recovered its canonical binding and positional preferences. Together, these results indicate the applicability of RNAMaRs across cell types and independent datasets, extending its use beyond the ENCODE reference context.

## Discussion

Understanding how multivalent *cis*-acting motifs are interpreted by RBPs to regulate alternative splicing remains a central challenge in gene regulation. While large-scale CLIP and RNA-seq perturbation studies have substantially expanded our knowledge of RBP-RNA interactions, translating these data into mechanistic and generalizable rules has been constrained by experimental noise, context dependence, and limited interpretability. Here we introduce RNAMaRs, a computational framework that addresses these challenges by integrating motif discovery with *in vivo* binding and functional splicing evidence to infer interpretable relationships between MRMs and RBPs.

A central advance of RNAMaRs is its explicit integration of physical binding with functional outcome. By jointly leveraging eCLIP-derived binding profiles and RNA-seq splicing perturbations from ENCODE, RNAMaRs establishes an empirical expectation on splice-proximal binding regulation and then attempt to associate motifs with proteins. This differs with conventional motif discovery pipelines, which primarily rely on sequence enrichment and often emphasize predictive performance without directly anchoring motifs to regulatory mechanisms [33]. To further improve reliability, RNAMaRs introduce robustness-aware metrics, including SRR and BS, to down-weight weak or poorly reproducible eCLIP signals and to prioritize RBPs with consistent binding architectures. As a result, inferred associations are statistically supported and align with plausible mechanistic interpretations.

RNAMaRs also implements RBP-specific optimization of RNAmotifs parameters [1]. Rather than applying a single set of motif discovery parameters across diverse RBPs, RNAMaRs systematically tunes clustering and enrichment windows of MRMs using AUROC-based performance against known RBP binding profiles. This strategy reveals that optimal motif-detection scales differ substantially across RBPs, consistent with distinct regulatory modes ranging from splice-proximal recognition of short elements to broader intronic binding landscapes [7]. The concordance observed between optimized motif enrichment profiles and independent eCLIP-derived binding maps demonstrates that parameter tuning is not only a technical improvement but it captures biologically meaningful differences in how RBP engage their targets.

Beyond prioritizing individual regulators, RNAMaRs provides a framework to interpret shared and specific regulatory architectures among RBPs. The observation that functionally related proteins, such as U2AF1, U2AF2 and SRSF1, share subsets of MRMs, while others display highly specific motif repertoires, supports a model in which alternative splicing regulation emerges from partially overlapping multivalent elements. The concordance between shared MRMs and known protein-protein interaction modules further suggest that RNAMaRs can reveal aspects of higher-order RNA regulatory networks, including potentially cooperative or competitive relationships among RBPs that are based on multivalency [2,34].

Our benchmarks across ENCODE perturbation datasets and an independent RNA-seq experiment in prostate cancer cells support the robustness of RNAMaRs and its applicability across cellular context. We observed that prioritization accuracy is sensitive to the magnitude of the underlying splicing perturbation, with stronger splicing changes generally yielding higher-confidence results. Notably, accurate recovery of HNRNPK despite minimal overlap between liver- and prostate-cancer specific exon sets (*i.e.*, HepG2 and PC3) indicates that RNAMaRs can capture transferable binding principles rather than dataset-dependent signatures. The generally stronger performance observed for silenced exons compared to enhanced exons is also consistent with prior observations that repressive splicing architectures are more conserved [35], and thus easier to infer from sequence alone.

Despite these strengths, several limitations should be considered. First, RNAMaRs depends on available eCLIP datasets and it is therefore currently limited to RBPs and cellular contexts profiled with high quality by ENCODE. While the framework is extensible to future datasets, incomplete coverage remains a constraint. Second, the current implementation focuses on splice-proximal regions and does not explicitly model deep long-range interactions, RNA secondary structure, or dynamic changes in RBP concentration and competition. Incorporating structural features and context-dependent RBP expression could therefore refine MRM-RBP associations. Third, extending the framework beyond cassette exons, for example to intron retention and other alternative splicing events, will be important to assess its generality. Finally, although RNAMaRs improves interpretability relative to learning approaches, it remains an inference framework and does not directly establish causality, thus predicted interactions should be prioritized for targeted experimental validation.

## Conclusions

We present RNAMaRs, a publicly available framework that links multivalent *cis*-regulatory RNA motifs to candidate *trans*-acting RBPs by jointly integrating sequence enrichment, *in vivo* binding profiles, and splicing responses to perturbation in an interpretable statistical model. By explicitly conditioning motif discovery on binding patterns that are consistent with regulation and optimizing motif-detection parameters on a protein basis, RNAMaRs captures biologically meaningful differences in splicing regulatory principles, moving beyond sequence-based motif discovery toward mechanistically grounded hypotheses. Across ENCODE perturbation datasets, RNAMaRs consistently prioritized RBPs associated with exon repression and activation, with confidence tracking the magnitude of the underlying splicing effect and particularly strong performance for repressive architectures. An independent HNRNPK knockdown in PC3 cells further supported the transferability of inferred motif-protein relationships across cellular contexts despite minimal overlap between exon sets. Together, RNAMaRs enables the transition from catalogues of motifs and binding maps to interpretable regulatory models, and offers a general strategy for decoding RNA regulatory networks shaped by multivalency.

## Methods

### RNAMaRs workflow

RNAMaRs requires: (*i*) CE coordinates annotated with differential inclusion status (1, enhanced; −1, silenced; 0, constitutive) and (*ii*) the cell line used to source eCLIP peak data (K562 or HepG2, Additional file 1: Fig. S1). Optionally, a differential gene expression table in DESeq2 format [36] could be provided for visualization purposes.

The RNAMaRs workflow comprised three steps. First, as the regulatory RBP was unknown *a priori*, RNAmotifs was run on the input exon set using all combinations of previously optimized parameters as described below. Second, for each enriched MRM, (*i*) the ASs with all RBPs and (*ii*) the RNA splicing maps derived from the RNAmotifs enrichment scores (ES) were computed for enhanced and silenced exons as described below. If an MRM was not enriched under the optimal parameters for a given RBP, the corresponding AS was set to zero. Enhanced and silenced ASs were assembled into two matrices with RBPs as rows and MRMs as columns and summarized together with MSs and MRM-specific splicing maps in a unified visualization. In particular, AS values were shown as a dot heatmap, with dot size and color proportional to AS. For each MRM, regional enrichments across RNAmotifs runs were aggregated using the Fisher’s method, namely combined regional enrichment (CRE), and displayed as a top heatmap annotation. CRE values were then summed across regions for each MRM, termed as cumulative CRE (CCRE), and visualized as a top bar plot (top panel, Additional file 1: Fig. S1). RBP-specific BSs were displayed as separate left heatmaps for enhanced and silenced exons. Finally, mean AS values across MRMs were computed for each RBP, used to rank proteins, and displayed as a left annotation bar plot. Third, for each RBP, RNAmotifs-based RNA splicing maps were generated on subsets of input exons that contained instances of corresponding enriched MRMs, as described below, and displayed these maps on the right for enhanced and silenced exons.

When differential gene expression results were provided, absolute log2 fold changes (|log2FC|) were visualized as triangles on the left (Additional file 1: Fig. S1). The upper and lower vertices indicated up- and down-regulation, respectively, and filled triangles denoted differential expression changes (*i.e*., |log2FC| ≥0.1 and adjusted p-values ≤0.1). All heatmaps were generated using the R package ‘ComplexHeatmap’ v2.10.0 [37].

### Selection of RBP-regulated cassette exons

The catalogue of alternatively spliced CEs upon RBP depletion in HepG2 and K562 cell lines were obtained from [7] (Additional file 2: Table S4). Similarly, RBP cross-linking sites, as iCount peak instances, from eCLIP experiments in HepG2 and K562 cells were collected [24]. Since at least 50 exons are required to disentangle RBPs regulatory patterns [22], only RBPs with at least 50 regulated CEs were retained for downstream analyses (Additional file 1: Fig. S2A). To ensure binding specificity, only RBPs with available 11-nt PWMs in the mCross database [11] were retained (Additional file 2: Table S1). For each retained RBP, the PWM was selected to satisfy two criteria: (*i*) the lowest allelic interaction p-value, indicating the strongest evidence for sequence-dependent binding, and (*ii*) the highest consistency score, reflecting reproducible binding specificity (Additional file 2: Table S5).

Exon inclusion was quantified using the percent spliced-in (Ψ) metric. CEs with (*i*) significant changes in average Ψ changes across replicates upon RBP depletion (*i.e.,* |Δμ(Ψ)| >0.1 and FDR <0.1) and (*ii*) annotated as “cassetteExon” in the UCSC hg19 “knownAlt” table [38] were considered as alternatively spliced and used to benchmark the RNAMaRs framework. Alternatively spliced CEs with at least one iCount peak instance within 300 and 30 nucleotides (nts) into introns and exons, respectively, from the splice sites (ss) (if introns and exon were shorter than 600 and 60 nts, respectively, then the whole introns and exons were evaluated) were considered as RBP-regulated exons and used to optimize RNAmotifs parameters (Additional file 1: Fig. S2B). The sizes of putative regulatory regions were defined according to established guidelines to best capture RBP binding preferences and motif enrichment [7,39,40]. Constitutive exons were defined as previously proposed [41]. Specifically, CEs with (*i*) no significant inclusion changes (*i.e.,* |Δμ(Ψ)| <0.01 and FDR >0.1) upon RBP depletion and (*ii*) annotated in the RefSeq database version 210 [42] but not in the UCSC hg19 “knownAlt” table [38] were considered as constitutively spliced and used as controls.

### Signal recovery rate of eCLIP-derived splicing maps

eCLIP-based RNA splicing maps were generated as previously proposed with minor modifications [31]. Given that RBPs modulate exon inclusion in a position-dependent manner [1], maps were constructed separately for enhanced and silenced CEs (*i.e.,* extending 300 and 30 nts from ss into introns and exons, Additional file 1: Fig. 2C). At each position along the map, a cross-linking enrichment score (CES) was computed comparing the proportion of enhanced (or silenced) and constitutive CEs exhibiting at least one iCount peak using a one-tailed Fisher’s Exact test:

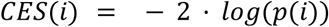

where *p(i)* is the Fisher’s Exact test p-value at *i*-th position along the map. To account for the ability of RBPs to bind clusters of motifs (*i.e.,* MRMs) [1], CES profiles were averaged in a scrolling window of 15 nts using the *filter* function of R package ‘stats’ v4.1.2 and visualized as eCLIP-based RNA splicing map. To evaluate the accuracy of each map in predicting exon inclusion by the corresponding RBP, a bootstrap procedure with downsampling was implemented (Additional file 1: Fig. S2C). For 500 iterations, a percentage *j* of randomly selected CEs ranging from 10% to 100% with 10% increasing step was selected. For each subset of enhanced or silenced exons, a CES profile was calculated as described above. This profile was then compared with the observed one using the cosine similarity (CS), which emphasizes concordance in profile shape rather than absolute signal intensity [43]. For each percentage of exons, CS profiles were averaged across iterations. The Area Under the Curve (AUC), namely Signal Recovery Rate (SRR), was used to interpret the ability of the eCLIP-based RNA splicing map to predict RBP-mediated splicing regulation.

### RBP binding strength from eCLIP data

Impact of RBP regulation on exon inclusion was evaluated across three canonical enrichment regions flanking exon-intron junctions as previously proposed [1]. Regions were defined as follows: R1, upstream intronic sequence extending 300 nts from the 3′ss; R2, exonic sequence spanning 30 nts downstream the 3′ss and 30 nts upstream the 5′ss; and R3, downstream intronic sequence extending 300 nts from the 5′ss. When introns or exons were shorter than 600 nts or 60 nts, respectively, the entire intron and/or exon was evaluated.

An empirical background distribution of CESs along the splicing map (CES*_emp_*) was generated using bootstrap resampling (Additional file 1: Fig. S2D). For each RBP, 1,000 iterations were performed in which exon labels (enhanced/silenced versus constitutive) were randomly shuffled, and CES profiles were computed across the map to form an empirical distribution 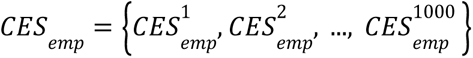 Positions within the enrichment regions (R1-R3) for which the observed CES exceeded the 95^th^ percentile of the CES*_emp_* distribution were classified as significantly bound by the RBP. Positional binding propensity was quantified as the difference between the observed and empirical CES (*ΔCES*). For each region, preferential positional binding was summarized by the maximum *ΔCES*, hereafter termed the binding score (BS). To allow comparisons across different RBPs, the BSs were scaled to the maximum value.

### Identification of RBP-specific MRMs through RNAmotifs

RNAmotifs [1] was employed to identify MRMs of length 4 nts among 256 degenerate and 256 non-degenerate tetramers occurring around CEs. Degenerate tetramers were defined as central dinucleotides (*i.e.,* A,C,G, and T) flanked by two degenerate positions constrained to A/C to G/T substitutions (*i.e.,* R, Y, W, and S). RNAmotifs was adapted to analyze intronic regions of 300 nts. MRMs with region-specific enrichment p-value ≤0.05 (or ≤ the 1^st^ percentile of the p-value distribution when the 1^st^ percentile was ≤0.05) and associated empirical p-value ≤0.0005 were considered as significantly enriched and retained for further analysis. MRMs enriched in enhanced and silenced exons were represented by two separate splicing maps. Specifically, the original RNAmotifs enrichment score (ES) was decomposed into its two regulation-specific components as follows:

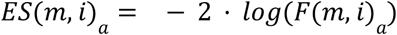

where *m* is the MRM of interest, *i* indexes nucleotide positions along the splicing map, *F* is the RNAmotifs positional enrichment p-value at position *i*, and *a* refers to either enhanced or silenced exons.

To quantify global motif enrichment, RNAmotifs region-specific enrichment p-values of enriched MRMs were combined using Fisher’s Method, hereafter termed regional score (RS), within each splice-proximal regulatory region (R1-R3) for enhanced and silenced exons, respectively, as follows:

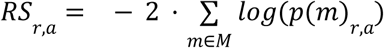

where *r* in {R1,R2,R3}, *a* refers to enhanced or silenced exons, *m* denotes an enriched MRM, and *M* is the set of enriched MRMs. These six RS values were further combined into a single MRM score (MS), which captured overall motif enrichment across R1-R3 regions and regulation types as follows:

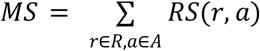

where *A* refers to the set of either enhanced or silenced exons.

RNAmotifs [1] was run with 10,000 bootstrap iterations either (*i*) within the RNAMaRs framework using RBP-specific optimal parameters or (*ii*) during the grid search used to identify these parameters.

### Association between MRMs and RBPs

To associate MRMs with RBPs, RNAmotifs splicing maps were compared with eCLIP-derived RBP splicing maps, separately for enhanced and silenced exons. For each MRM, the subset of input exons containing at least one occurrence of the motif within enriched regions (R1-R3) was identified and an eCLIP-derived RBP splicing map restricted to these exons was constructed. Similarity between MRM- and RBP-related splicing maps was quantified using CS, as follows:

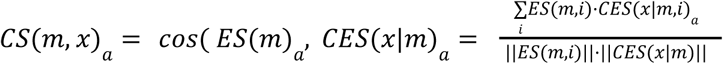

where *m* is the MRM of interest, *x* is the RBP of interest, *a* refers to either enhanced or silenced exons, *i* indexes positions along the map, and ES is the MRM enrichment score provided by RNAmotifs [1].

Finally, to account for differences in binding signal robustness, CSs were weighted by the SRR of the RBP in the AS as follows:

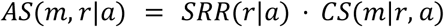

The AS quantified the strength of the relationship between a MRM and an RBP. This procedure was repeated across all RNAmotifs parameter combinations, producing AS matrices used for parameter optimization and downstream RBP prioritization.

### ROC-guided selection of RNAmotifs parameters

For each RBP, optimal RNAmotifs parameters were identified by grid search. Specifically, the size of regions in which motifs were (*i*) clustered with a defined spacing (*i.e.,* the clustering window, *n*) and (*ii*) enriched around exons of interest (*i.e.,* the enrichment window, *e*) were tested for *n*={10, 30, 50, 70} and *e*={30, 50, 100, 200, 300} nts, respectively (Additional file 1: Fig. S3). For each combination of *n* and *e*, RNAmotifs was run as described above. For each MRM, an AS with the RBP of interest was measured as described above. The resulting ASs were scaled using a min-max transformation. To assess how well scores distinguished RBP-associated from RBP-non-associated MRMs, a decision threshold *L* was applied to the normalized ASs. This threshold converted the continuous scores into a binary classification (associated versus not associated), enabling the calculation of sensitivity and specificity. By varying *L* across its full range [0,1] in steps of 0.01, receiver operating characteristic (ROC) curves were generated. For each parameter combination, sensitivity and specificity were computed based on the association of enriched MRMs with the corresponding RBP defined as *AS ≥ L*. Sensitivity was calculated as the number of enriched MRMs associated with the RBP divided by the total number of enriched MRMs. Specificity was calculated as the number of enriched MRMs not associated with the RBP divided by the total number of enriched MRMs. Based on these metrics, the area under the ROC curve (AUROC) was computed for each RBP across cell lines and regulation types, ranking the combinations according to *n* and *e*. The parameter set maximizing the AUROC across cell lines and regulation types was selected as optimal for each RBP. Only RNAmotifs parameter combinations identifying at least 5 enriched MRMs were selected. In addition, when the selected optimal AUROC was <0.5 (*i.e.*, performance worse than random), the default RNAmotifs parameters were used (*n*=30, *e*=30) [1].

### MRM-derived PWM, similarity, and information content evaluation

MRM-specific PWMs were represented as 4×4 positional matrices, with columns corresponding to the four tetramer positions and rows to the four nucleotides. For non-degenerate tetramers, entries were set to 1 for the matching nucleotide at each position and 0 otherwise. For degenerate tetramers, entries corresponding to nucleotides encoded by the degeneracy symbol were set to 0.5, and to 0 for all other nucleotides. A consensus PWM was then obtained by collapsing the tetramer matrices using a weighted average of nucleotide frequencies at each position. Weights were defined as the sum of MS scores across all regulatory regions in which the tetramer was significantly enriched, such that MRMs showing stronger enrichment contributed more to the final consensus. The resulting consensus PWM contained entries representing the weighted frequency of nucleotides at each position across MRMs.

Similarity between MRM-derived PWMs was quantified using the Pearson Correlation Coefficient (PCC) [44]. Given two PWMs, for each of the four positions a PCC was measured between the corresponding frequency vectors, yielding four correlation values (one per position). The mean PCC across position was used as the overall similarity. The Information Content (IC) of a column *i* of a PWM was measured as follows:

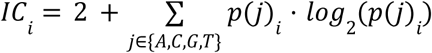

where *p_j_* is the probability of having the letter *j* in the position *i* of the PWM. Probabilities equal to zero were excluded from this calculation, as they do not contribute to the positional information. The overall IC of a PWM was defined as the mean IC across all four positions.

### Multivariable covariance analysis

Relative contributions of biological and technical features to a response variable were quantified as previously proposed [31]. Specifically, the following regressors considered were: |Δμ(Ψ)| levels, number of input exons, PWM ICs, and number of differentially expressed (*i.e*., |log2FC| ≥0.1 and adjusted p-values ≤0.1) RBPs considered in this study (Additional file 2: Table S2). Regressor values were preprocessed using the *preProcess* function from the R package ‘caret’ v6.0-94 [45] with method=c(“center”,“scale”,“YeoJohnson”,“nzv”), which applied near-zero variance filtering, Yeo-Johnson transformation, mean centering, and scaling to unit variance. Datasets with no enriched MRMs were excluded. A generalized linear model (GLM) was then fitted to the response variable using the normalized regressors with the *glm* function in the R package ‘stats’ v4.1.2. Relative importance of each regressor was estimated using *calc.relimp* function from the R package ‘relaimpo’ v2.2-6 [46]. The model *R^2^* was partitioned into regressor-specific contributions using the “averaging over orderings” approach [47]. Confidence intervals for regressor contributions were obtained by bootstrap resampling using *boot.relimp* function from the R package ‘relaimpo’ v2.2-6 [46]. Across 1,000 iterations, full observation vectors were resampled and regressor contributions were recalculated.

### STRING protein-protein interaction network

The protein-protein interaction network of the RBP of interest was obtained from STRING v11.5 [26]. Only edges with a confidence score ≥0.7 and supported by physical interaction evidence were retained.

### Cell culture and transfection

PC3 cells (ATCC CRL-7934; RRID: CVCL_0035) were obtained from ATCC, and cell line identity was verified by short tandem repeat (STR) profiling (DDC Medical). Cells were cultured at 37 °C in a humidified incubator with 5% CO₂ and maintained at sub-confluency in RPMI-1640 medium (21875-034, Gibco) or DMEM (41966-029, Gibco) containing 2 mM L-glutamine, supplemented with 10% foetal calf serum (Gibco), 100 units/mL penicillin and 100 μg/mL streptomycin (15140-122, Gibco). Cells were routinely tested for mycoplasma contamination. siRNA duplexes were commercially designed (ON-TARGETplus, Dharmacon Horizon Discovery) and are listed in Additional file 2: Table S6. Transfections siRNA duplexes were performed as previously described [31] using Lipofectamine RNAiMAX (13778-075, Thermo Fisher Scientific) following the manufacturers’ instructions. Briefly, PC3 cells were transfected with siRNA at a final concentration of 20 nM against either a control sequence or specifically targeting HNRNPK and were harvested 72 hours post-transfection for subsequent analysis.

### RNA-seq library preparation and data processing

RNA-seq library preparation, sequencing, and expression analyses were performed as previously described [31]. PC3 cells were lysed in Tri Reagent (AM9738, Invitrogen), and total RNA was extracted by phase separation using 1-bromo-3-chloropropane. To remove residual genomic DNA, RNA was treated with DNase I and purified using the RNA Clean & Concentrator kit (R1013, Zymo Research). RNA concentration was measured using a Qubit 4 Fluorometer (Q33238, Thermo Fisher Scientific), and RNA integrity was assessed with the RNA 6000 Nano kit (5067-1511, Agilent Technologies) on a 2100 Bioanalyzer (G2939BA, Agilent Technologies). High-quality RNA samples (RNA integrity number >9.7) were selected for library preparation. RNA-seq libraries were generated from 1μg of RNA using the TruSeq Stranded Total RNA Library Prep kit (20040529, Illumina), according to manufacturer’s recommendations. PC3 libraries were sequenced on the NovaSeq6000 (Illumina) in 101nt-long paired-end read modality.

Raw sequencing reads were aligned to the human genome reference GENCODE GRCh37 version 28 [42] using STAR v.2.7.3a [48] in basic two-pass mode using the following parameters: --alignInsertionFlush Right --outSAMstrandField intronMotif --outSAMattributes NH HI NM MD AS XS --peOverlapNbasesMin 20 --peOverlapMMp 0.25 --chimSegmentMin 12 --chimJunctionOverhangMin 8 --chimOutJunctionFormat 1 --chimMultimapScoreRange 3 --chimScoreJunctionNonGTAG -4 --chimMultimapNmax 20 --chimNonchimScoreDropMin 10 --outFilterIntronStrands RemoveInconsistentStrands --outFilterMultimapNmax 1 --bamRemoveDuplicatesType UniqueIdentical. Gene level were estimated using featureCounts (Subread v. 2.0.0) [49] with -p, -B and -s 2 parameters. Fragment counts were finally normalized as transcripts per million reads. The R package ‘DESeq2’ v1.34.0 was used to quantify differential gene expression [36] between siRNA-treated and control samples. Genes with |log2FC| ≥0.1 and adjusted p-value ≤0.1 were considered as differentially expressed (Additional file 2: Table S7). CEs were detected using rMATS v4.1.1 [48,50]. CEs with FDR <0.01 and |Δμ(Ψ)| >0.1 across conditions were defined as differentially spliced. CEs with FDR >0.1, |Δμ(Ψ)| <0.005, and not labeled as “alt” in UCSC hg19 “knownAlt” database [38] were considered as constitutive (Additional file 2: Table S8).

## Availability of data and materials

The current source code has been deposited on GitHub and is publicly available at GitHub at https://github.com/ceredamatteo-lab/theRNAMaRs and through Zenodo at https://doi.org/10.5281/zenodo.18427940. The code and data used to generate results and figures used in the manuscript is archived on Zenodo (DOI:10.5281/zenodo.18414809). The raw sequencing data of HNRNPK knockdown experiment in PC3 cell line generated in this study have been deposited in NCBI Sequence Read Archive database under the accession code PRJNA1400590 and are publicly available. All remaining information can be found in the Additional files.

## Supplementary information

Additional file 1: All supplementary figures for this study.

Additional file 2: All supplementary tables for this study.

## Acknowledgements

We thank Prof Jernej Ule (Francis Crick Institute, UK) for providing iCount processed data of ENCODE eCLIP experiments, and Dr Serena Peirone for reading the manuscript and useful discussion.

## Funding

M.C. is supported by AIRC under BRIDGE 2023 ID 28739 project and the Compagnia di San Paolo Foundation. M.G. and T.B. are PhD students within the European School of Molecular Medicine (SEMM). T.B. is supported by the UniverLecco Association.

## Author information

Mariachiara Grieco, Tommaso Becchi and Gabriele Boscagli contributed equally to this work.

## Contributions

Conceptualization, U.P. and M.C.; methodology, F.P., G.B., and M.C.; software, F.P., G.B., T.B., M.G., and M.C.; validation, F.P., G.B., and L.C.; formal analysis, F.P., T.B., and G.B.; investigation, F.P., G.B., T.B., M.G., U.P., and M.C.; resources, M.C.; data curation, F.P., G.B., T.B., and M.G.; writing - original draft, F.P., and M.C.; writing - review & editing, G.B., T.B., M.G., and M.C.; visualization, F.P., G.B., and M.C.; supervision, M.C.; project administration, M.C.; funding acquisition, M.C.. All authors read and approved the final manuscript.

## Ethics declarations

### Competing interests

The authors declare no competing interests.

## Notes

https://github.com/ceredamatteo-lab/theRNAmars

### Competing Interest Statement

The authors have declared no competing interest.

