## Supplementary Information for "RNAMaRs: an interpretable framework for inferring multivalent RNA Motifs and cognate Regulators of Splicing"

### Additional File 1

**Figure S1. Overview of the RNAMaRs framework for identifying splicing regulatory RBPs.**

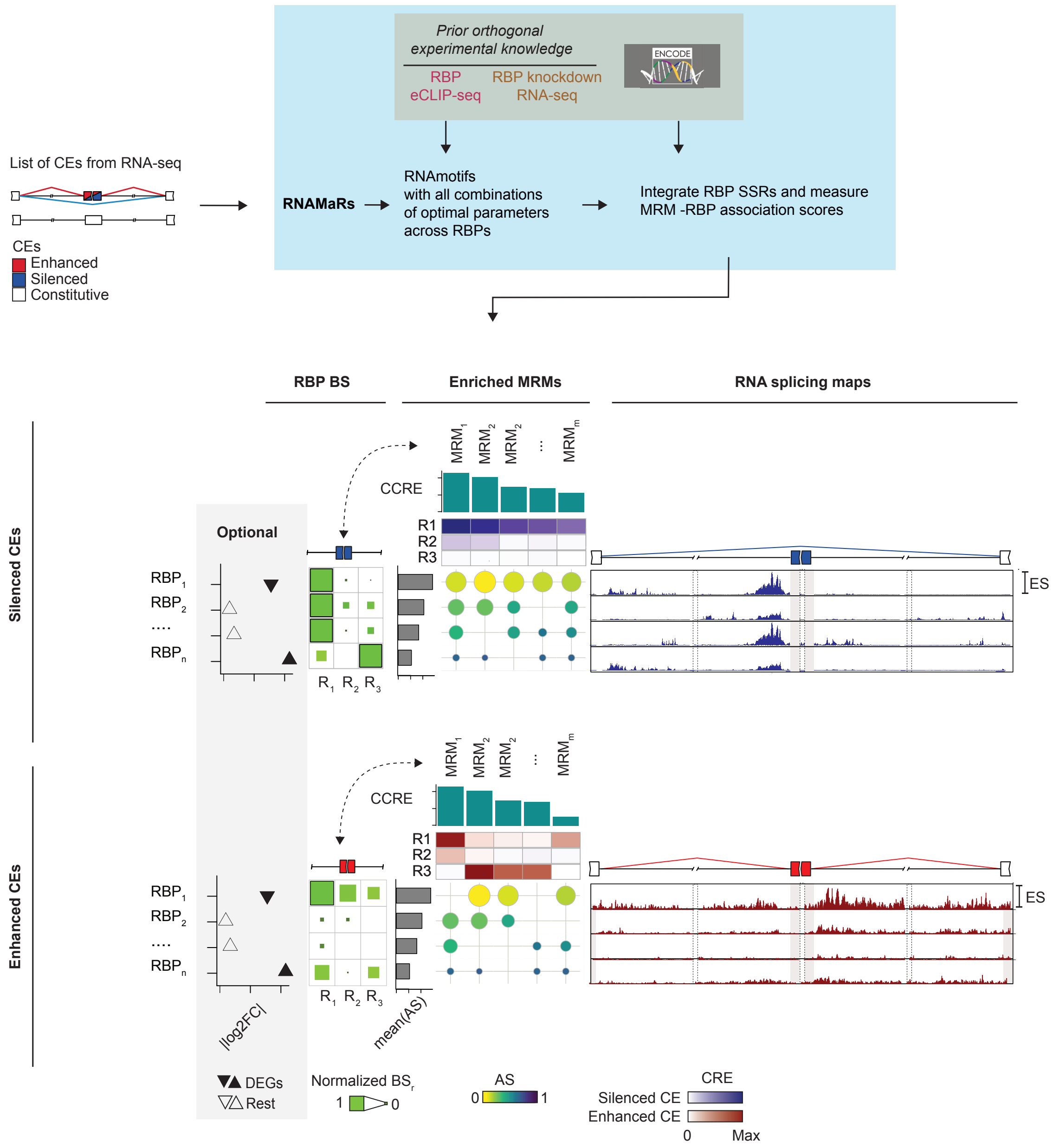


RNAMaRs takes as input a set of cassette exons (CEs) derived from RNA-seq and annotated as enhanced, silenced, or constitutive based on differential exon inclusion levels. The framework leverages prior orthogonal experimental knowledge from ENCODE, including eCLIP-seq binding profiles (physical association) and RNA-seq following RBP knockdown (functional evidence), together with optimized RNAmotifs settings to identify multivalent RNA motifs (MRMs). Next, RNAMaRs links MRMs to cognate RBPs by comparing MRM-specific and RBP-specific RNA splicing maps, producing MRM-RBP association scores (AS) alongside region-specific RBP binding scores (BS). Outputs include ranked candidate RBPs, enriched MRMs, and RNA splicing maps that summarize positional binding and regulatory signatures. For each MRM, combined regional enrichment (CRE) values are displayed as a top heatmap annotation, while cumulative CRE (CCRE) is visualized as a top bar plot. Differential gene expression (DEG) results for candidate RBPs are optionally included.

**Figure S2. Robustness and positional specificity of eCLIP-derived RBP binding profiles**

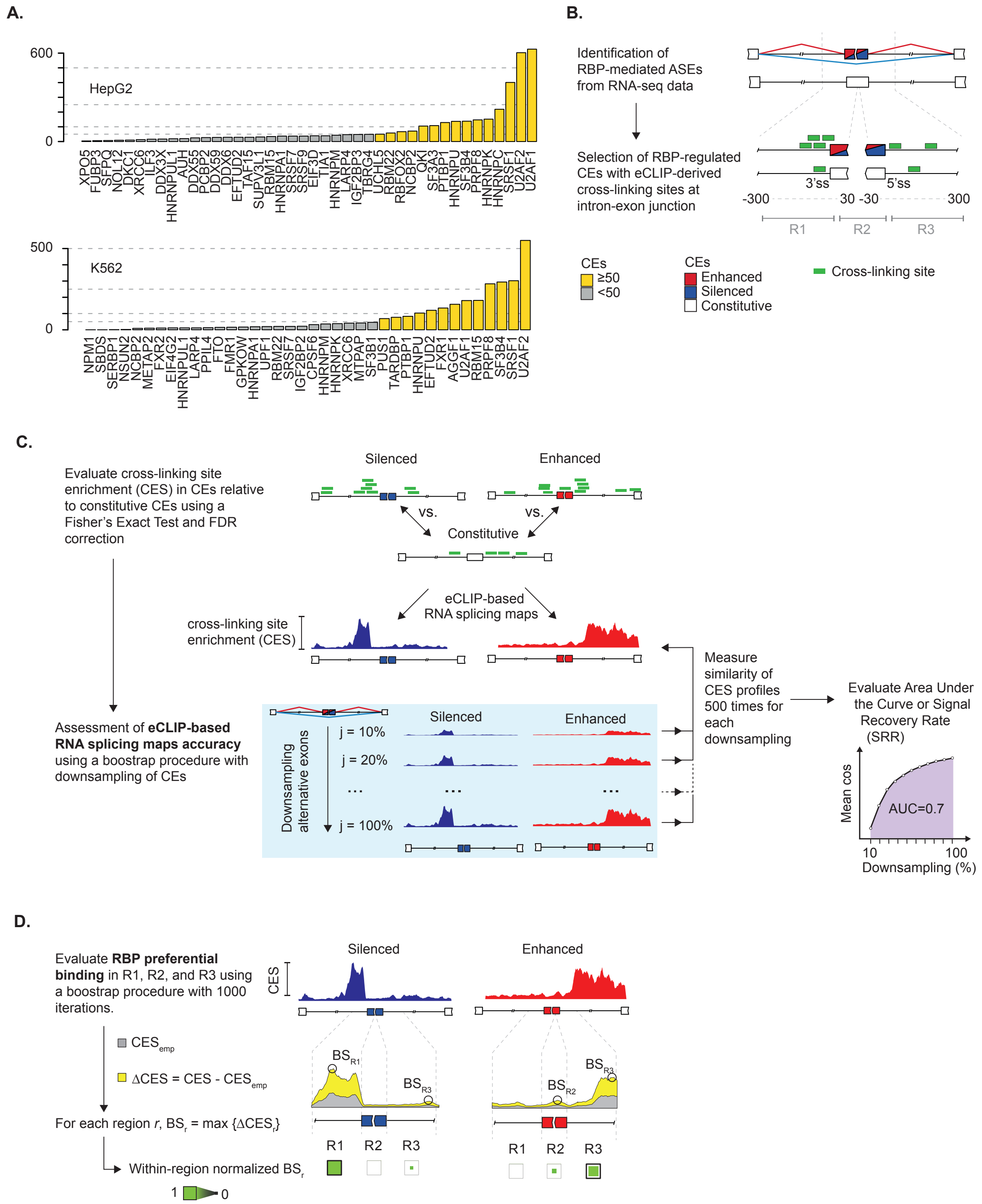


**A.** Number of alternative splicing events (ASEs) per RBP in HepG2 and K562 cells with cross-linking sites at intron/exon junctions. Yellow bars represent RBPs with at least 50 regulated cassette exons (CEs). **B.** Overview of the framework for the selection of CEs regulated by RBP binding. **C.** Schematic of the computation of CES profiles (top) and evaluation of their robustness through downsampling for SRR calculation (bottom). **D.** Overview of the framework for evaluating RBP preferential binding and quantifying the binding score (BS) through comparison with a background obtained via bootstrapping.

**Figure S3. Overview of the framework for the ROC-guided selection of optimal RNAmotifs parameters.**

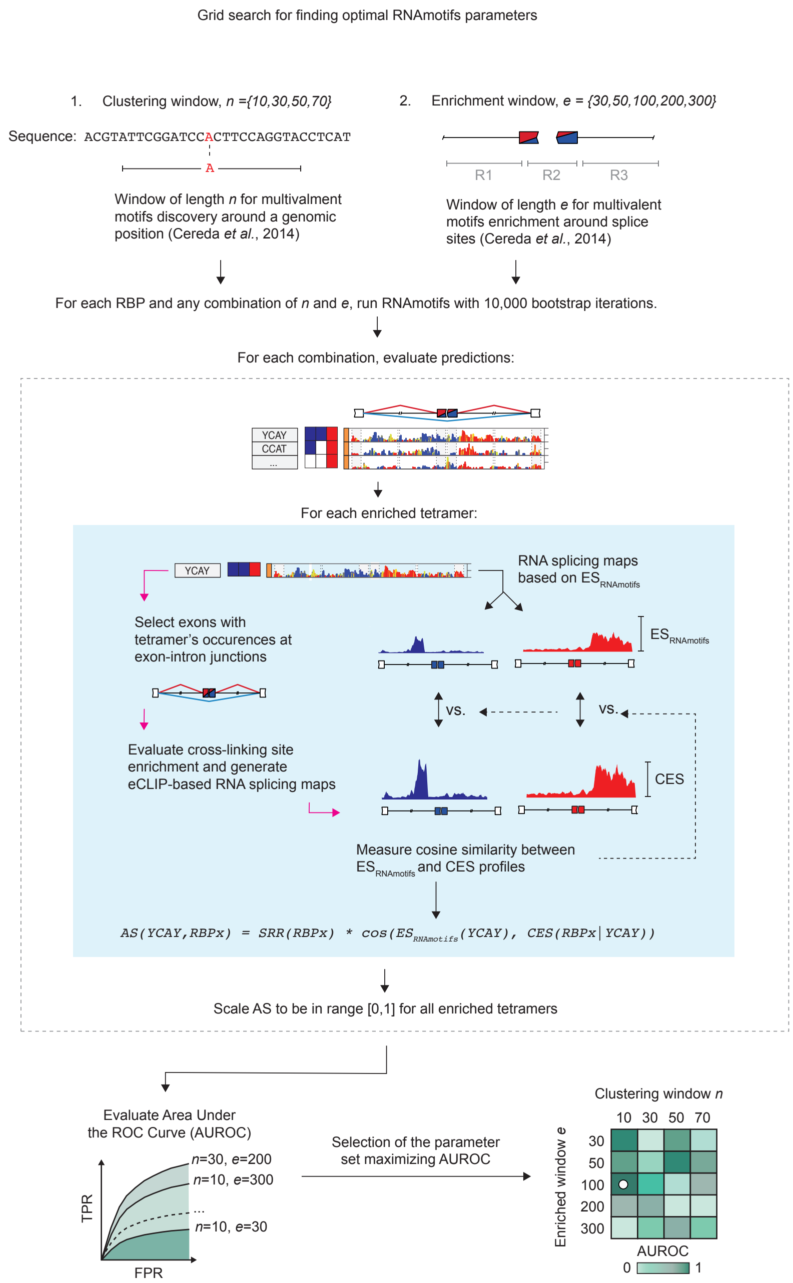


For each RBP, RNAmotifs results are generated using 20 different combinations of clustering (n) and enrichment (e) windows. Based on RNAmotifs results, the association score (AS) for each MRM is computed by comparing the enrichment score (ES) and cross-linking site enrichment (CES) profiles restricted to exons containing the MRM at exon/intron junctions. AS values are scaled to the [0,1] range and used to compute combination-specific AUROC values.

**Figure S4. Optimization of RNAmotifs parameters and evaluation of MRM-RBP association scores.**

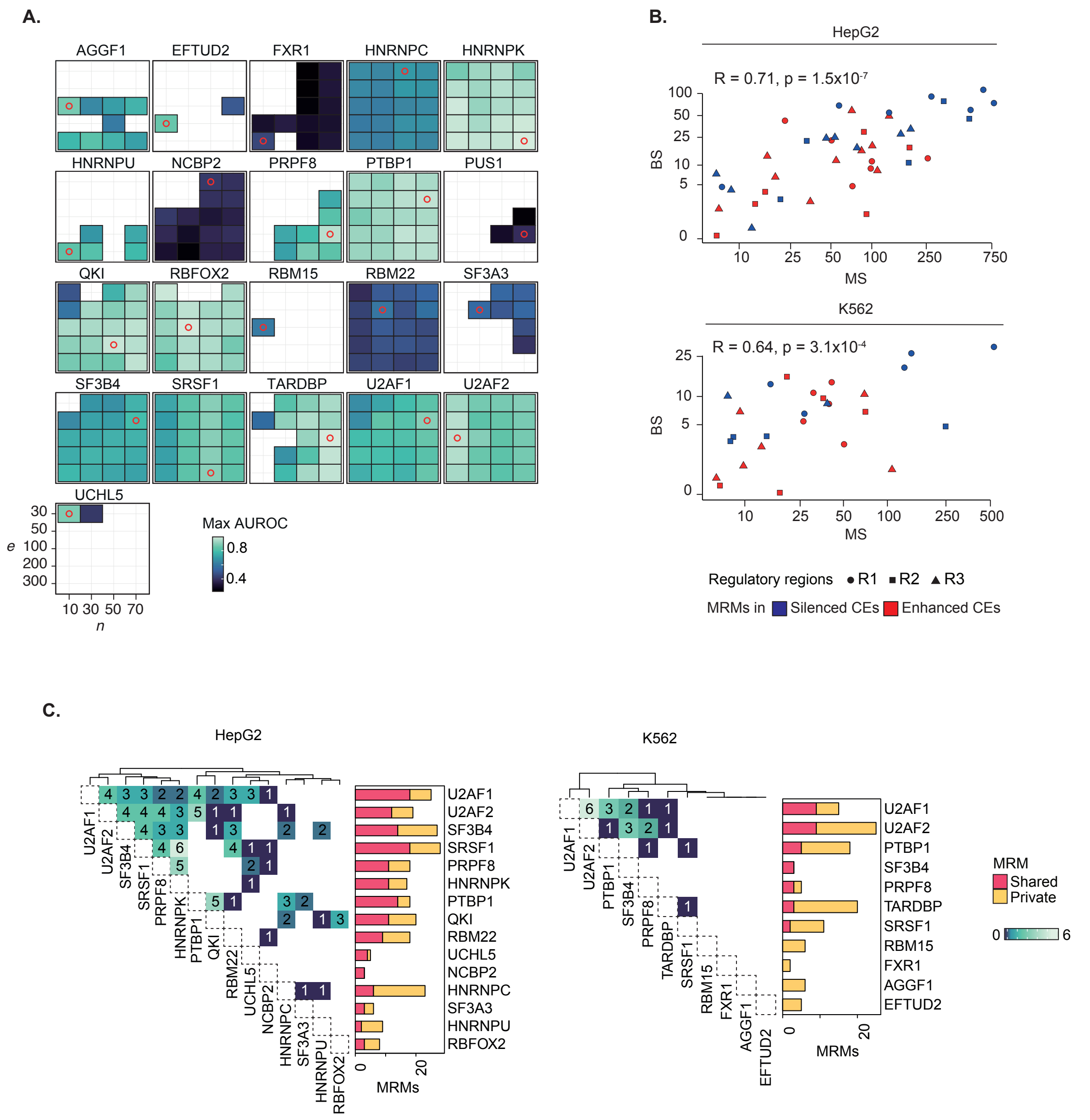


**A.** Heatmaps summarizing AUROC values obtained for each combination of clustering (n) and enrichment (e) window for each RBP. For each combination, the maximum AUROC across regulation classes and cell lines is reported. Only values derived from RNAmotifs results with at least five enriched MRMs are included. For each RBP, the maximum AUROC across all combinations is highlighted with a red circle. **B.** Relationship between raw binding (BS) and MRM (MS) values for each RBP across R1-R3 regions and regulation classes, stratified by cell line. Point shape indicates the regulatory regions, while the color represents the regulation classes. Only cases in which both scores are greater than 0 are included. For each cell line, the overall Pearson correlation coefficient is reported. **C.** Overlap of enriched MRMs across RBPs in each cell line, distinguishing shared and RBP-private motifs.

**Figure S5. RNAMaRs results on PTBP1-silenced exons in HepG2 and K562 cells.**

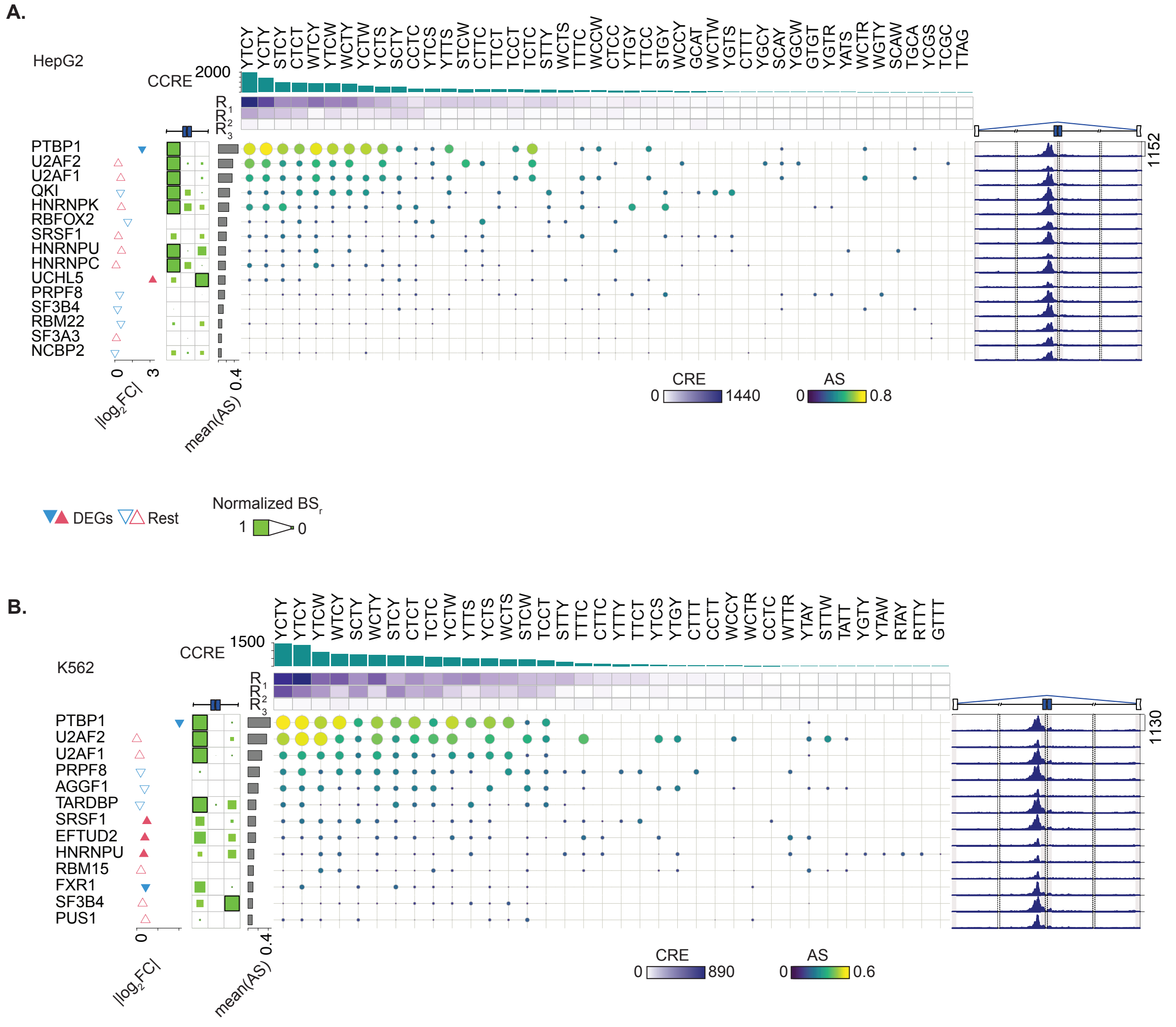


Complete RNAMaRs outputs for PTBP1-regulated silenced exons in HepG2 (**A**) and K562 (**B**). The visualization integrates normalized binding scores (BS) across splice-proximal regions (R1-R3) (left), MRM-RBP association scores (AS), cumulative combined regional enrichment (CCRE) across enriched MRMs (top), and the corresponding RNA splicing maps (right). Differential gene expression (DEG) values of corresponding RBP upon knockdown are shown as triangles. The upper and lower vertices indicated up- and down-regulation, respectively, and filled triangles denoted differential expression changes (*i.e.*, |log2FC| ≥0.1 and adjusted p-values ≤0.1). Barplots indicate the mean AS per the candidate regulator.

**Figure S6. RNAMaRs identifies HNRNPK as a splicing regulator in HNRNPK-depleted PC3 cells**

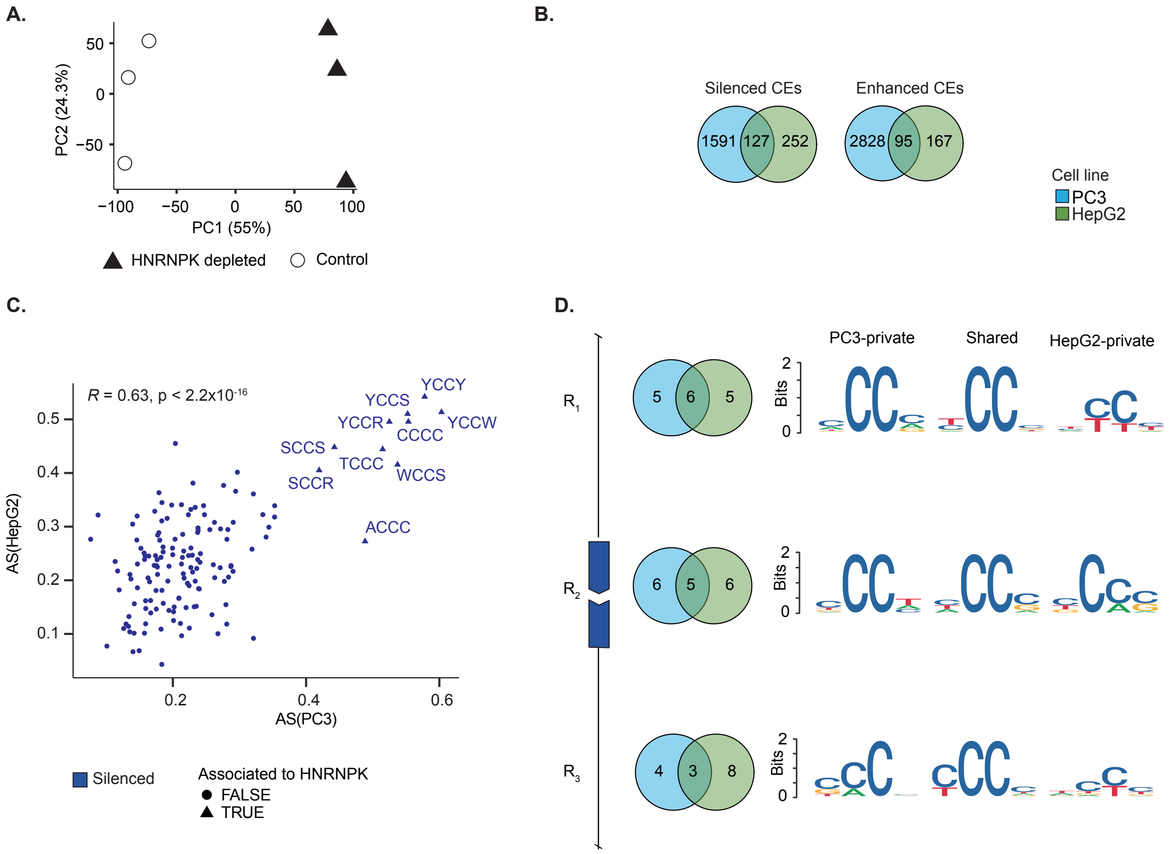


**A.** Scatter plot of the first two components of principal component analysis (PCA) for PC3 dataset. Percentage of variance explained by each component is reported on each axis. **B.** Venn diagram showing the overlap of silenced and enhanced differentially spliced CEs between PC3 and HepG2 cell lines. **C.** Relationship between association scores (AS) of each tetramer in PC3 and HepG2 cells for silenced exons. Tetramers identified as significantly associated to HNRNPK in both datasets are indicated with triangles. The overall Pearson correlation coefficient is reported. **D.** Venn diagram showing the overlap of MRMs enriched in silenced exons between PC3 and HepG2 cell lines, along with the corresponding sequence logos summarizing MRMs shared and cell line-specific, stratified by splice-proximal regions (R1-R3).

**Figure S7. RNAMaRs results for HNRNPC-regulated exons in HepG2 cells.**

**A.**

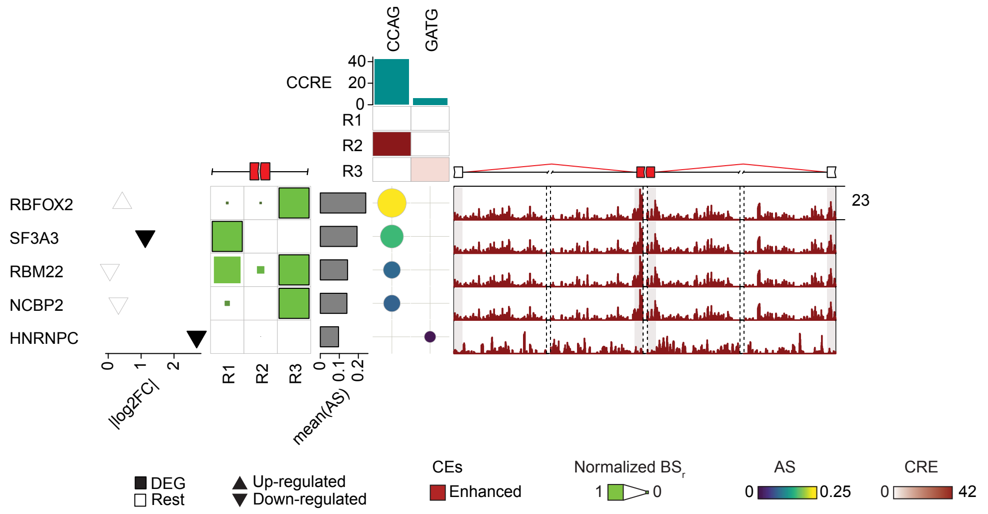


**B.**

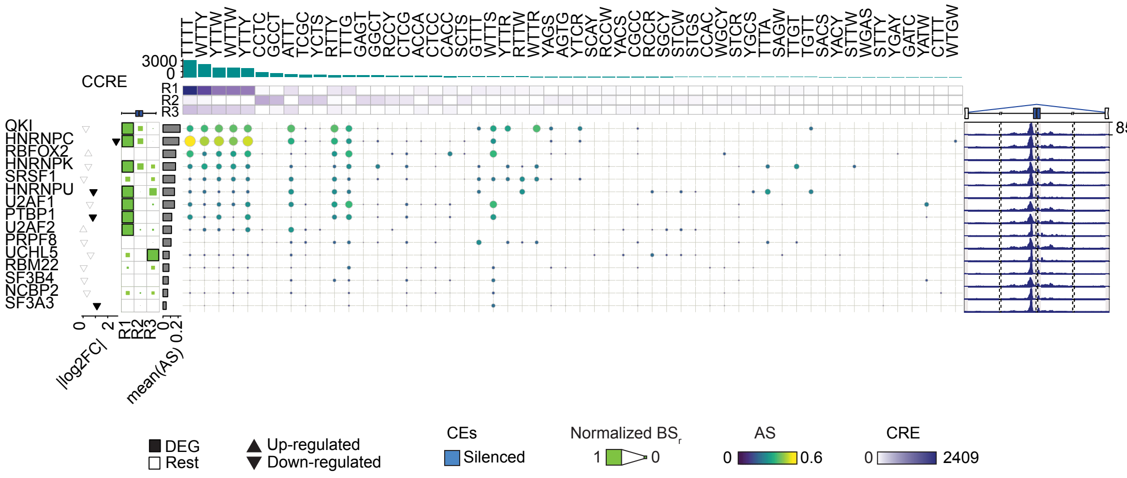


Complete RNAMaRs outputs for HNRNPC-regulated enhanced **(A)** and silenced **(B)** exons in HepG2 cells. The visualization integrates normalized binding scores (BS) across splice-proximal regions (R1-R3) (left), MRM-RBP association scores (AS), cumulative combined regional enrichment (CCRE) across enriched MRMs (top), and the corresponding RNA splicing maps (right). Differential gene expression (DEG) values of corresponding RBP upon knockdown are shown as triangles. The upper and lower vertices indicated up- and down-regulation, respectively, and filled triangles denoted differential expression changes (*i.e.*, |log2FC| ≥0.1 and adjusted p-values ≤0.1). Barplots indicate the mean AS per the candidate regulator.

**Figure S8. RNAMaRs results for HNRNPK-regulated exons in HepG2 cells.**

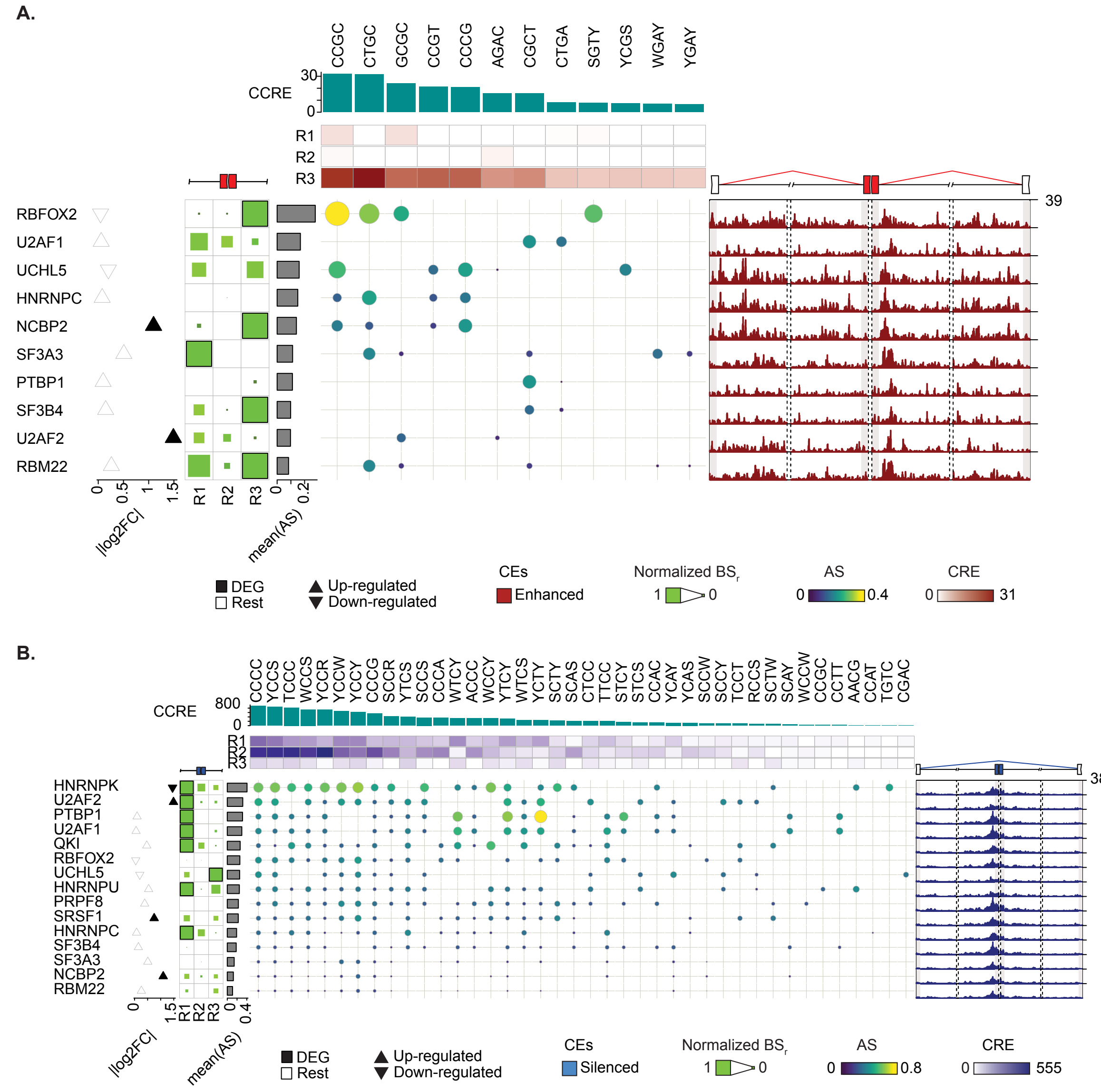


Complete RNAMaRs outputs for HNRNPK-regulated enhanced **(A)** and silenced **(B)** exons in HepG2 cells. The visualization integrates normalized binding scores (BS) across splice-proximal regions (R1-R3) (left), MRM-RBP association scores (AS), cumulative combined regional enrichment (CCRE) across enriched MRMs (top), and the corresponding RNA splicing maps (right). Differential gene expression (DEG) values of corresponding RBP upon knockdown are shown as triangles. The upper and lower vertices indicated up- and down-regulation, respectively, and filled triangles denoted differential expression changes (*i.e.*, |log2FC| ≥0.1 and adjusted p-values ≤0.1). Barplots indicate the mean AS per the candidate regulator.

**Figure S9. RNAMaRs results for HNRNPU-regulated exons in HepG2 cells.**

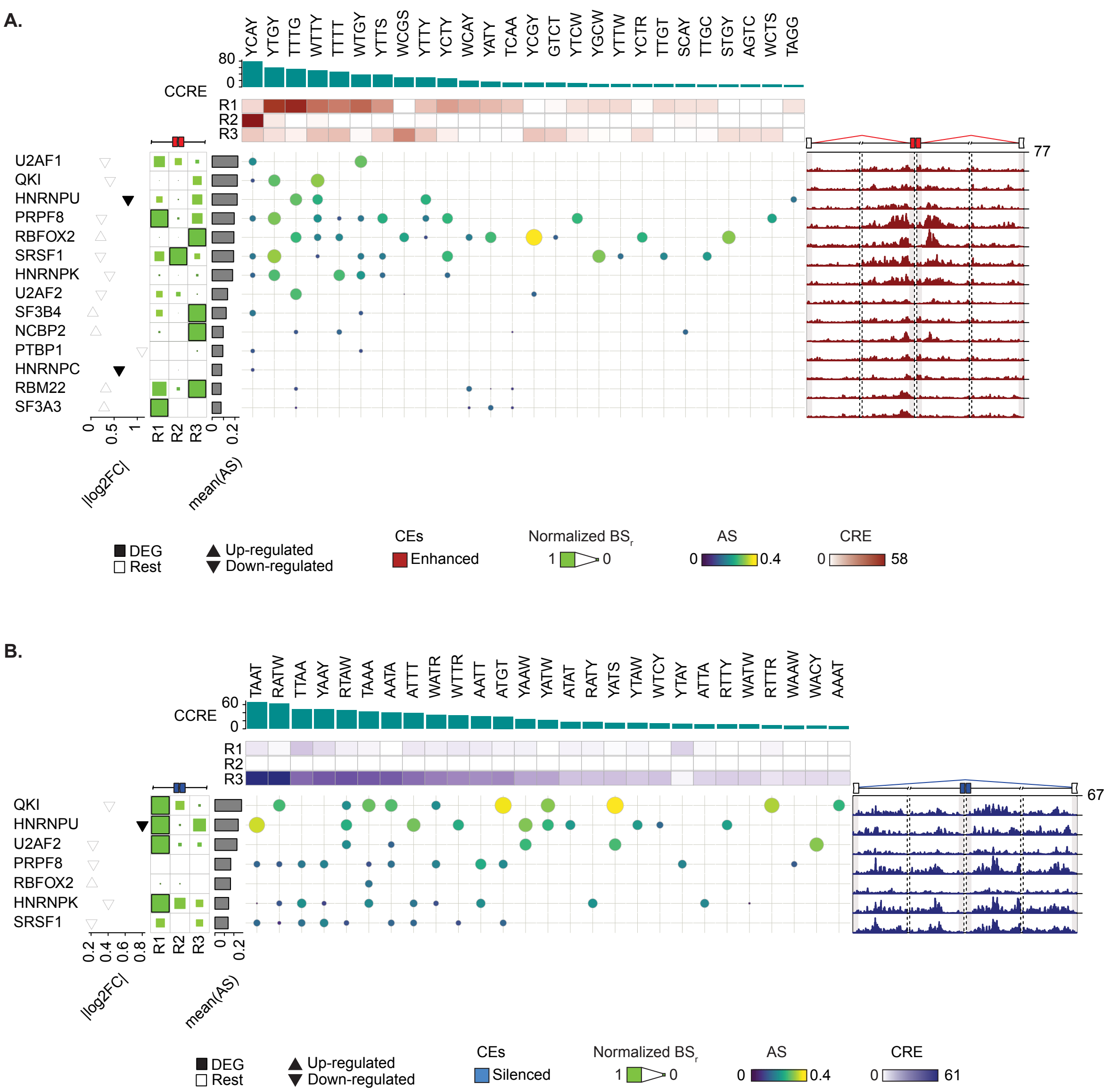


Complete RNAMaRs outputs for HNRNPU-regulated enhanced **(A)** and silenced **(B)** exons in HepG2 cells. The visualization integrates normalized binding scores (BS) across splice-proximal regions (R1-R3) (left), MRM-RBP association scores (AS), cumulative combined regional enrichment (CCRE) across enriched MRMs (top), and the corresponding RNA splicing maps (right). Differential gene expression (DEG) values of corresponding RBP upon knockdown are shown as triangles. The upper and lower vertices indicated up- and down-regulation, respectively, and filled triangles denoted differential expression changes (*i.e.*, |log2FC| ≥0.1 and adjusted p-values ≤0.1). Barplots indicate the mean AS per the candidate regulator.

**Figure S10. RNAMaRs results for NCBP2-regulated exons in HepG2 cells.**

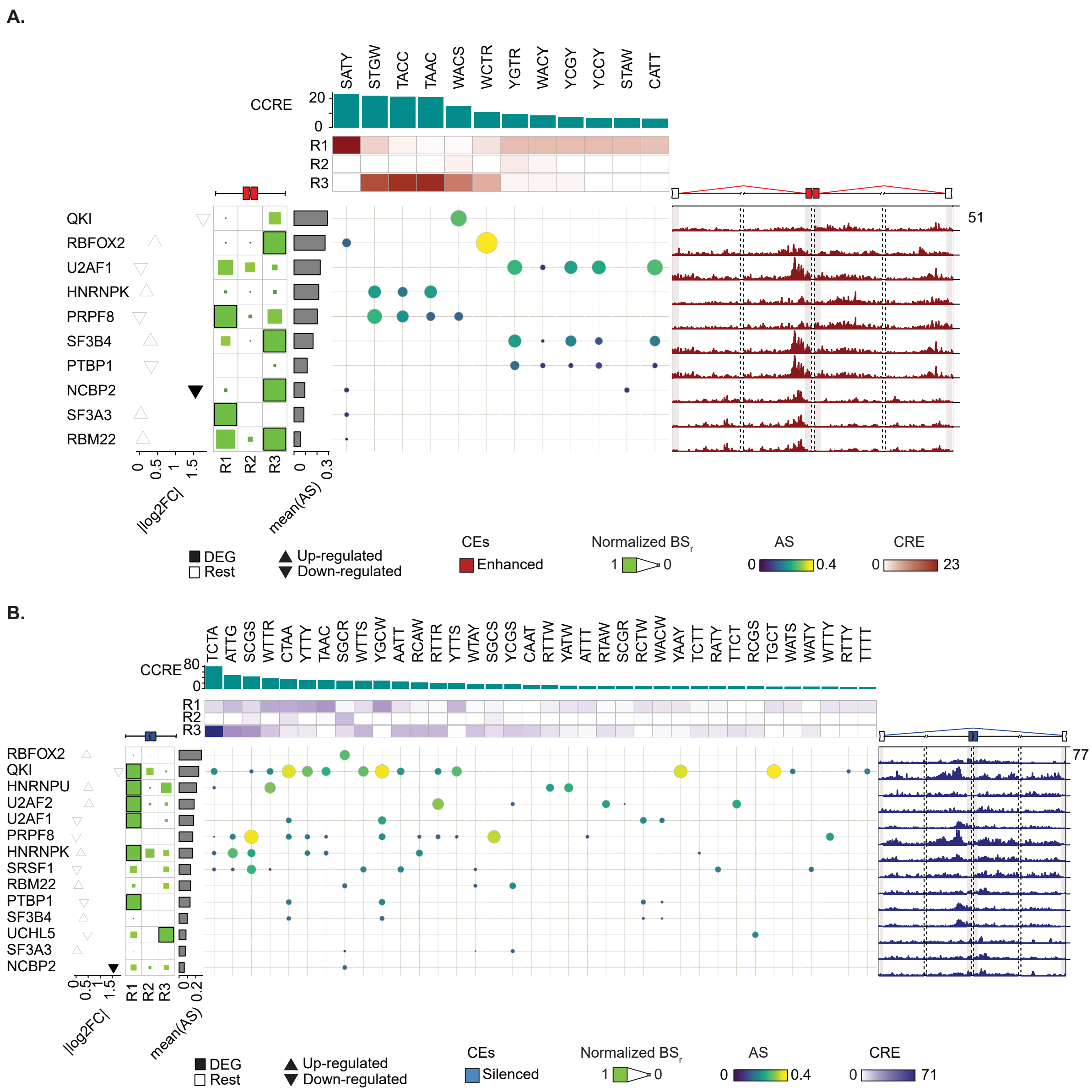


Complete RNAMaRs outputs for NCBP2-regulated enhanced **(A)** and silenced **(B)** exons in HepG2 cells. The visualization integrates normalized binding scores (BS) across splice-proximal regions (R1-R3) (left), MRM-RBP association scores (AS), cumulative combined regional enrichment (CCRE) across enriched MRMs (top), and the corresponding RNA splicing maps (right). Differential gene expression (DEG) values of corresponding RBP upon knockdown are shown as triangles. The upper and lower vertices indicated up- and down-regulation, respectively, and filled triangles denoted differential expression changes (*i.e.*, |log2FC| ≥0.1 and adjusted p-values ≤0.1). Barplots indicate the mean AS per the candidate regulator.

**Figure S11. RNAMaRs results for PRPF8-regulated exons in HepG2 cells.**

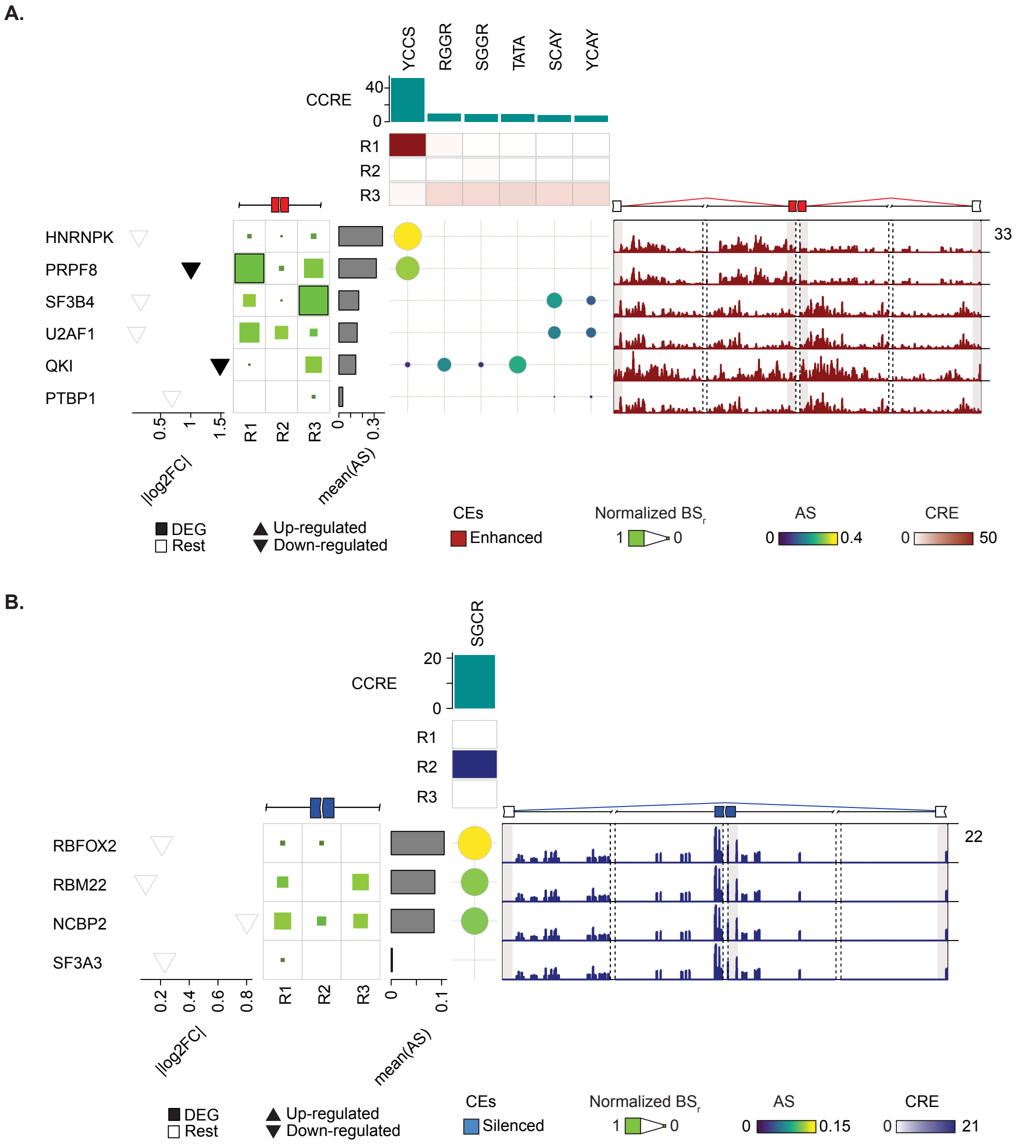


Complete RNAMaRs outputs for PRPF8-regulated enhanced **(A)** and silenced **(B)** exons in HepG2 cells. The visualization integrates normalized binding scores (BS) across splice-proximal regions (R1-R3) (left), MRM-RBP association scores (AS), cumulative combined regional enrichment (CCRE) across enriched MRMs (top), and the corresponding RNA splicing maps (right). Differential gene expression (DEG) values of corresponding RBP upon knockdown are shown as triangles. The upper and lower vertices indicated up- and down-regulation, respectively, and filled triangles denoted differential expression changes (*i.e.*, |log2FC| ≥0.1 and adjusted p-values ≤0.1). Barplots indicate the mean AS per the candidate regulator.

**Figure S12. RNAMaRs results for PTPB1-regulated exons in HepG2 cells.**

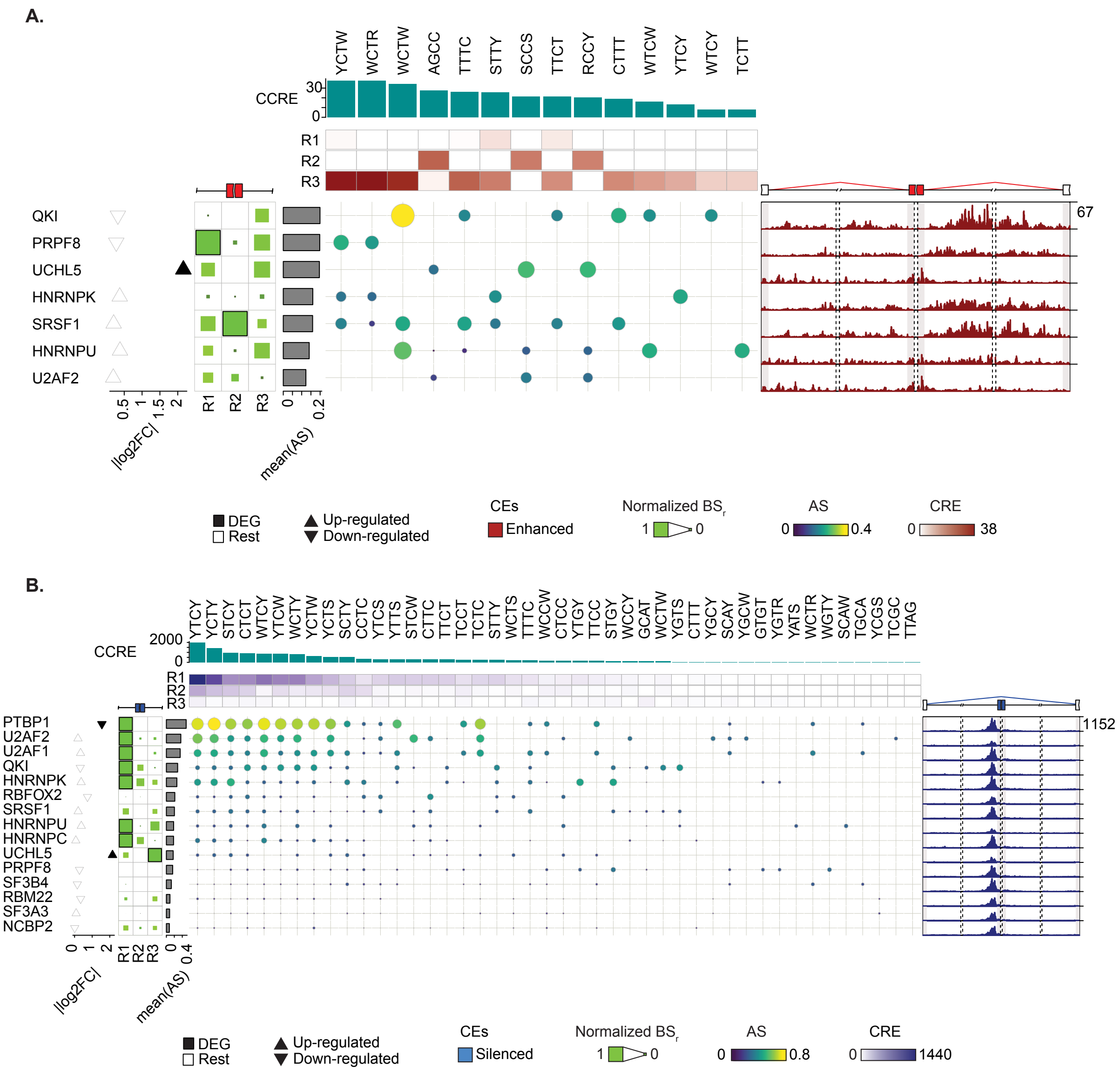


Complete RNAMaRs outputs for PTPB1-regulated enhanced **(A)** and silenced **(B)** exons in HepG2 cells. The visualization integrates normalized binding scores (BS) across splice-proximal regions (R1-R3) (left), MRM-RBP association scores (AS), cumulative combined regional enrichment (CCRE) across enriched MRMs (top), and the corresponding RNA splicing maps (right). Differential gene expression (DEG) values of corresponding RBP upon knockdown are shown as triangles. The upper and lower vertices indicated up- and down-regulation, respectively, and filled triangles denoted differential expression changes (*i.e.*, |log2FC| ≥0.1 and adjusted p-values ≤0.1). Barplots indicate the mean AS per the candidate regulator.

**Figure S13. RNAMaRs results for QKI-regulated exons in HepG2 cells.**

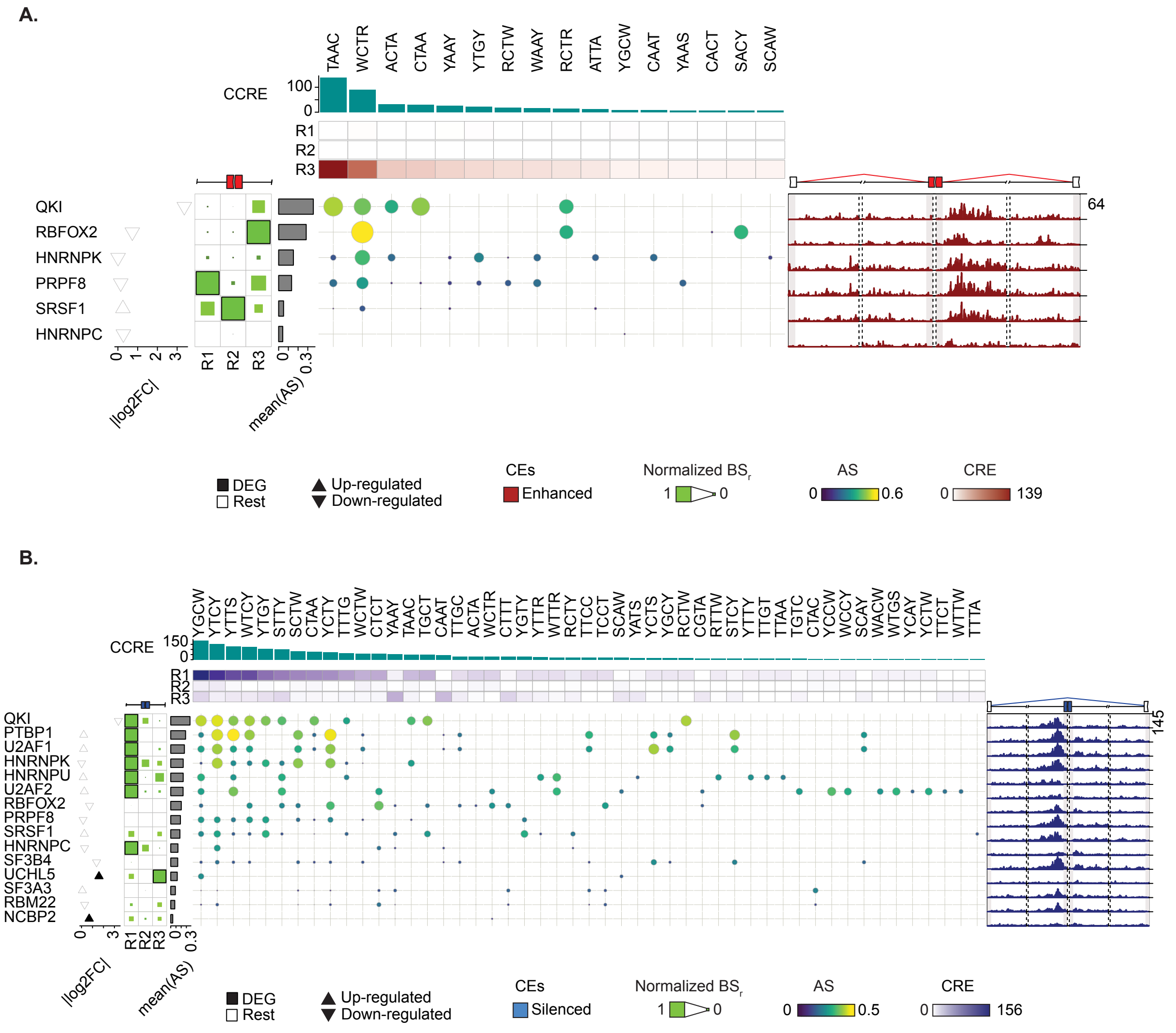


Complete RNAMaRs outputs for QKI-regulated enhanced **(A)** and silenced **(B)** exons in HepG2 cells. The visualization integrates normalized binding scores (BS) across splice-proximal regions (R1-R3) (left), MRM-RBP association scores (AS), cumulative combined regional enrichment (CCRE) across enriched MRMs (top), and the corresponding RNA splicing maps (right). Differential gene expression (DEG) values of corresponding RBP upon knockdown are shown as triangles. The upper and lower vertices indicated up- and down-regulation, respectively, and filled triangles denoted differential expression changes (*i.e.*, |log2FC| ≥0.1 and adjusted p-values ≤0.1). Barplots indicate the mean AS per the candidate regulator.

**Figure S14. RNAMaRs results for RBFOX2-regulated exons in HepG2 cells.**

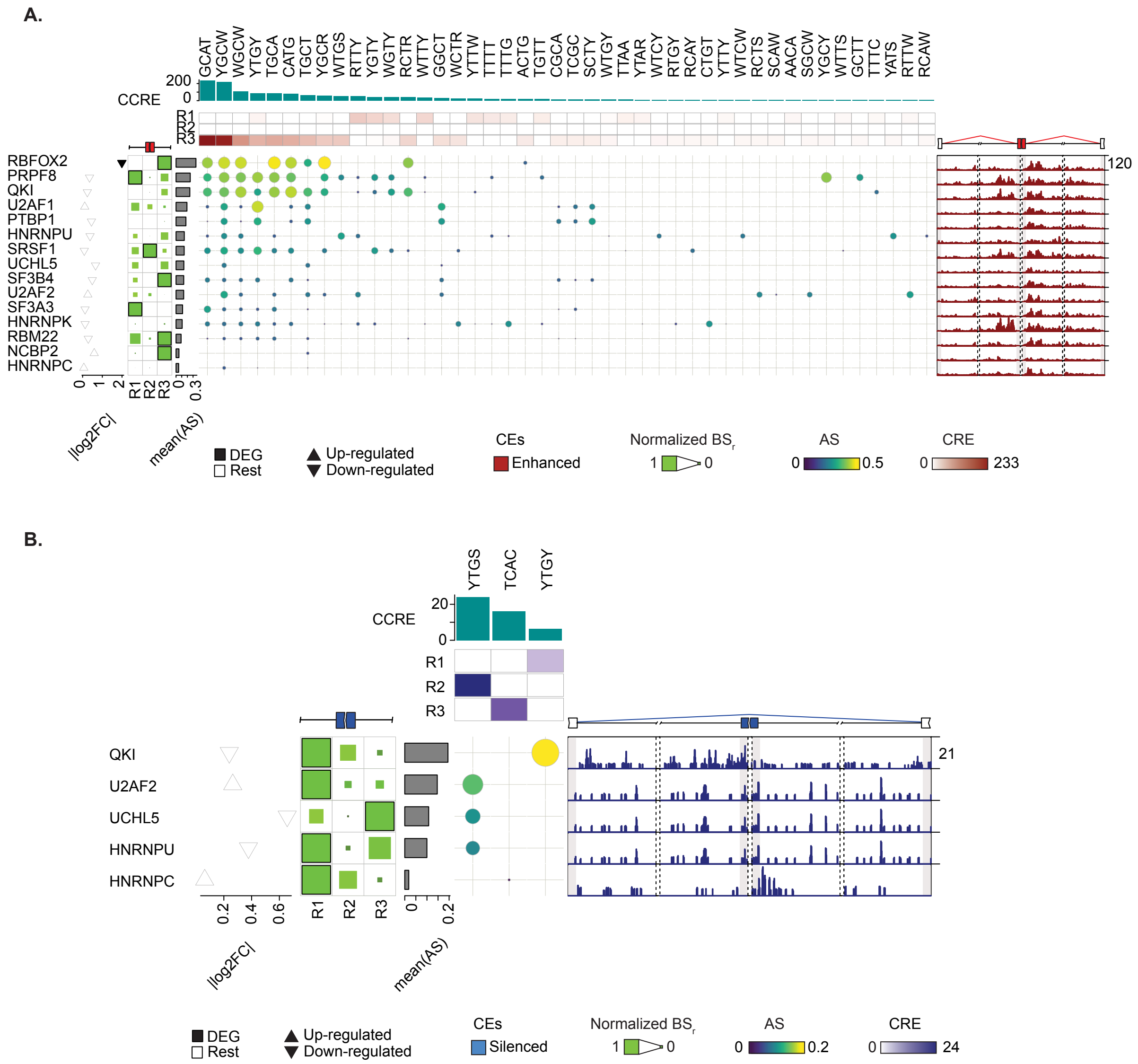


Complete RNAMaRs outputs for RBFOX2-regulated enhanced **(A)** and silenced **(B)** exons in HepG2 cells. The visualization integrates normalized binding scores (BS) across splice-proximal regions (R1-R3) (left), MRM-RBP association scores (AS), cumulative combined regional enrichment (CCRE) across enriched MRMs (top), and the corresponding RNA splicing maps (right). Differential gene expression (DEG) values of corresponding RBP upon knockdown are shown as triangles. The upper and lower vertices indicated up- and down-regulation, respectively, and filled triangles denoted differential expression changes (*i.e.*, |log2FC| ≥0.1 and adjusted p-values ≤0.1). Barplots indicate the mean AS per the candidate regulator.

**Figure S15. RNAMaRs results for RBM22-regulated exons in HepG2 cells.**

A.

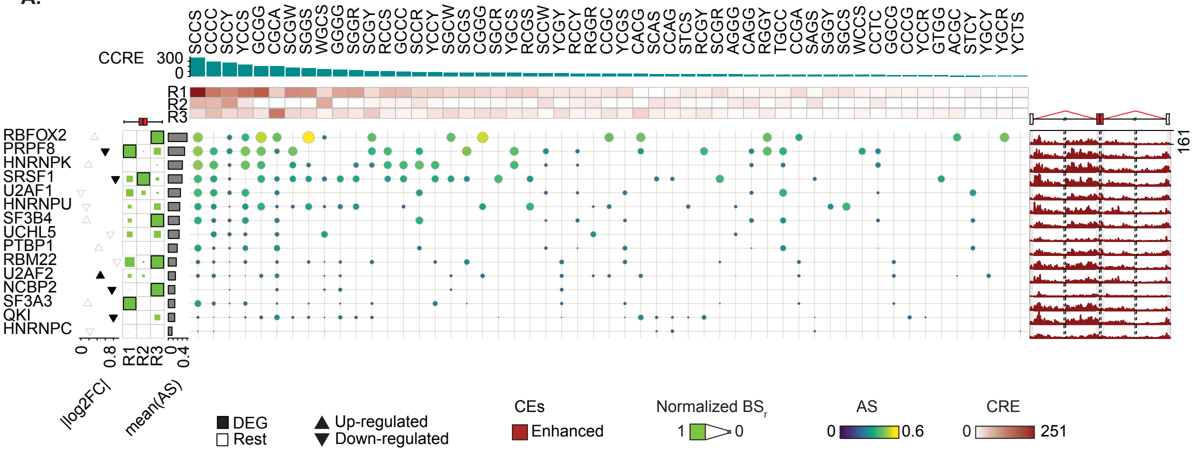


B.

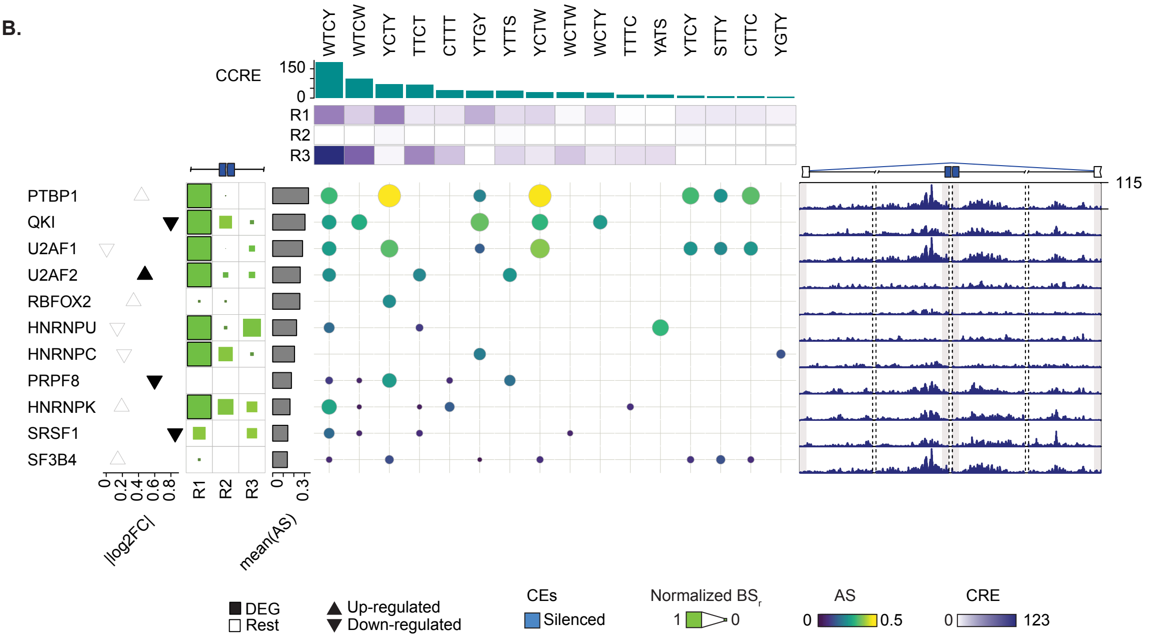


Complete RNAMaRs outputs for RBM22-regulated enhanced **(A)** and silenced **(B)** exons in HepG2 cells. The visualization integrates normalized binding scores (BS) across splice-proximal regions (R1-R3) (left), MRM-RBP association scores (AS), cumulative combined regional enrichment (CCRE) across enriched MRMs (top), and the corresponding RNA splicing maps (right). Differential gene expression (DEG) values of corresponding RBP upon knockdown are shown as triangles. The upper and lower vertices indicated up- and down-regulation, respectively, and filled triangles denoted differential expression changes (*i.e.*, |log2FC| ≥0.1 and adjusted p-values ≤0.1). Barplots indicate the mean AS per the candidate regulator.

**Figure S16. RNAMaRs results for SF3A3-regulated exons in HepG2 cells.**

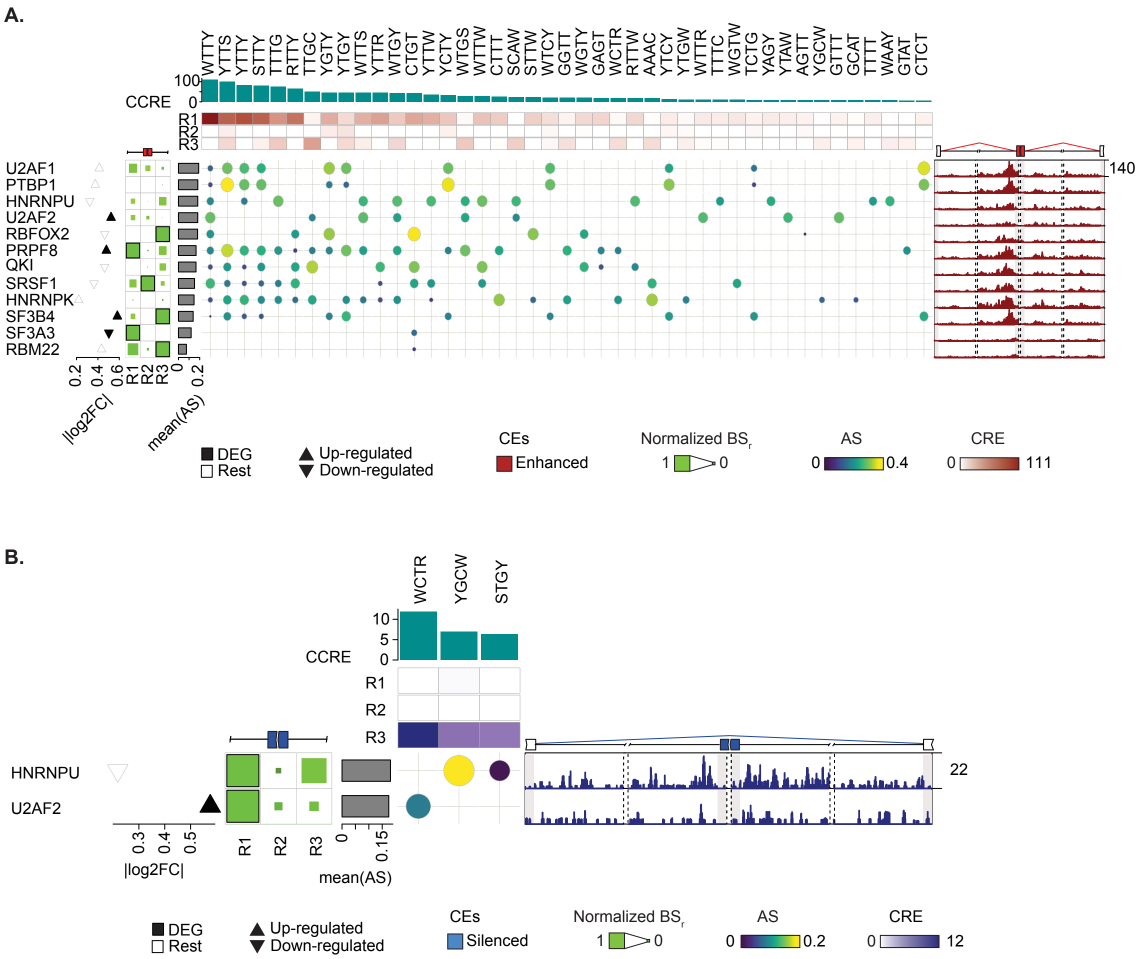


Complete RNAMaRs outputs for SF3A3-regulated enhanced **(A)** and silenced **(B)** exons in HepG2 cells. The visualization integrates normalized binding scores (BS) across splice-proximal regions (R1-R3) (left), MRM-RBP association scores (AS), cumulative combined regional enrichment (CCRE) across enriched MRMs (top), and the corresponding RNA splicing maps (right). Differential gene expression (DEG) values of corresponding RBP upon knockdown are shown as triangles. The upper and lower vertices indicated up- and down-regulation, respectively, and filled triangles denoted differential expression changes (*i.e.*, |log2FC| ≥0.1 and adjusted p-values ≤0.1). Barplots indicate the mean AS per the candidate regulator.

### Figure S17. RNAMaRs results for SF3B4-regulated exons in HepG2 cells.

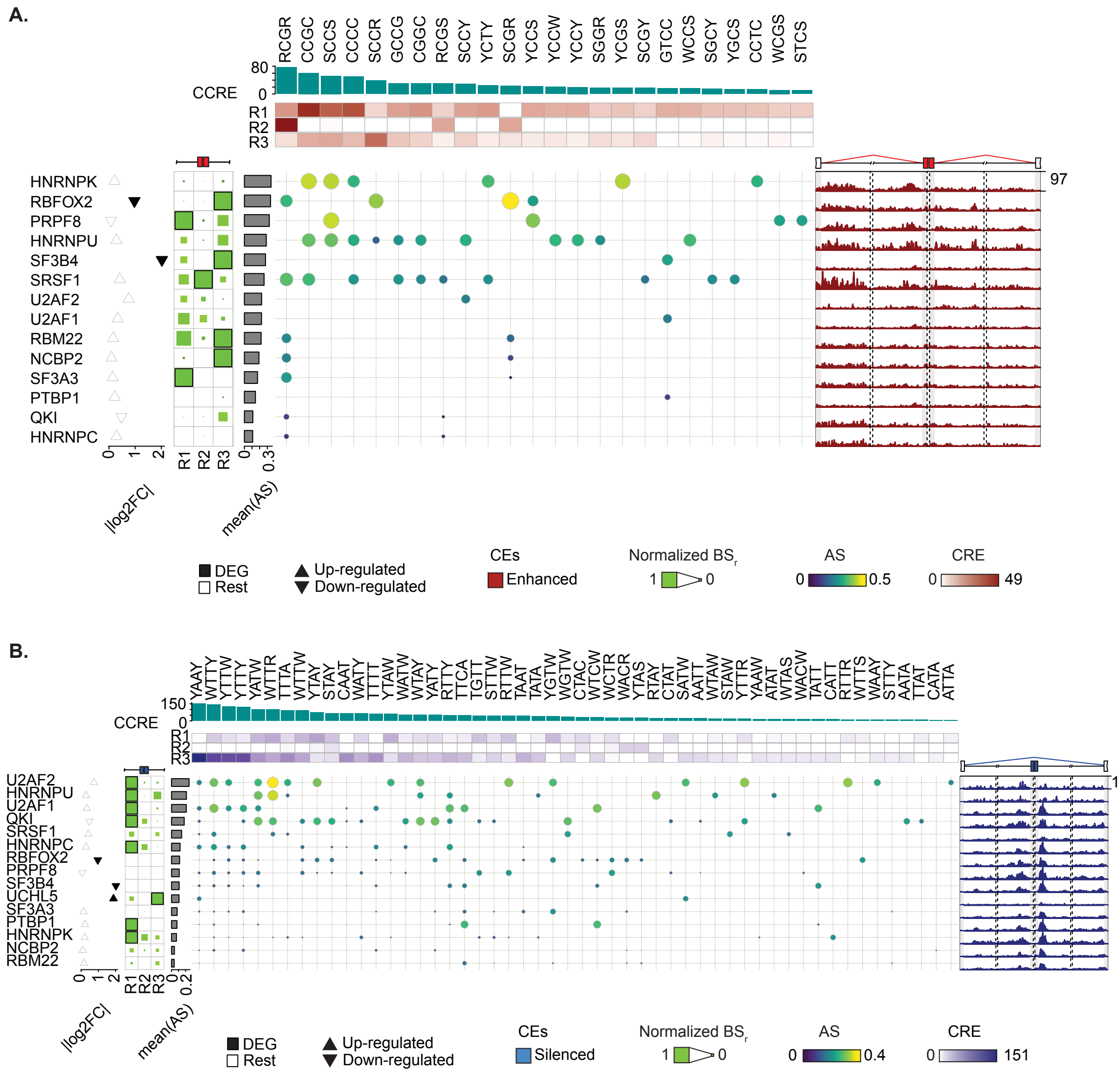


Complete RNAMaRs outputs for SF3B4-regulated enhanced **(A)** and silenced **(B)** exons in HepG2 cells. The visualization integrates normalized binding scores (BS) across splice-proximal regions (R1-R3) (left), MRM-RBP association scores (AS), cumulative combined regional enrichment (CCRE) across enriched MRMs (top), and the corresponding RNA splicing maps (right). Differential gene expression (DEG) values of corresponding RBP upon knockdown are shown as triangles. The upper and lower vertices indicated up- and down-regulation, respectively, and filled triangles denoted differential expression changes (*i.e.*, |log2FC| ≥0.1 and adjusted p-values ≤0.1). Barplots indicate the mean AS per the candidate regulator.

**Figure S18. RNAMaRs results for SRSF1-regulated exons in HepG2 cells.**

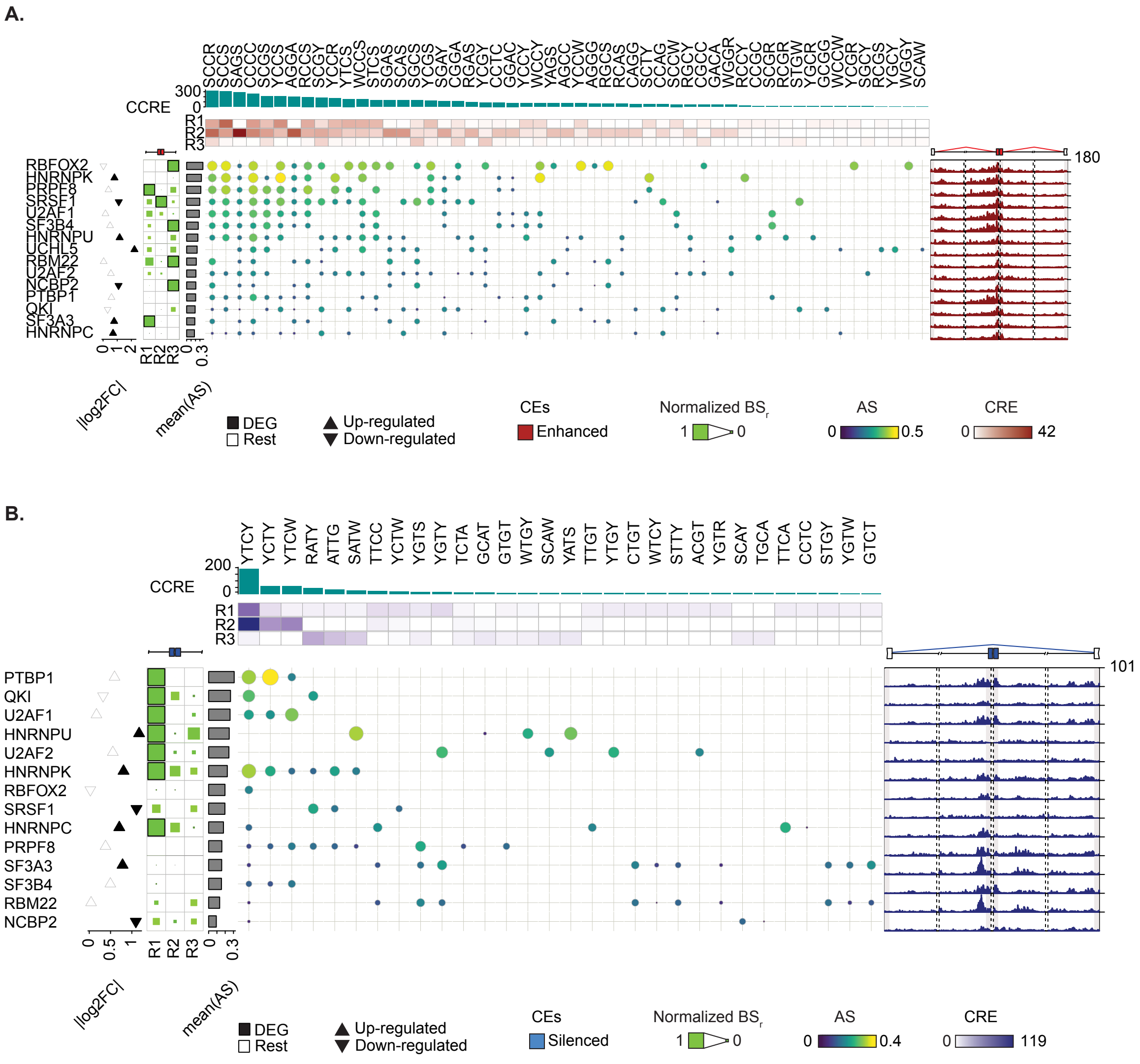


Complete RNAMaRs outputs for SRSF1-regulated enhanced **(A)** and silenced **(B)** exons in HepG2 cells. The visualization integrates normalized binding scores (BS) across splice-proximal regions (R1-R3) (left), MRM-RBP association scores (AS), cumulative combined regional enrichment (CCRE) across enriched MRMs (top), and the corresponding RNA splicing maps (right). Differential gene expression (DEG) values of corresponding RBP upon knockdown are shown as triangles. The upper and lower vertices indicated up- and down-regulation, respectively, and filled triangles denoted differential expression changes (*i.e.*, |log2FC| ≥0.1 and adjusted p-values ≤0.1). Barplots indicate the mean AS per the candidate regulator.

### Figure S19. RNAMaRs results for U2AF1-regulated exons in HepG2 cells.

**A.**

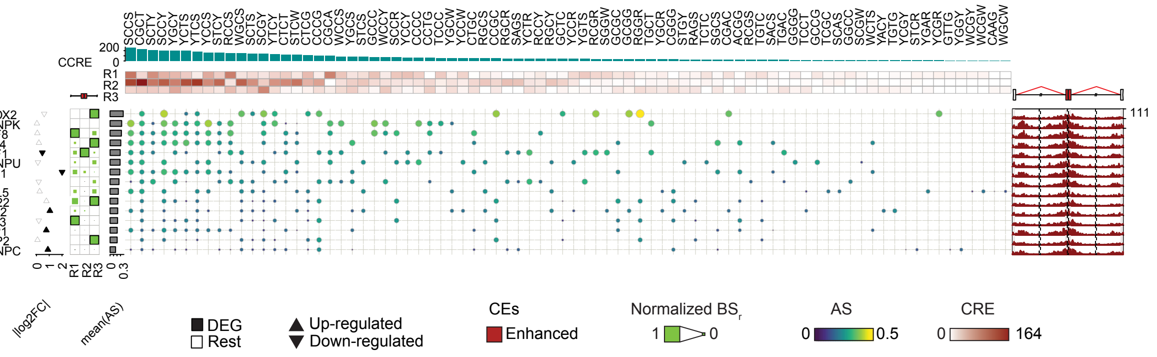


**B.**

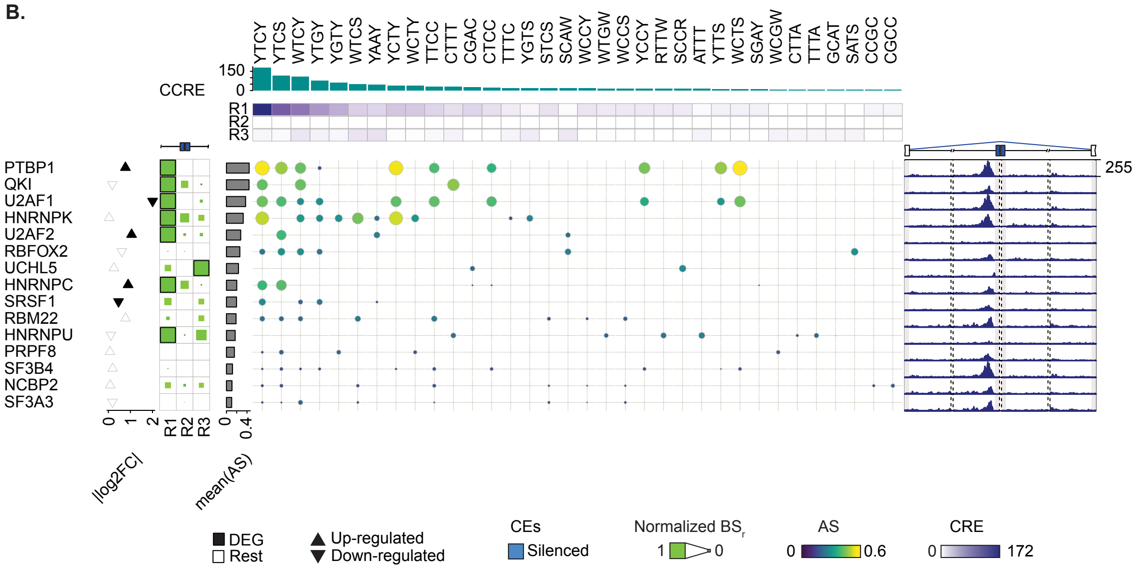


Complete RNAMaRs outputs for U2AF1-regulated enhanced **(A)** and silenced **(B)** exons in HepG2 cells. The visualization integrates normalized binding scores (BS) across splice-proximal regions (R1-R3) (left), MRM-RBP association scores (AS), cumulative combined regional enrichment (CCRE) across enriched MRMs (top), and the corresponding RNA splicing maps (right). Differential gene expression (DEG) values of corresponding RBP upon knockdown are shown as triangles. The upper and lower vertices indicated up- and down-regulation, respectively, and filled triangles denoted differential expression changes (*i.e.*, |log2FC| ≥0.1 and adjusted p-values ≤0.1). Barplots indicate the mean AS per the candidate regulator.

**Figure S20. RNAMaRs results for U2AF2-regulated exons in HepG2 cells.**

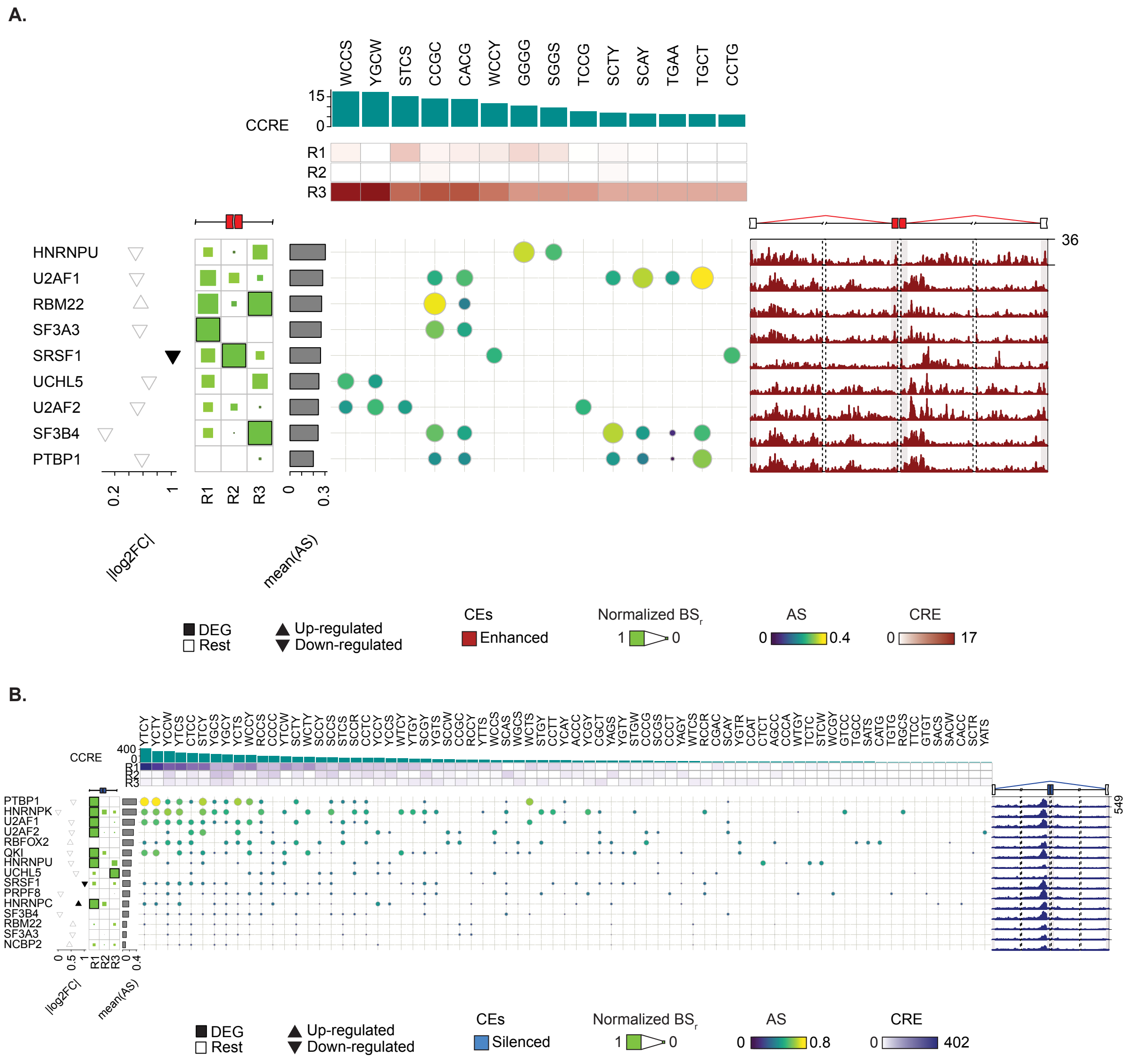


Complete RNAMaRs outputs for U2AF2-regulated enhanced **(A)** and silenced **(B)** exons in HepG2 cells. The visualization integrates normalized binding scores (BS) across splice-proximal regions (R1-R3) (left), MRM-RBP association scores (AS), cumulative combined regional enrichment (CCRE) across enriched MRMs (top), and the corresponding RNA splicing maps (right). Differential gene expression (DEG) values of corresponding RBP upon knockdown are shown as triangles. The upper and lower vertices indicated up- and down-regulation, respectively, and filled triangles denoted differential expression changes (*i.e.*, |log2FC| ≥0.1 and adjusted p-values ≤0.1). Barplots indicate the mean AS per the candidate regulator.

**Figure S21. RNAMaRs results for UCHL5-regulated exons in HepG2 cells.**

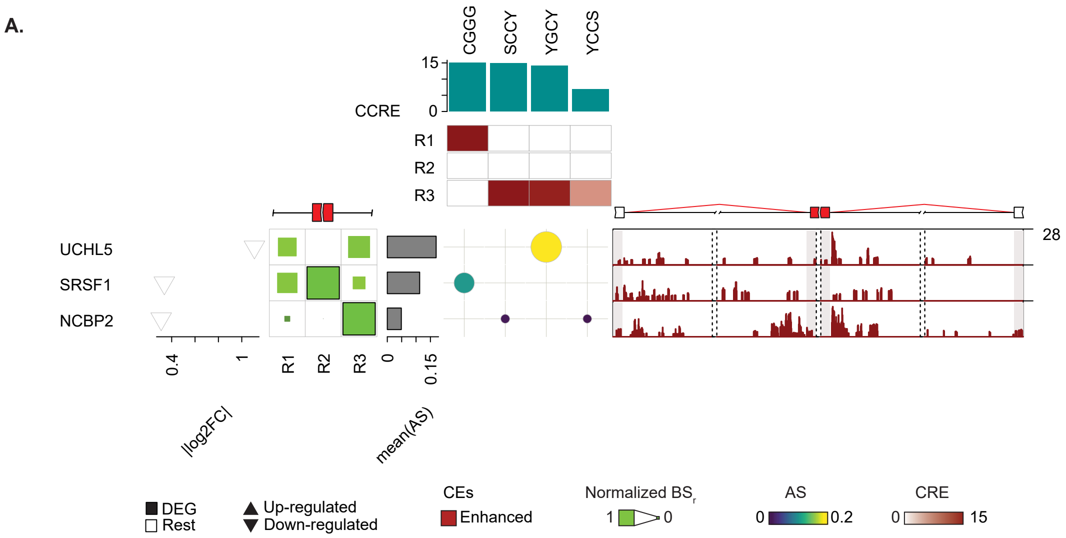


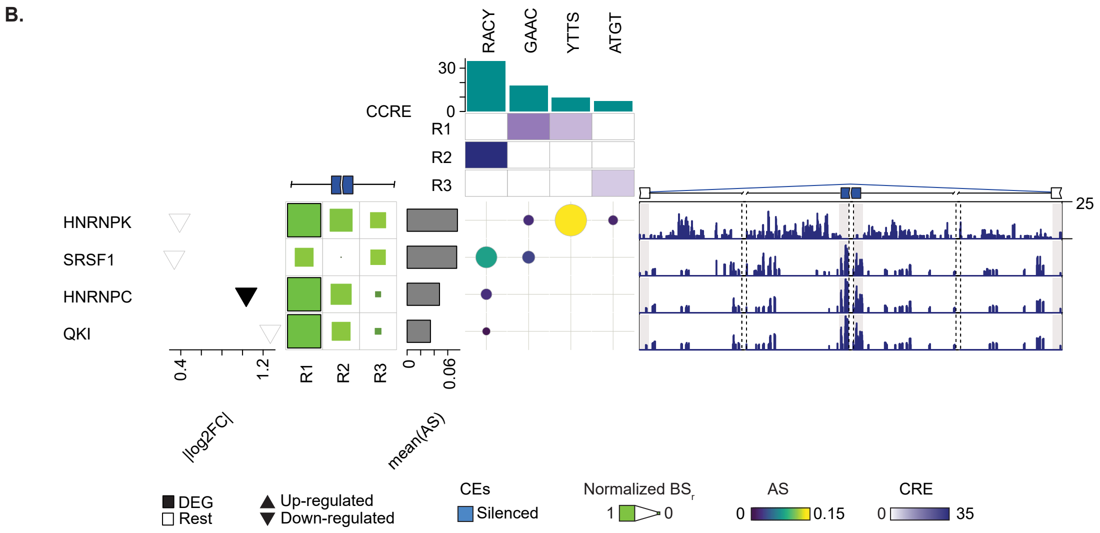


Complete RNAMaRs outputs for UCHL5-regulated enhanced **(A)** and silenced **(B)** exons in HepG2 cells. The visualization integrates normalized binding scores (BS) across splice-proximal regions (R1-R3) (left), MRM-RBP association scores (AS), cumulative combined regional enrichment (CCRE) across enriched MRMs (top), and the corresponding RNA splicing maps (right). Differential gene expression (DEG) values of corresponding RBP upon knockdown are shown as triangles. The upper and lower vertices indicated up- and down-regulation, respectively, and filled triangles denoted differential expression changes (*i.e.*, |log2FC| ≥0.1 and adjusted p-values ≤0.1). Barplots indicate the mean AS per the candidate regulator.

**Figure S22. RNAMaRs results for AGGF1-regulated exons in K562 cells.**

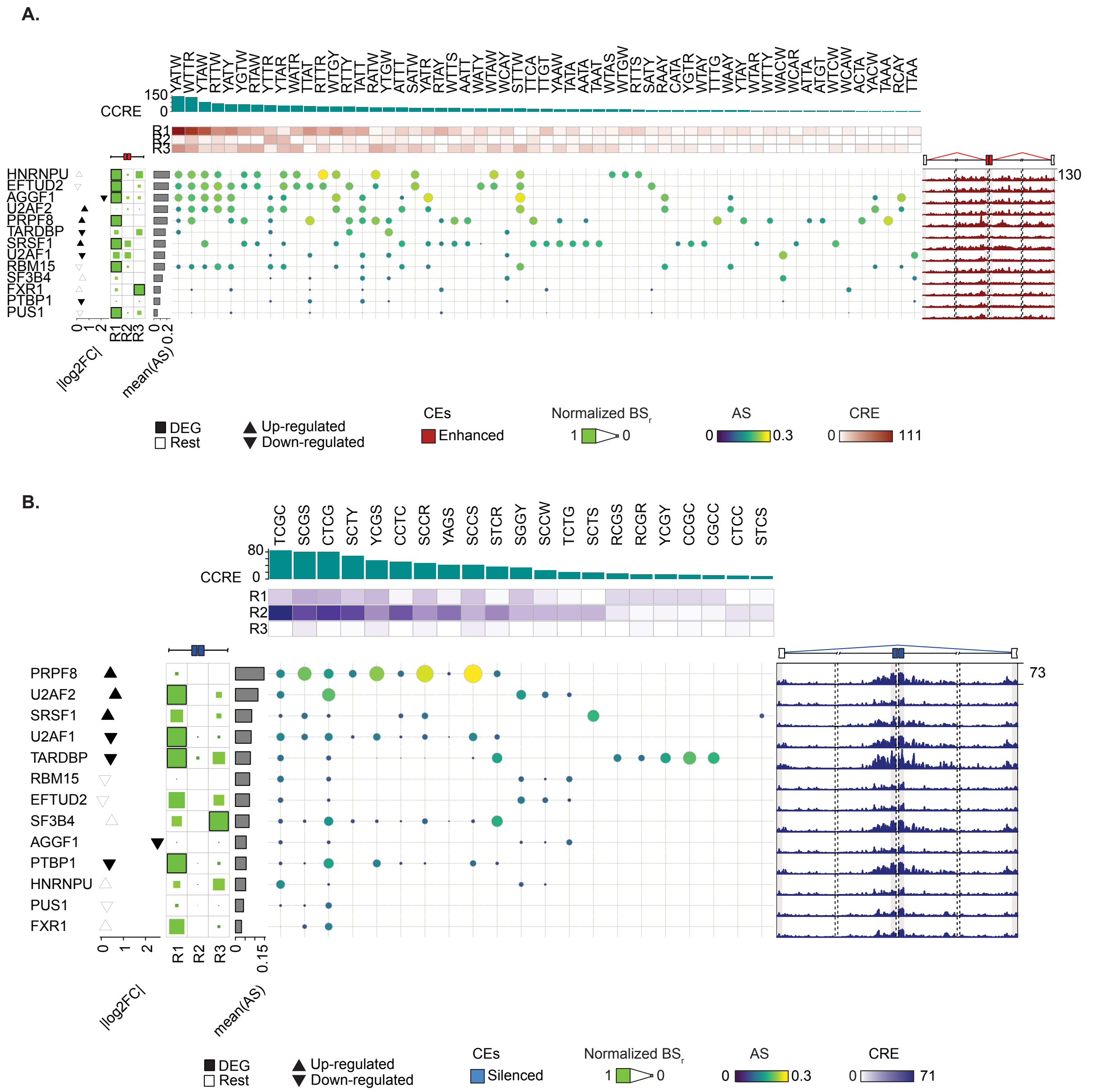


Complete RNAMaRs outputs for AGGF1-regulated enhanced **(A)** and silenced **(B)** exons in K562 cells. The visualization integrates normalized binding scores (BS) across splice-proximal regions (R1-R3) (left), MRM-RBP association scores (AS), cumulative combined regional enrichment (CCRE) across enriched MRMs (top), and the corresponding RNA splicing maps (right). Differential gene expression (DEG) values of corresponding RBP upon knockdown are shown as triangles. The upper and lower vertices indicated up- and down-regulation, respectively, and filled triangles denoted differential expression changes (*i.e.*, |log2FC| ≥0.1 and adjusted p-values ≤0.1). Barplots indicate the mean AS per the candidate regulator.

**Figure S23. RNAMaRs results for EFTUD2-regulated exons in K562 cells.**

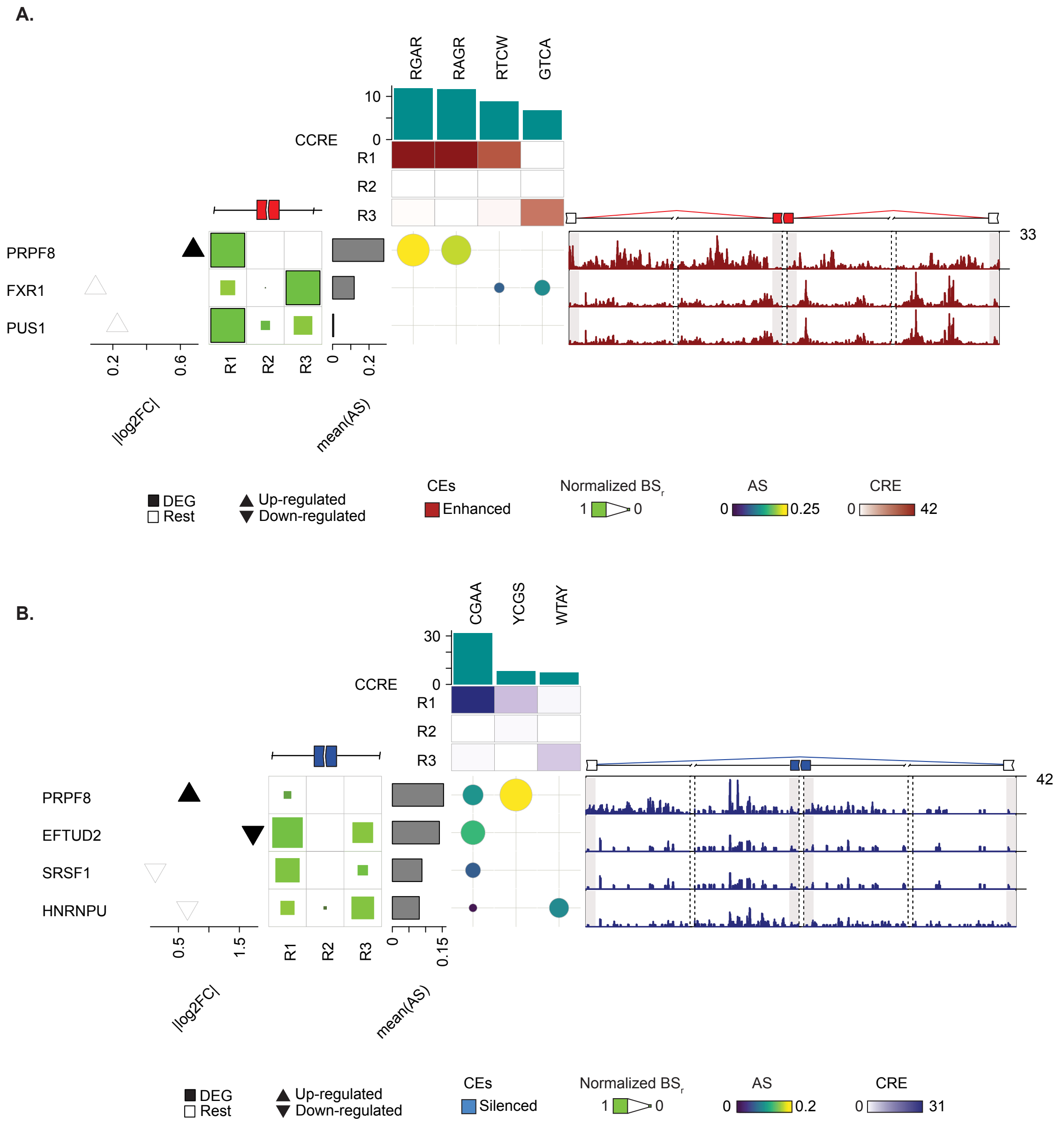


Complete RNAMaRs outputs for EFTUD2-regulated enhanced **(A)** and silenced **(B)** exons in K562 cells. The visualization integrates normalized binding scores (BS) across splice-proximal regions (R1-R3) (left), MRM-RBP association scores (AS), cumulative combined regional enrichment (CCRE) across enriched MRMs (top), and the corresponding RNA splicing maps (right). Differential gene expression (DEG) values of corresponding RBP upon knockdown are shown as triangles. The upper and lower vertices indicated up- and down-regulation, respectively, and filled triangles denoted differential expression changes (*i.e.*, |log2FC| ≥0.1 and adjusted p-values ≤0.1). Barplots indicate the mean AS per the candidate regulator.

**Figure S24. RNAMaRs results for FXR1-regulated exons in K562 cells.**

A.

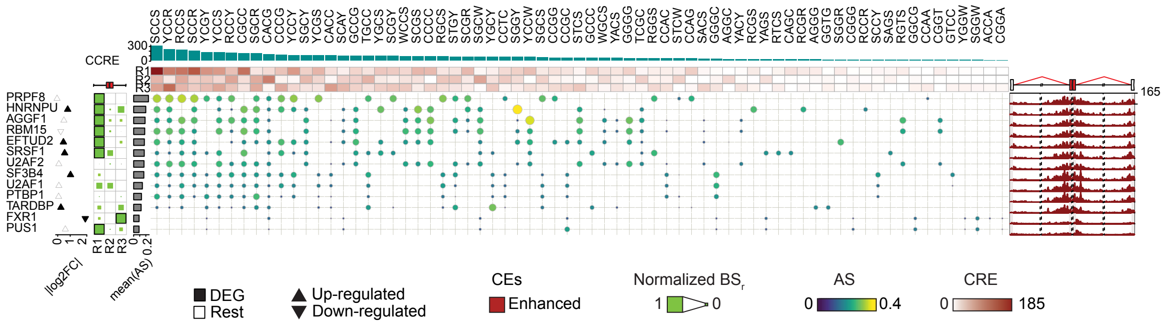


**A.** Complete RNAMaRs outputs for FXR1-regulated enhanced exons in K562 cells. The visualization integrates normalized binding scores (BS) across splice-proximal regions (R1-R3) (left), MRM-RBP association scores (AS), cumulative combined regional enrichment (CCRE) across enriched MRMs (top), and the corresponding RNA splicing maps (right). Differential gene expression (DEG) values of corresponding RBP upon knockdown are shown as triangles. The upper and lower vertices indicated up- and down-regulation, respectively, and filled triangles denoted differential expression changes (*i.e.*, |log2FC| ≥0.1 and adjusted p-values ≤0.1). Barplots indicate the mean AS per the candidate regulator.

**Figure S25. RNAMaRs results for HNRNPU-regulated exons in K562 cells.**

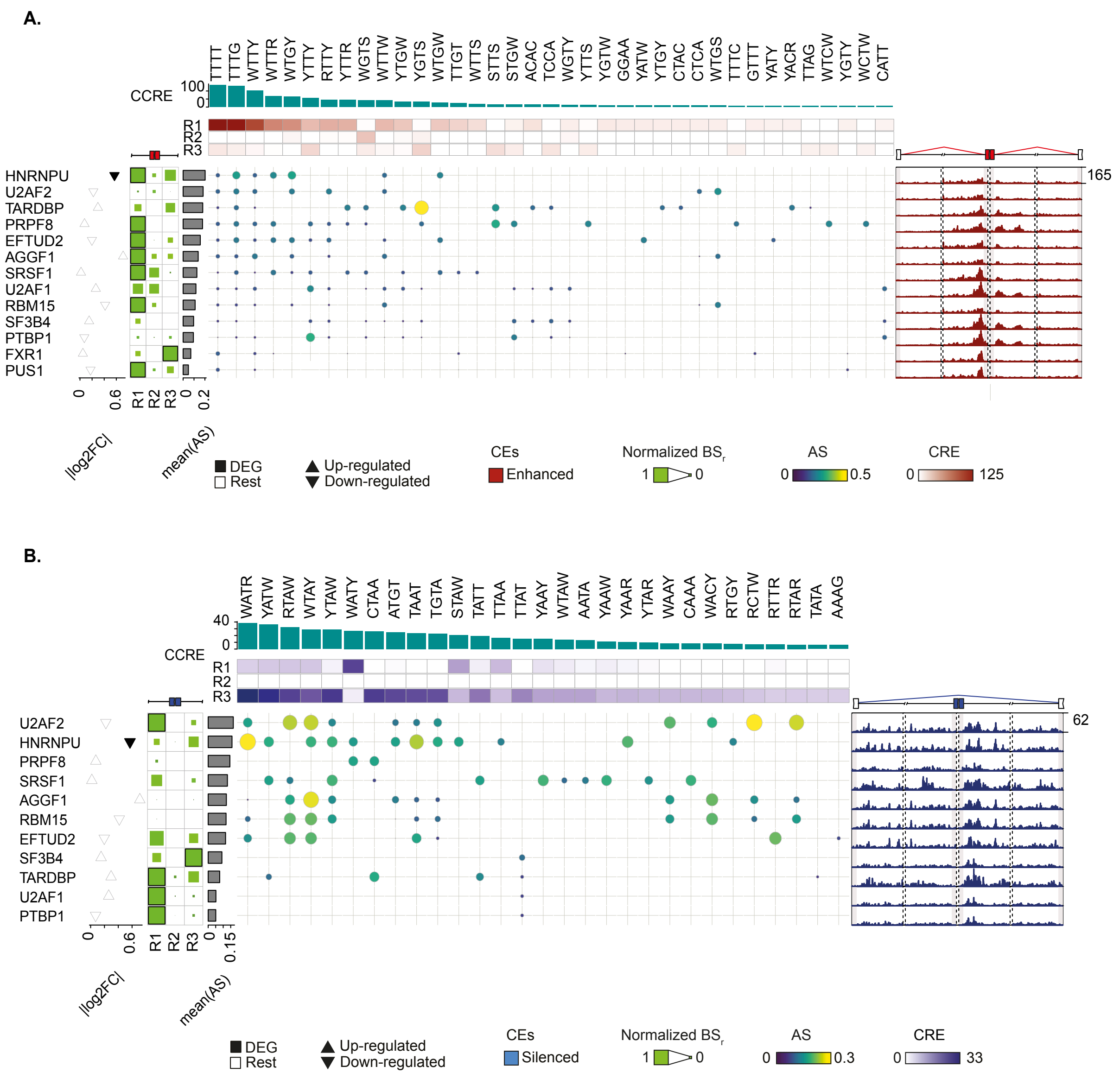


Complete RNAMaRs outputs for HNRNPU-regulated enhanced **(A)** and silenced **(B)** exons in K562 cells. The visualization integrates normalized binding scores (BS) across splice-proximal regions (R1-R3) (left), MRM-RBP association scores (AS), cumulative combined regional enrichment (CCRE) across enriched MRMs (top), and the corresponding RNA splicing maps (right). Differential gene expression (DEG) values of corresponding RBP upon knockdown are shown as triangles. The upper and lower vertices indicated up- and down-regulation, respectively, and filled triangles denoted differential expression changes (*i.e.*, |log2FC| ≥0.1 and adjusted p-values ≤0.1). Barplots indicate the mean AS per the candidate regulator.

**Figure S26. RNAMaRs results for PRPF8-regulated exons in K562 cells.**

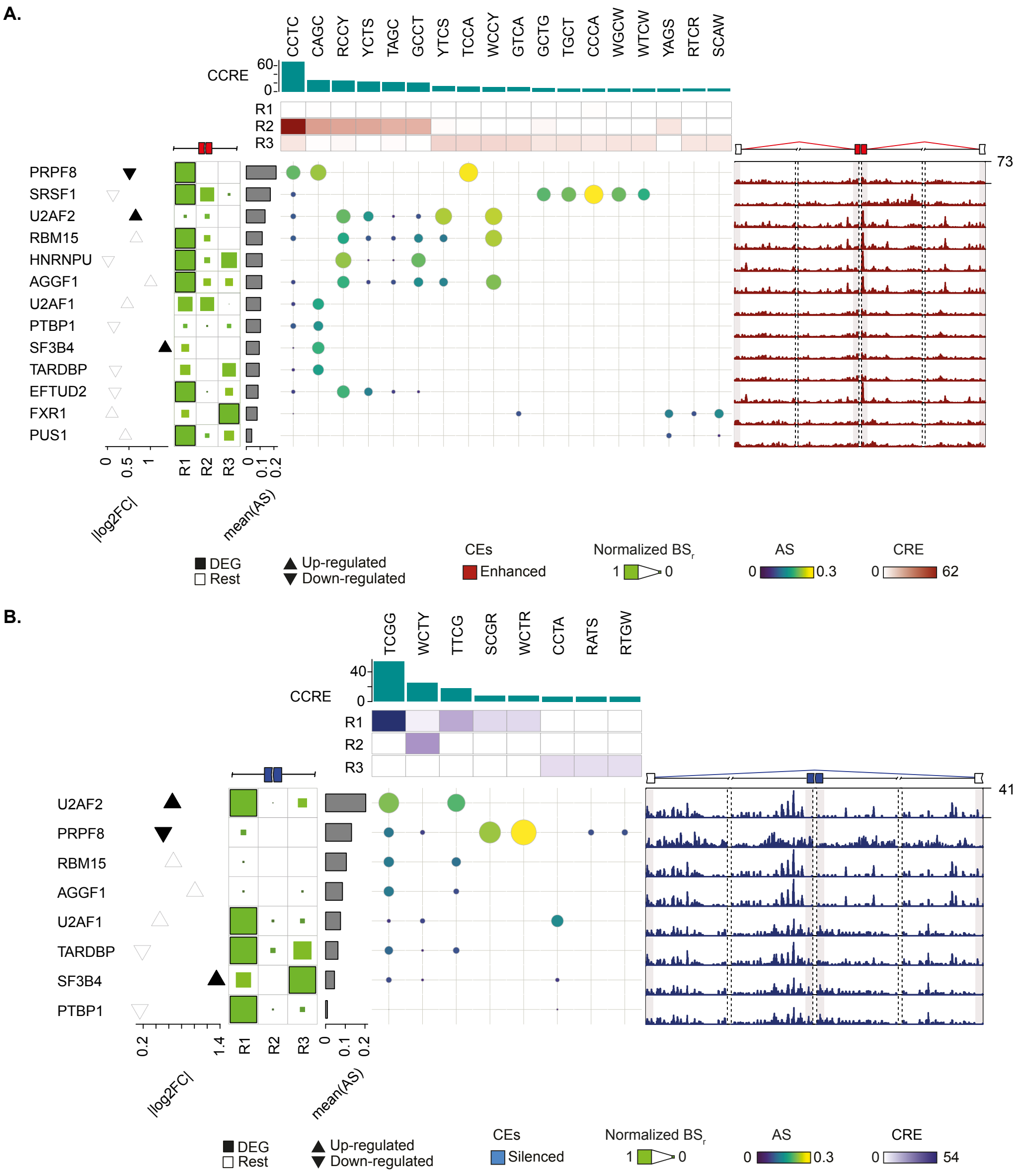


Complete RNAMaRs outputs for PRPF8-regulated enhanced **(A)** and silenced **(B)** exons in K562 cells. The visualization integrates normalized binding scores (BS) across splice-proximal regions (R1-R3) (left), MRM-RBP association scores (AS), cumulative combined regional enrichment (CCRE) across enriched MRMs (top), and the corresponding RNA splicing maps (right). Differential gene expression (DEG) values of corresponding RBP upon knockdown are shown as triangles. The upper and lower vertices indicated up- and down-regulation, respectively, and filled triangles denoted differential expression changes (*i.e.*, |log2FC| ≥0.1 and adjusted p-values ≤0.1). Barplots indicate the mean AS per the candidate regulator.

**Figure S27. RNAMaRs results for PTBP1-regulated exons in K562 cells.**

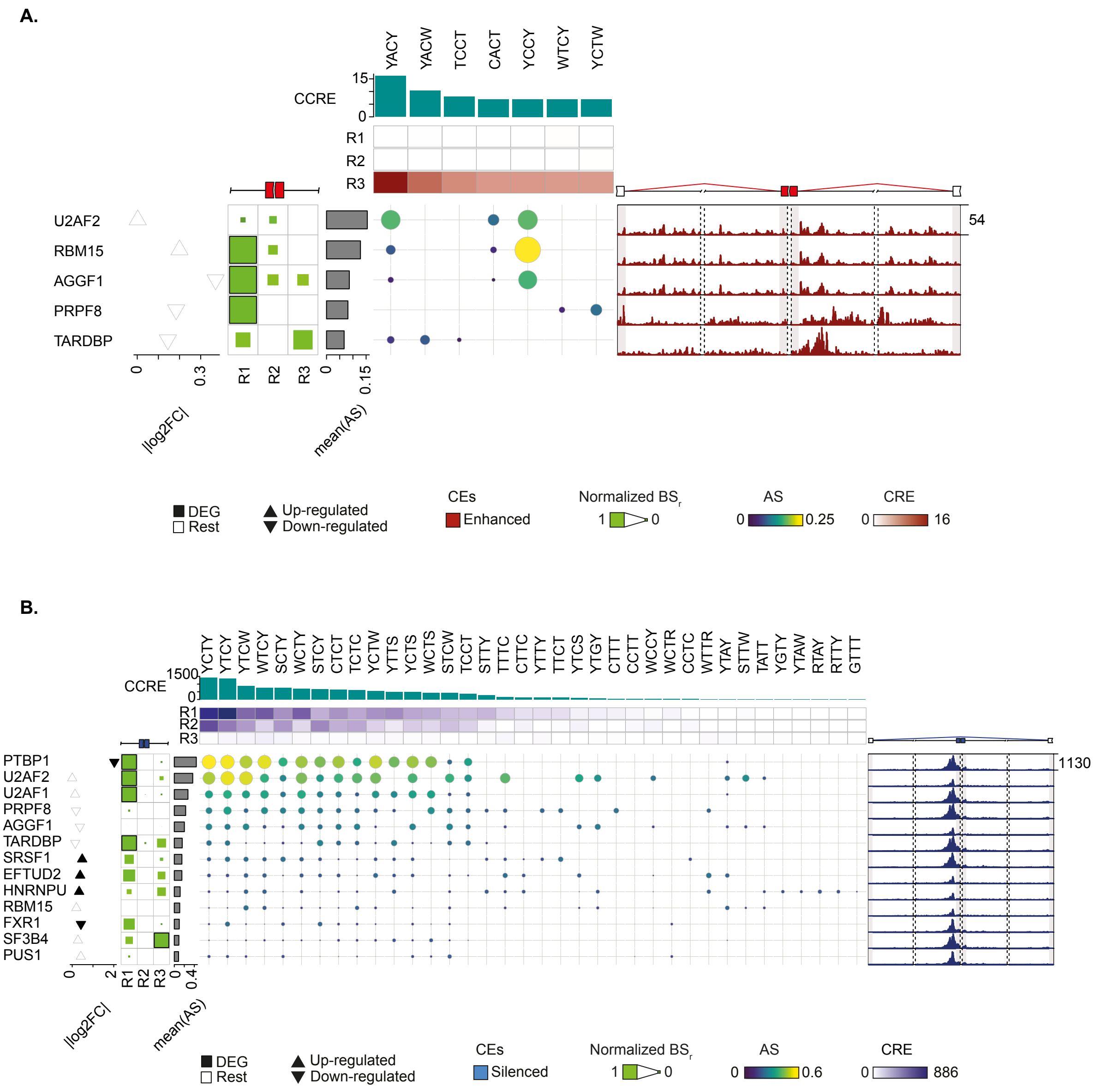


Complete RNAMaRs outputs for PTBP1-regulated enhanced **(A)** and silenced **(B)** exons in K562 cells. The visualization integrates normalized binding scores (BS) across splice-proximal regions (R1-R3) (left), MRM-RBP association scores (AS), cumulative combined regional enrichment (CCRE) across enriched MRMs (top), and the corresponding RNA splicing maps (right). Differential gene expression (DEG) values of corresponding RBP upon knockdown are shown as triangles. The upper and lower vertices indicated up- and down-regulation, respectively, and filled triangles denoted differential expression changes (*i.e.*, |log2FC| ≥0.1 and adjusted p-values ≤0.1). Barplots indicate the mean AS per the candidate regulator.

**Figure S28. RNAMaRs results for PUS1-regulated exons in K562 cells.**

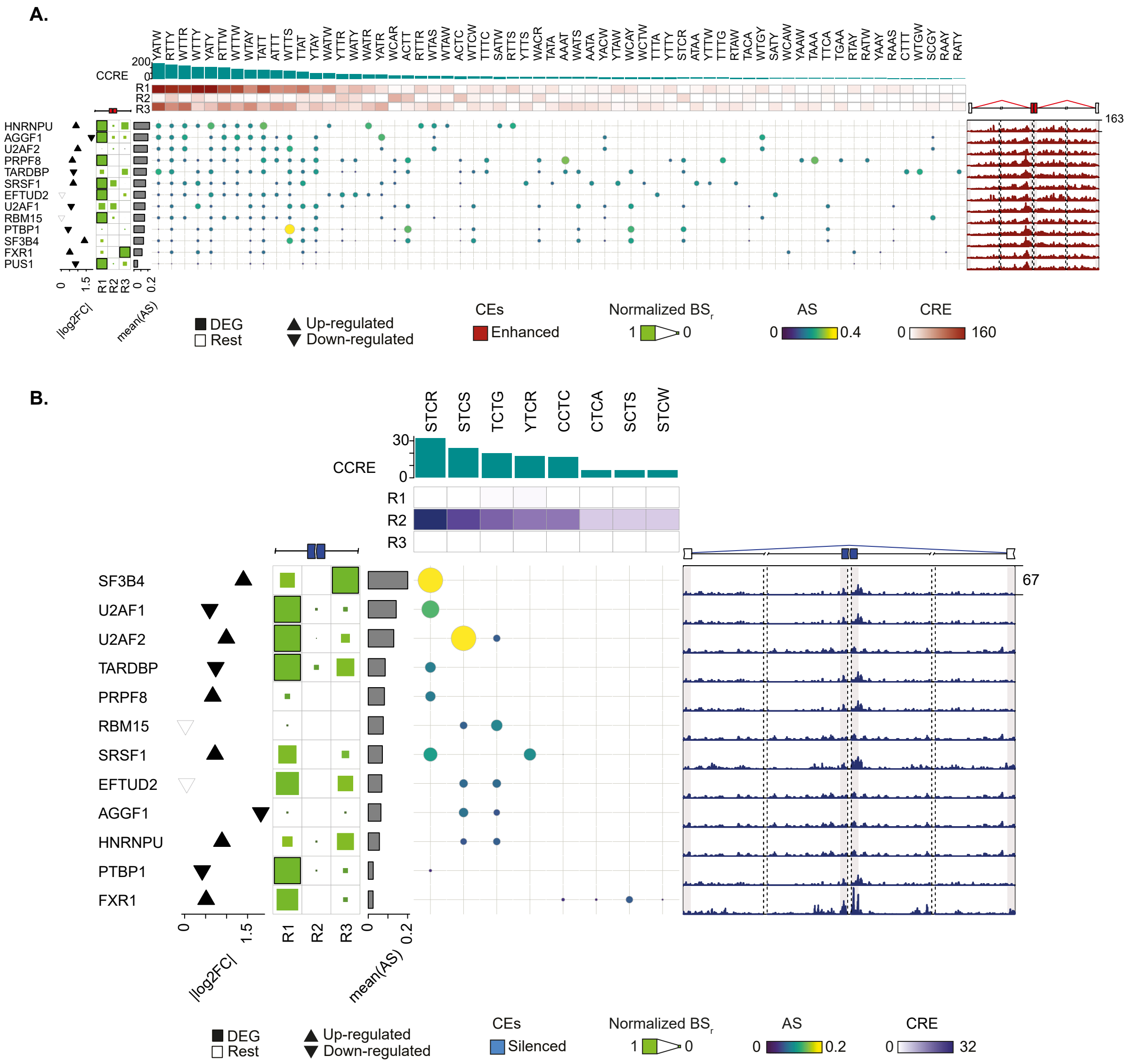


Complete RNAMaRs outputs for PUS1-regulated enhanced **(A)** and silenced **(B)** exons in K562 cells. The visualization integrates normalized binding scores (BS) across splice-proximal regions (R1-R3) (left), MRM-RBP association scores (AS), cumulative combined regional enrichment (CCRE) across enriched MRMs (top), and the corresponding RNA splicing maps (right). Differential gene expression (DEG) values of corresponding RBP upon knockdown are shown as triangles. The upper and lower vertices indicated up- and down-regulation, respectively, and filled triangles denoted differential expression changes (*i.e.*, |log2FC| ≥0.1 and adjusted p-values ≤0.1). Barplots indicate the mean AS per the candidate regulator.

**Figure S29. RNAMaRs results for RBM15-regulated exons in K562 cells.**

A.

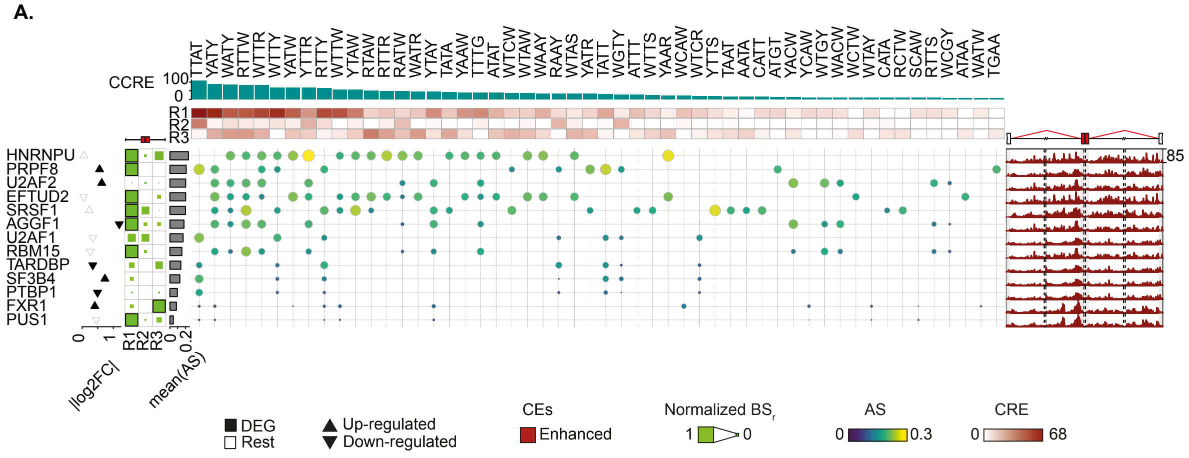


**A.** Complete RNAMaRs outputs for RBM15-regulated enhanced exons in K562 cells. The visualization integrates normalized binding scores (BS) across splice-proximal regions (R1-R3) (left), MRM-RBP association scores (AS), cumulative combined regional enrichment (CCRE) across enriched MRMs (top), and the corresponding RNA splicing maps (right). Differential gene expression (DEG) values of corresponding RBP upon knockdown are shown as triangles. The upper and lower vertices indicated up- and down-regulation, respectively, and filled triangles denoted differential expression changes (*i.e.*, |log2FC| ≥0.1 and adjusted p-values ≤0.1). Barplots indicate the mean AS per the candidate regulator.

**Figure S30. RNAMaRs results for SF3B4-regulated exons in K562 cells.**

A.

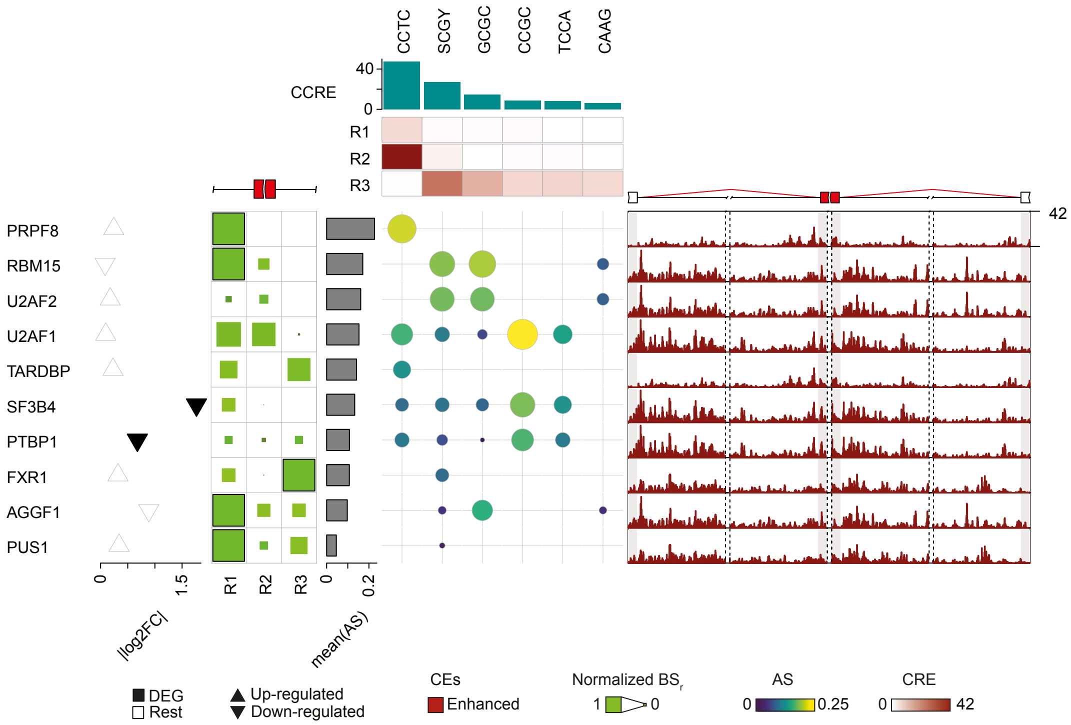


B.

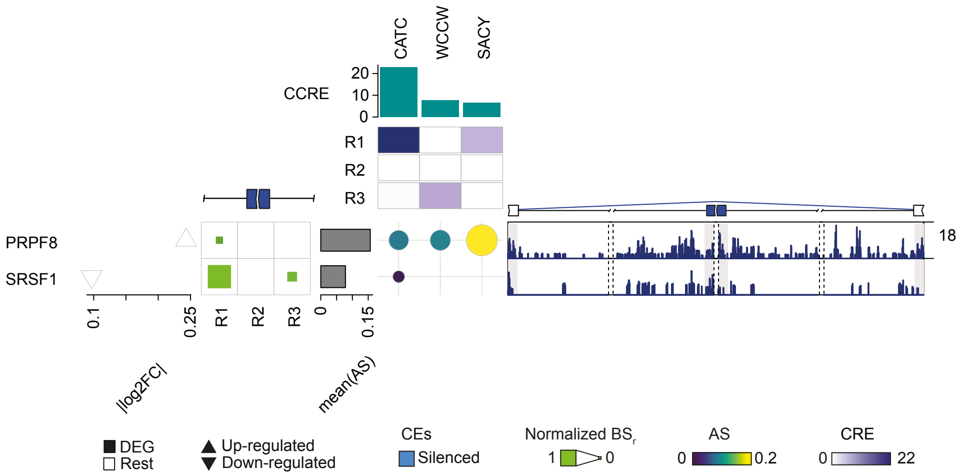


Complete RNAMaRs outputs for SF3B4-regulated enhanced **(A)** and silenced **(B)** exons in K562 cells. The visualization integrates normalized binding scores (BS) across splice-proximal regions (R1-R3) (left), MRM-RBP association scores (AS), cumulative combined regional enrichment (CCRE) across enriched MRMs (top), and the corresponding RNA splicing maps (right). Differential gene expression (DEG) values of corresponding RBP upon knockdown are shown as triangles. The upper and lower vertices indicated up- and down-regulation, respectively, and filled triangles denoted differential expression changes (*i.e.*, |log2FC| ≥0.1 and adjusted p-values ≤0.1). Barplots indicate the mean AS per the candidate regulator.

**Figure S31. RNAMaRs results for SRSF1-regulated exons in K562 cells.**

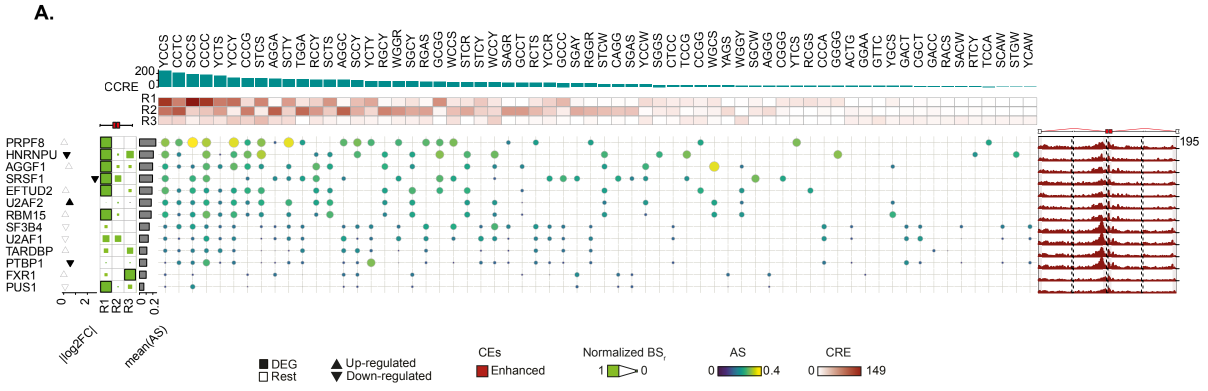


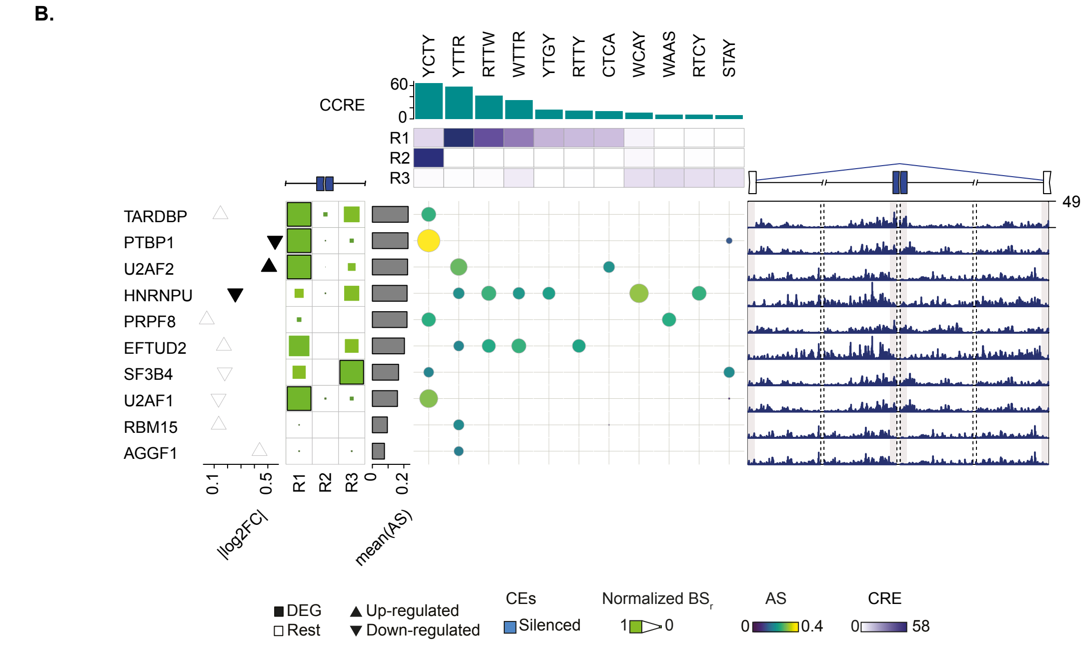


Complete RNAMaRs outputs for SRSF1-regulated enhanced **(A)** and silenced **(B)** exons in K562 cells. The visualization integrates normalized binding scores (BS) across splice-proximal regions (R1-R3) (left), MRM-RBP association scores (AS), cumulative combined regional enrichment (CCRE) across enriched MRMs (top), and the corresponding RNA splicing maps (right). Differential gene expression (DEG) values of corresponding RBP upon knockdown are shown as triangles. The upper and lower vertices indicated up- and down-regulation, respectively, and filled triangles denoted differential expression changes (*i.e.*, |log2FC| ≥0.1 and adjusted p-values ≤0.1). Barplots indicate the mean AS per the candidate regulator.

**Figure S32. RNAMaRs results for TARDBP-regulated exons in K562 cells.**

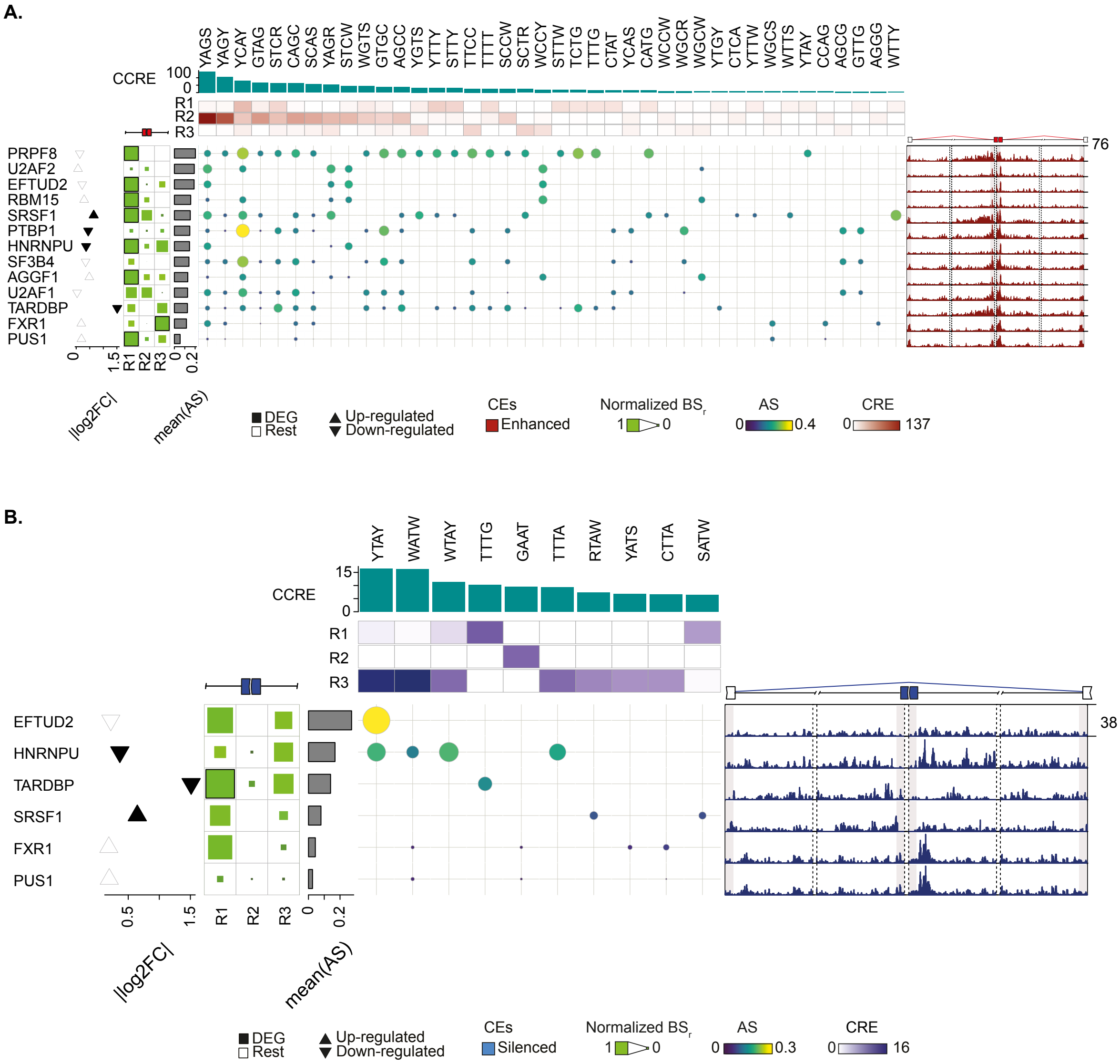


Complete RNAMaRs outputs for TARDBP-regulated enhanced **(A)** and silenced **(B)** exons in K562 cells. The visualization integrates normalized binding scores (BS) across splice-proximal regions (R1-R3) (left), MRM-RBP association scores (AS), cumulative combined regional enrichment (CCRE) across enriched MRMs (top), and the corresponding RNA splicing maps (right). Differential gene expression (DEG) values of corresponding RBP upon knockdown are shown as triangles. The upper and lower vertices indicated up- and down-regulation, respectively, and filled triangles denoted differential expression changes (*i.e.*, |log2FC| ≥0.1 and adjusted p-values ≤0.1). Barplots indicate the mean AS per the candidate regulator.

**Figure S33. RNAMaRs results for U2AF1-regulated exons in K562 cells.**

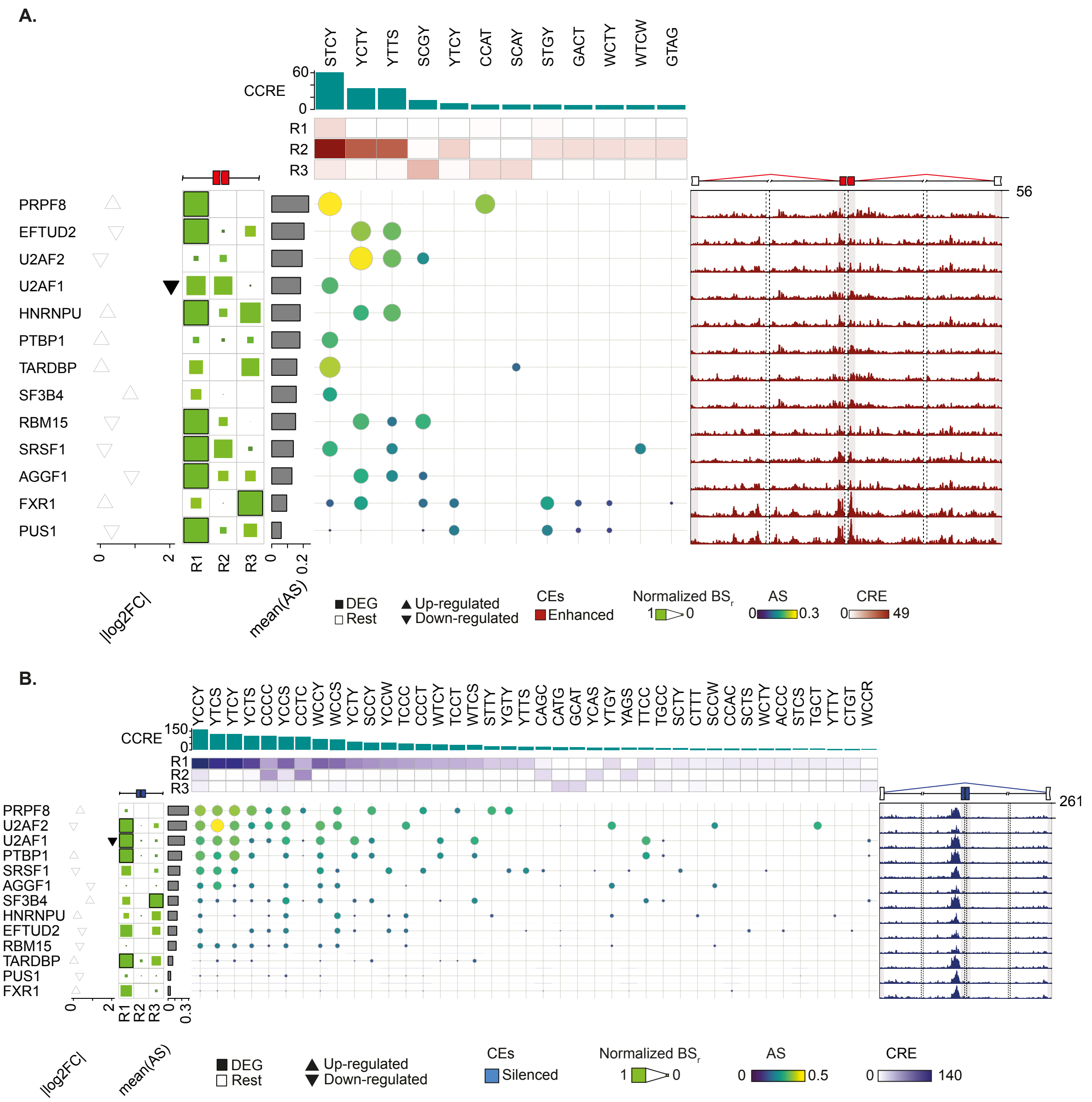


Complete RNAMaRs outputs for U2AF1-regulated enhanced **(A)** and silenced **(B)** exons in K562 cells. The visualization integrates normalized binding scores (BS) across splice-proximal regions (R1-R3) (left), MRM-RBP association scores (AS), cumulative combined regional enrichment (CCRE) across enriched MRMs (top), and the corresponding RNA splicing maps (right). Differential gene expression (DEG) values of corresponding RBP upon knockdown are shown as triangles. The upper and lower vertices indicated up- and down-regulation, respectively, and filled triangles denoted differential expression changes (*i.e.*, |log2FC| ≥0.1 and adjusted p-values ≤0.1). Barplots indicate the mean AS per the candidate regulator.

**Figure S34. RNAMaRs results for U2AF2-regulated exons in K562 cells.**

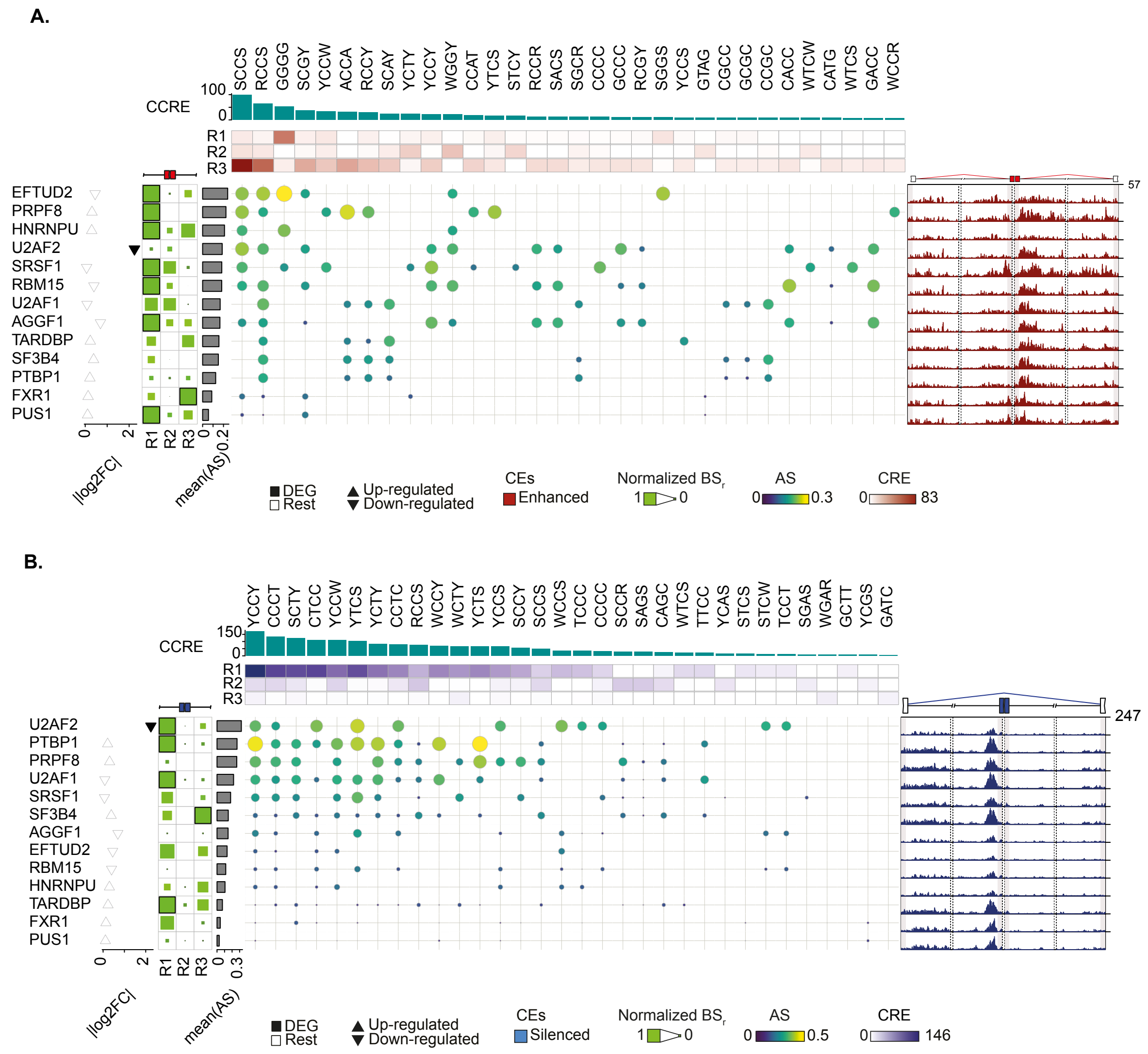


Complete RNAMaRs outputs for U2AF2-regulated enhanced **(A)** and silenced **(B)** exons in K562 cells. The visualization integrates normalized binding scores (BS) across splice-proximal regions (R1-R3) (left), MRM-RBP association scores (AS), cumulative combined regional enrichment (CCRE) across enriched MRMs (top), and the corresponding RNA splicing maps (right). Differential gene expression (DEG) values of corresponding RBP upon knockdown are shown as triangles. The upper and lower vertices indicated up- and down-regulation, respectively, and filled triangles denoted differential expression changes (*i.e.*, |log2FC| ≥0.1 and adjusted p-values ≤0.1). Barplots indicate the mean AS per the candidate regulator.
